# Molecular logic for cellular specializations that initiate the auditory parallel processing pathways

**DOI:** 10.1101/2023.05.15.539065

**Authors:** Junzhan Jing, Ming Hu, Tenzin Ngodup, Qianqian Ma, Shu-Ning Natalie Lau, Cecilia Ljungberg, Matthew J. McGinley, Laurence O. Trussell, Xiaolong Jiang

## Abstract

The cochlear nuclear complex (CN), the starting point for all central auditory processing, comprises a suite of neuronal cell types that are highly specialized for neural coding of acoustic signals, yet molecular logic governing cellular specializations remains unknown. By combining single-nucleus RNA sequencing and Patch-seq analysis, we reveal a set of transcriptionally distinct cell populations encompassing all previously observed types and discover multiple new subtypes with anatomical and physiological identity. The resulting comprehensive cell-type taxonomy reconciles anatomical position, morphological, physiological, and molecular criteria, enabling the determination of the molecular basis of the remarkable cellular phenotypes in the CN. In particular, CN cell-type identity is encoded in a transcriptional architecture that orchestrates functionally congruent expression across a small set of gene families to customize projection patterns, input-output synaptic communication, and biophysical features required for encoding distinct aspects of acoustic signals. This high-resolution account of cellular heterogeneity from the molecular to the circuit level illustrates molecular logic for cellular specializations and enables genetic dissection of auditory processing and hearing disorders with unprecedented specificity.

## Introduction

Information processing along parallel pathways is an important feature of most biological sensory systems. In the mammalian auditory system, auditory signals are encoded by cochlear hair cells and transmitted to the first processing station in the brain, the cochlear nuclear complex (CN), where all higher-level parallel pathways are initiated^1^. Neurons in the CN are organized tonotopically into precise parallel sensory maps and each is highly specialized for processing different aspects of acoustic information including sound location, intensity, frequency content, and spectrotemporal modulations^2,3^. Classical studies, using histology, in-vivo and in-vitro recording, and pathway tracing, have classified CN projection neurons into at least four types: the bushy, T-stellate, fusiform, and octopus cells, which each carry distinct ascending signals needed for high-level sound processing^4–6^. In addition, multiple types of interneurons within CN determine the pattern and magnitude of signals that ascend through the neuraxis^6^. These cell types in CN have been extensively characterized and clearly defined based on their unique morpho-physiological properties, synaptic connectivity, projection targets, and functions^4–6^. We term these well-defined projection neurons and interneurons as “established cell types”. Despite their remarkable phenotypic differences, molecular distinctions and mechanisms that specify and distinguish these cell types are still largely unknown. Since most of these cell types remain molecularly inaccessible, the exact contribution of any given cell type in CN to sound representations in higher brain regions is still unknown. In addition, numerous lines of evidence suggest that the established cell types do not capture the true range of cellular heterogeneity and their functions. For example, in vivo recordings document diverse patterns of acoustic response within a given type of neurons, suggesting that existing definitions of cell types do not have sufficient resolution to account for the variety of downstream anatomical projections and physiological responses^7–9^. In addition, recent in vitro physiological studies using Cre mouse lines and optogenetics have suggested the existence of previously undescribed cell types in CN^10,11^. Therefore, a full spectrum of cell types and their functions in CN have yet to be established, preventing a full understanding of central auditory processing.

Single-cell transcriptomics offers a powerful, unbiased approach to profile cellular and molecular heterogeneity and define cell types through gene expression profiling of individual cells^12–14^. To capture the full spectrum of cellular heterogeneity and their functions in CN, we employed single-nucleus RNA sequencing (snRNA-seq) to profile the transcriptome of the whole CN at the single-cell level and generate a comprehensive taxonomy of transcriptomic cell types. To relate each transcriptionally distinct cell population to the established CN cell types with functional relevance, we performed parallel Patch-seq experiments that link each cell’s transcriptome to their morphological and physiological features^15^, followed by the validation using fluorescent in situ hybridization (FISH) and transgenic mouse lines. These approaches allow us to establish a clear correspondence between transcriptionally distinct cell types and all previously established CN cell types. Importantly, we also discover new subtypes based on transcriptomic profiling that reflect hitherto unresolved distinctions in anatomy, morphology, and physiology of cell types^16,17^, suggesting novel auditory processing pathways. Our analyses of single-cell transcriptomes identified novel molecular markers for CN cell types and illustrated potential molecular and gene regulatory mechanisms that specify cell type identity, connectivity and projection patterns, and remarkable synaptic and biophysical features, providing molecular delineations of parallel auditory pathways at their initiation.

## Results

### Identification of transcriptionally distinct cell populations in mouse CN

To investigate transcriptional and cellular heterogeneity within the CN, we performed RNA sequencing on individual nuclei isolated from CN of male and female C57BL/6 mice (postnatal day (P) 22-28) (Extended Data Fig. 1a). Our final dataset contained 61,211 CN nuclei (Methods and Extended Data Fig. 2-3), and analysis of sequence reads revealed ∼4,000 uniquely detected transcripts from ∼1,900 genes for each nucleus. Following data integration, unsupervised dimensionality reduction techniques identified 14 major transcriptionally distinct cell populations (Extended Data Fig. 1b). By inspection of known markers, we identified all non-neuronal cell types in CN (Extended Data Fig. 1c and Extended Data Table 1). Using the pan-neuronal marker *Snap25*, we identified 31,639 neuronal nuclei expressing either the gene for vesicular glutamate transporters *Slc17a7* or *Slc17a6* or the gene for the glycine transporter *Slc6a5*, thereby separating glutamatergic and glycinergic populations (Extended Data Fig. 1b-c). Clustering analysis of neuronal nuclei resulted in 13 molecularly distinct populations (Fig. 1b) including seven glutamatergic/excitatory clusters (*Slc17a7*^+^ and/or *Slc17a6*^+^) and six glycinergic/inhibitory clusters (*Slc6a5^+^*, Fig. 1c).

**Fig. 1:**
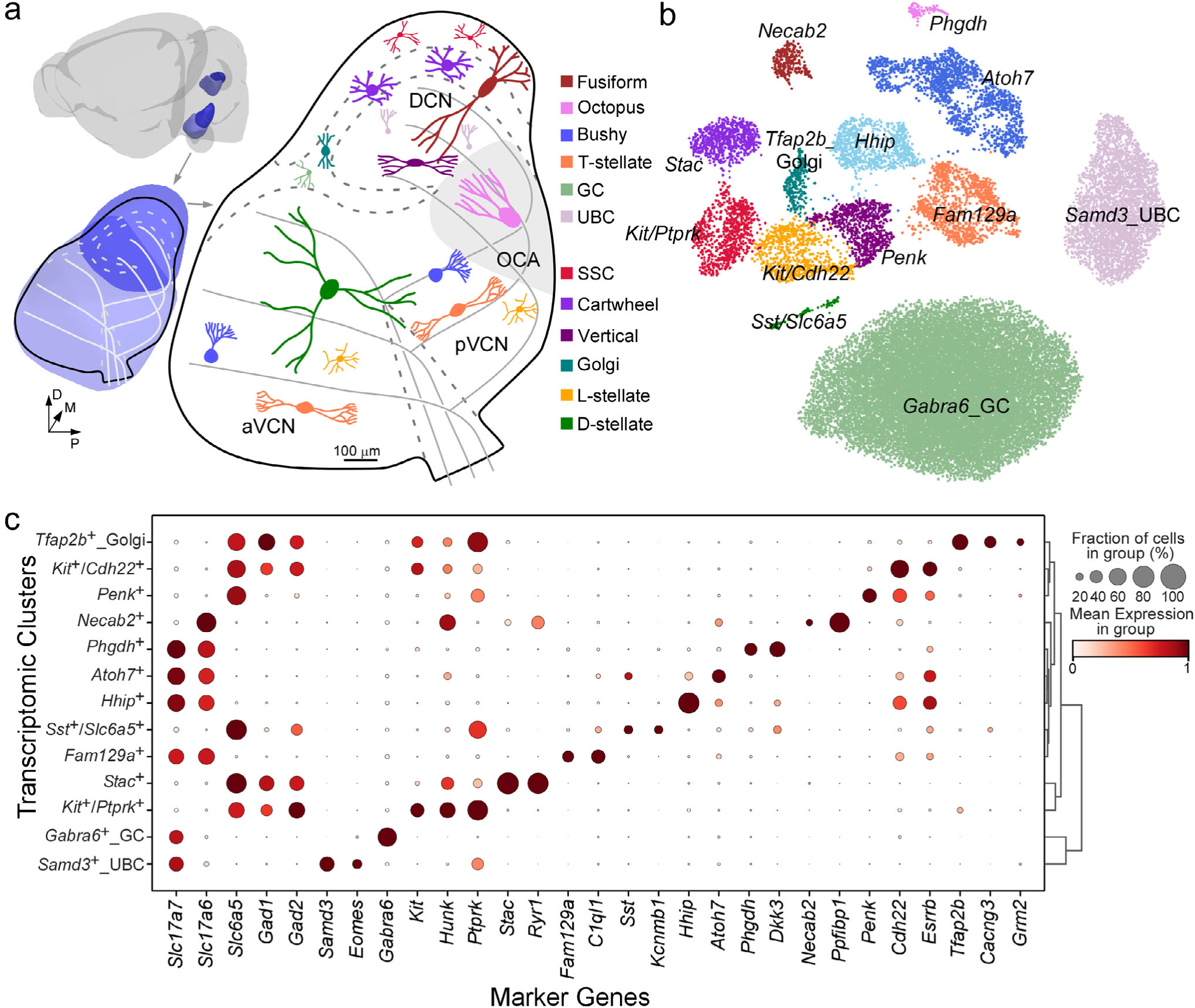
Comprehensive transcriptional profiling of cell types across the mouse CN. **a**, Anatomy of mouse cochlear nucleus (CN; Light blue: VCN. Dark blue: DCN) and depiction of neuronal cell types (cartoon drawings) across the CN (sagittal view), colored by cell type identity. D: dorsal, P: posterior; M: medial. **b**, UMAP visualization of 31,639 nuclei from CN neurons clustered by over-dispersed genes, colored by cluster identity. Each cluster labeled by key marker genes. **c**, Right: dendrogram indicating hierarchical relationships between clusters. Left: dot plot of scaled expression of selected marker genes for each cluster shown in (**b**); see Extended Data Table 1. Circle size depicts percentage of cells in the cluster in which the marker was detected (≥ 1 UMI), and color depicts average transcript count in expressing cells. See Extended Data Fig. 1-3 for more technical details and annotations.

Based on known markers, we assigned three clusters to known CN cell types. Two glutamatergic clusters that expressed only *Slc17a7* (not *Slc17a6*) were either enriched with *Gabra6*, a marker gene for granule cells (GCs)^18,19^, or with *Tbr2*/*Eomes* and *Samd3,* gene markers for unipolar brush cells (UBCs)^20,21^, thus corresponding to GCs and UBCs in CN, respectively (Fig. 1b-c, Extended Data Fig. 1d). The UBC cluster was further validated with RNA fluorescent in situ hybridization (FISH) staining for *Samd3* (Extended Data Fig. 1d) and characterized by graded and inversely correlated expression of glutamate excitatory signaling pathways and inhibitory pathways, reminiscent of UBCs in cerebellum (Extended Data Fig. 1e)^21–23^. One glycinergic cluster was enriched with *Grm2* (Fig. 1c), a marker gene for Golgi cells in CN (in addition to UBCs)^24,25^, thus corresponding to auditory Golgi cells (Fig. 1b-c). Differentially expressed gene (DEG) analysis identified *Cacng3* and *Tfap2b* as more specific marker genes for auditory Golgi cells (Fig. 1b-c), which were validated with FISH (Extended Data Fig. 1f). Other major clusters are unknown in terms of their potential correspondence to well-studied CN cell types (Fig. 1a) but could be labeled by one or two candidate gene markers identified via DEG analysis (Fig. 1b-c and Extended Data Table 1).

### Molecular profiles of CN excitatory neurons

To relate the unknown clusters to known CN cell types, we performed Patch-seq, starting with four major excitatory projection neurons, bushy, T-stellate, octopus, and fusiform cells^4^, targeting each by their known anatomical location aided by transgenic mouse lines (see Methods). Membrane potential responses to current injections were recorded, cytosol and nucleus mRNA were extracted and reverse transcribed, and the resulting cDNA was sequenced (Extended Data Fig. 3g-k, Methods). The morphology of each neuron was *post hoc* recovered and reconstructed. Based on their stereotypical morphological and electrophysiological properties, each neuron could be readily identified and assigned to a type, a process we term “expert classification” (Methods; Fig. 1a, 2a-b, Extended Data Table 2)^26^. To validate expert classification and support cell-type assignment of each Patch-seq neuron, we used a random forest classifier (Methods; Fig. 2c-d, Extended Data Fig. 4a) or clustering-based k-means classifier (Methods; Extended Data Fig. 4b) based on either electrophysiology or morphology to distinguish between any two types of CN excitatory neurons as labeled manually, resulting in successful separation of almost all cell type pairs, thereby supporting our expert classification. UMAP projection of these Patch-seq cells resulted in five transcriptomic clusters, and cells labeled as the same type clustered together and were rarely confused with another cluster (Fig. 2e), except for bushy cells, which fell into two prominent clusters. Using the gene expression profiles, we mapped these Patch-seq neurons to the transcriptomic clusters identified via snRNA-seq (Fig. 1b)^27,28^, demonstrating close correspondence in the two datasets and identifying transcriptomic clusters for the established excitatory cell types (Methods, Fig 2f-g). Importantly, those Patch-seq bushy cells clustering together as Cluster 1 in Patch-seq UMAP space (Fig. 2e) were all mapped to the *Hhip*^+^ cluster (Fig. 2f), while almost all bushy cells in Cluster 2 (Fig. 2e) were mapped to the *Atoh7*^+^ cluster (Fig. 2f). Thus, our analysis not only reveals a strong congruence across three modalities in excitatory neuron classification, but also molecular heterogeneity among bushy cells (see below).

**Fig. 2:**
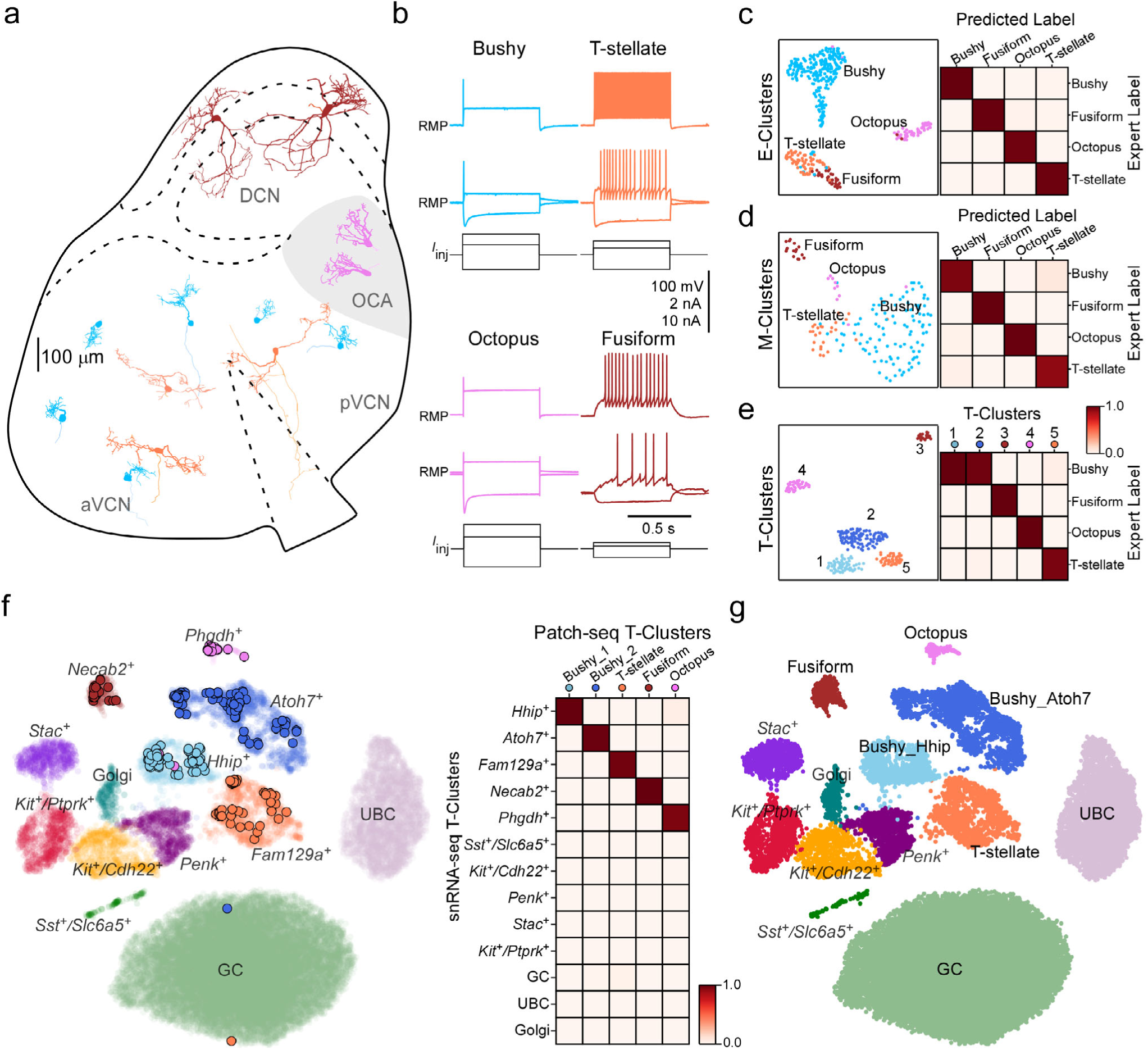
Using Patch-seq to identify transcriptomic correspondences for excitatory neurons. **a**, Examples of reconstructed excitatory neurons positioned at their site of origin in representative mouse CN (sagittal view), colored by cell type identity. **b**, Example responses of CN excitatory cell types to current steps. Bottom traces show the injected currents. Scalebar: 100 mV for potential, 2 nA for injected currents for all cells except octopus cells (10 nA). **c**, Left: UMAP visualization of 404 CN excitatory neurons clustered by over-dispersed physiological features (E-cluster), colored by expert cell type. n=243 for bushy, n=67 for T-stellate, n=58 for octopus, n=36 for fusiform. Right: Confusion matrix shows performance (an average of 99.2% accuracy) of the cell-type classifier trained with physiological features. See Extended Data Fig. 4 for more details. **d**, Left: UMAP visualization of 157 CN excitatory neurons clustered by over-dispersed morphological features (M-cluster), colored by expert cell type. n=105 for bushy, n=16 for T-stellate, n=10 for octopus, n=26 for fusiform. Right: Confusion matrix shows performance (an average of 95.5% accuracy) of the classifier trained with morphological features. See Extended Data Fig. 4 for more details. **e**, Left: UMAP visualization of 293 Patch-seq excitatory neurons clustered by over-dispersed genes, colored by transcriptomic cluster (T-cluster). Right: the matrix shows proportion of Patch-seq cells assigned to a specific cell type (the expert classification) among each T-cluster. All cells in T-cluster 1 (n= 67/67) and almost all cells in T-cluster 2 (n=111/113) are bushy cells; all cells in T-cluster 3 are fusiform cells (n=22/22); all cells in T-cluster 4 are octopus cells (n=43/43); almost all cells in T-cluster 5 are T-stellate cells (n=46/48). **f**, Left: Patch-seq neurons (n=293) mapped to snRNA-seq UMAP space as shown in Fig. 1b. Individual dots represent each cell, colored by T-cluster. Right: matrix shows proportion of Patch-seq cells assigned to each T-cluster in (**e**) mapped to one of 13 clusters in UMAP space. **g**, UMAP visualization and annotation of molecular cell types with established morpho-electrophysiological types in CN.

DEG analysis was performed to find marker genes for each cell type (or cluster, Extended Data Table 1), while FISH and immunohistochemistry validated their expression and cell type-specificity by leveraging the known anatomical location of each cell type (Methods). With these approaches, a suite of novel discriminatory markers was identified for octopus cells (Extended Data Fig. 6a-b and Extended Data Fig. 7a, 8a, and Extended Data Table 1), a major projection neuron for coincidence detection^29,30^. Antibodies to one of these markers, PHGDH, labeled neurons (i.e., cells with a large soma) restricted to the octopus cell area (OCA) (Extended Data Fig. 6c). Indeed, when patch-clamp recordings from the OCA to label octopus cells with biocytin and *post hoc* immunostaining performed for PHGDH, most labeled cells (12/13) were immunopositive for PHGDH (Extended Data Fig. 6d). Similarly, *Necab2* and *Pbfibp1*, among other genes, were confirmed as major markers for fusiform cells of the DCN (Extended Data Fig. 6e-f, Extended Data Fig. 7b, 8b and Extended Data Table 1). Immunostaining combined with patch-clamp analysis provided translation-level evidence for *Necab2* as a highly selective marker for the entire population of fusiform cells (Extended Data Fig. 6g-h).

### Bushy cells consist of two major molecular subtypes

In anatomical and physiological studies, bushy cells have been divided into two populations, spherical and globular bushy cells (SBCs and GBCs, respectively), based largely on differences in their axonal projections^31^. Generally, SBCs are located anteriorly in VCN while GBCs are more posterior^6^. Some investigators have proposed two subtypes of SBCs (small and large), based on their axonal targets in lateral and medial superior olive^6^, although it is not clear how distinct these populations are. Our initial analysis identified two transcriptomically distinct bushy cell populations marked by high levels of *Atoh7* or *Hhip* (*Atoh7*^+^ and *Hhip*^+^, Fig. 3a-b), which may correspond to SBCs and GBCs. We therefore performed FISH co-staining for *Atoh7* and *Hhip* to determine the spatial location of these cell groups. Given that *Sst* is specifically expressed in the *Atoh7* subtype rather than the *Hhip* subtype (Fig. 3b-c) and is expressed preferentially in SBCs rather than GBCs in VCN^32^, we also performed FISH co-labeling for *Sst* and *Atoh7*, and for *Sst* and *Hhip*. Both *Atoh7* and *Hhip* signals were almost exclusively restricted to VCN (Fig. 3c), in line with our snRNA-seq data (Fig. 3b, Extended Data Fig. 5a-b, d) and Allen ISH data (Extended Data Fig. 8c). Furthermore, *Atoh7^+^* cells were predominantly localized in the rostral anterior VCN (AVCN), while *Hhip^+^* cells densely populated the posterior VCN (PVCN) and caudal AVCN, and less so rostrally (Fig. 3c), similar to SBCs and GBCs in terms of their typical locations^31,33,34^. In addition, *Sst* signals mostly overlapped with *Atoh7* and much less so with *Hhip*, and double-labeled cells (*Sst^+^/Atoh7^+^*and *Sst^+^/Hhip^+^*) were restricted to rostral AVCN (Fig. 3c), further validating two molecularly distinct bushy cell subtypes marked by *Atoh7* or *Hhip*. Based on the preferential location of each molecular subtype with differential *Sst* expression, the *Atoh7* subtype may correspond to SBCs, while the *Hhip* subtype may correspond to GBCs.

**Fig. 3:**
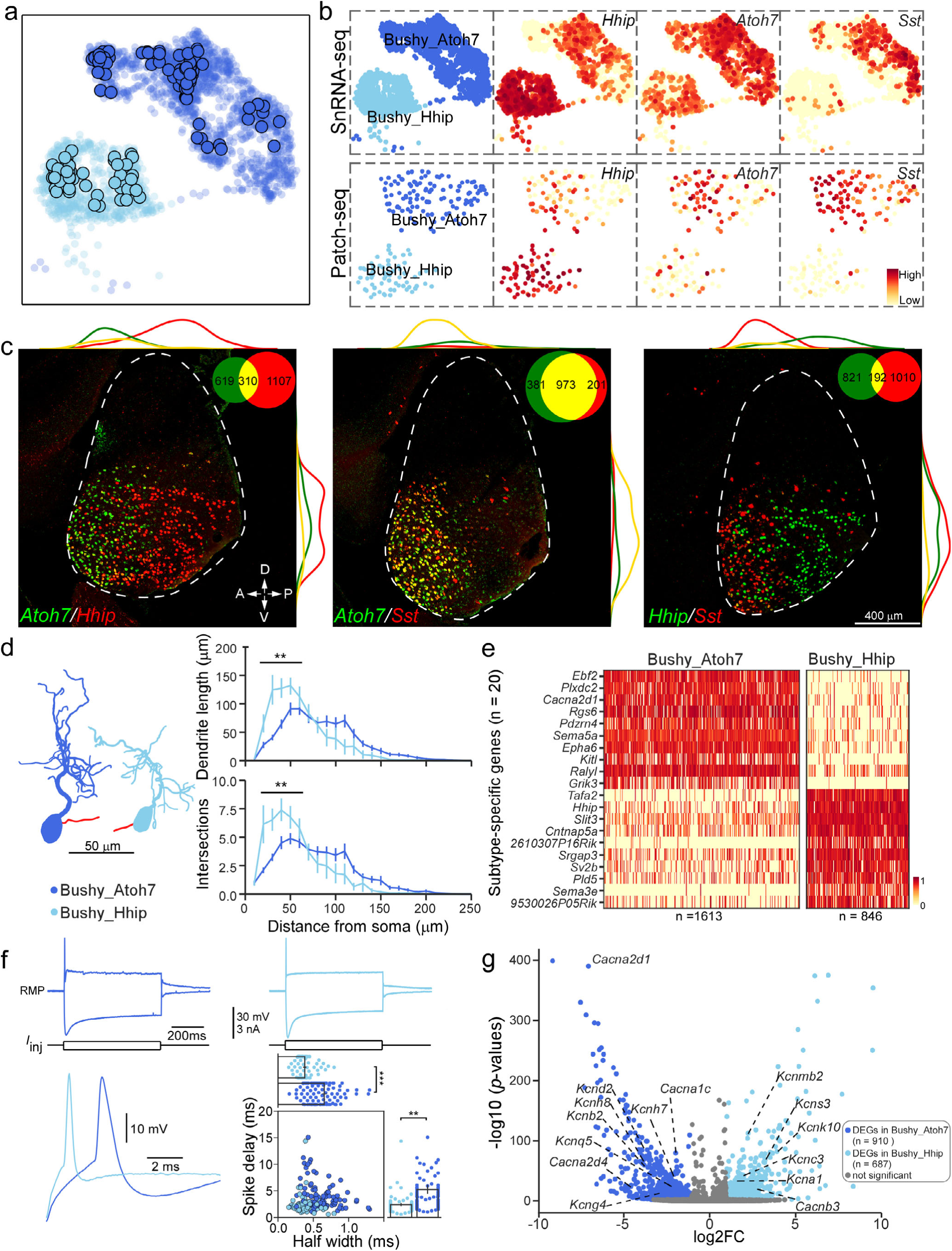
Two molecular subtypes of bushy cells. **a**, UMAP visualization of *Hhip*^+^ and *Atoh7*^+^ clusters and distribution of Patch-seq bushy cells in UMAP space, colored by cluster. Individual dots represent each patch-seq cell. **b**, UMAP visualization of two bushy cell clusters and normalized expression of their marker genes, *Hhip*, *Atoh7,* and *Sst*. Top: snRNA-seq, Bottom: Patch-seq. **c**, FISH co-staining for *Atoh7* and *Hhip* (left), *Atoh7* and *Sst* (middle), or *Hhip* and *Sst* (right) in CN sagittal sections. Insets show total number of single-labeled cells and double-labeled cells. Lines along the images indicate density of single-labeled or double-labeled cells along two axes of CN. Scale bar applies to all images. **d**, Left: A representative morphology of *Hhip*^+^ or *Atoh7*^+^ bushy cell. Axon (shown in red) is truncated. Right: Sholl analysis of *Hhip*^+^ or *Atoh7*^+^ bushy cell dendrites. n*=*20 for *Hhip*^+^ cells, n=51 for *Atoh7*^+^ cells, \*\**p* < 0.01, with two-way mixed model ANOVA. **e**, Heat map displaying normalized expression of the top 20 DEGs for two subtypes. Scale bar: Log2 (Expression level). The number of cells in each subcluster is indicated below the figure. **f**, Top: example responses of *Atoh7*^+^ bushy cell (left) or *Hhip*^+^ bushy cell (right) to a hyperpolarized current step and a near-threshold depolarizing current step. Bottom-left: individual APs from the *Hhip*^+^ and *Atoh7*^+^ cells shown above, aligned with onset of the depolarizing current. Bottom-right: two most-discriminating features for *Hhip*^+^ cells and *Atoh7*^+^ cells, spike delay and spike duration (half-width). Plotting these two features separates the majority of *Hhip*^+^ cells from *Atoh7*^+^ cells. n*=*67 for *Hhip*^+^ cells, n=110 for *Atoh7*^+^ cells, \*\**p* < 0.01; \*\*\**p* < 0.001; t-test. **g**, Volcano plot showing the log2 fold change (log2FC) and −log10(p-value) of detected genes comparing the two subtypes. Among DEGs are 15 genes encoding voltage-gated ion channels. See Extended Data Fig. 9 for more analysis.

To determine if molecular subtypes of bushy cells are associated with unique physiological and morphological features, we examined the properties of each Patch-seq cell that was assigned to the *Hhip* or *Atoh7* subtype. Two subtypes exhibited similar soma size and shape, but there were differences in their dendritic arborization (Extended Data Table 3). While the majority of *Hhip* bushy cells have only one primary dendrite (85%, 17 out of 20 cells), more than half of *Atoh7* bushy cells have two or even three primary dendrites (∼53%, 27 out of 51 cells), each with its own secondary and filamentous branches (Extended Data Table 3). In addition, *Hhip* bushy cells had short dendritic trunks capped with relatively compact dendritic tufts, while *Atoh7* bushy cells had longer dendritic trunks that bifurcated and sent off diffuse, thin dendritic processes (Fig. 3d and Extended Data Table 3). Dendrites of either *Atoh7* bushy cells or *Hhip* bushy cells were not oriented in any particular direction within the VCN.

Molecularly, two bushy cell subtypes were best discriminated by cell-adhesion molecule (CAM) genes (e.g., *Plxdc2*, *Sema5a*, *Epha6*, *Kitl, Slit3*, *Cntnap5a*, *Sema3e*), suggestive of their distinct connectivity and projection patterns (Fig.3e). Bushy cells are biophysically specialized for encoding precise timing by expressing a prominent low-threshold K^+^ current and a hyperpolarization-activated current, which together bestow single-spike firing, and low input resistance and short time constants enabling brief and sharply timed synaptic responses^35–37^. Both bushy cell subtypes exhibited single-spike firing property, but they were electrophysiologically distinguishable with significant differences in numerous features, with spike delay and half-width being the most prominent (Fig. 3f, Extended Data Fig. 9a, Extended Data Table 3). *Hhip* bushy cells had lower input resistances and shorter membrane time constant (thus shorter delays to firing onset), deeper and quicker voltage sags in response to hyperpolarizing currents, and fewer spikes in response to suprathreshold depolarizing currents than *Atoh7* bushy cells (Fig. 3f, Extended Data Table 3), indicative of larger near-rest outward current with faster kinetics. These properties match a higher expression of ion channels active in that voltage range (*Kcna1, Hcn1)* in *Hhip* bushy cells (Fig. 3g, Fig. 7b)^35,38^. In addition, *Hhip* bushy cells had faster depolarization and repolarizations (larger dV/dt) resulting in narrower spikes (Fig. 3f, Extended Data Table 3), matching their enrichment of Kv3 (*Kcnc3*) (Fig. 3g), a potassium channel enabling brief APs for auditory neurons to follow high-frequency input with temporal precision^39–41^. In contrast, *Atoh7*^+^ cells had a higher expression of Kv2 (*Kcnb2*) in addition to a lower expression of Kv1 (Fig. 3g, Fig. 7b), which may enable them to fire more spikes than *Hhip*^+^ cells^42^.

Patch-seq profiles (with more detected genes than snRNA-seq) mapped to two main subtypes of bushy cells form distinct subclusters in snRNA-seq UMAP space (Fig.3a), suggesting further bushy cell-type molecular heterogeneity. Subsequent subclustering analysis identified three subpopulations of *Atoh7^+^*bushy cells (*Atoh7*/*Dchs2*, *Atoh7*/*Tox*, and *Atoh7*/*Sorcs3*) (Extended Data Fig. 9b) and two subpopulations of *Hhip^+^* bushy cells (*Hhip*/*Calb1* and *Hhip*/*Galnt18*) (Extended Data Fig. 9f). To further assess these subpopulations, we examined the properties of Patch-seq cells that were assigned to each subcluster, including their anatomical locations. Within *Atoh7^+^* bushy cells, *Atoh7*/*Dchs2* cells appear to be restricted to the nerve root area and PVCN, while *Atoh7*/*Sorcs3* cells and *Atoh7*/*Tox* cells appear to have less spatial preferences (Extended Data Fig. 9c). Electrophysiologically, *Atoh7*/*Dchs2* cells displayed distinct firing properties compared to *Atoh7*/*Sorcs3* and *Atoh7*/*Tox* cells, which shared similar firing patterns (Extended Data Fig. 9d, Extended Data Table 3). Morphologically, *Atoh7*/*Dchs2* cells generally had the longest dendritic trunks (stem) among three subclusters (Extended Data Fig. 9e, Extended Data Table 3). Within *Hhip^+^* bushy cells, *Hhip*/*Calb1* cells were preferentially localized in the posteroventral VCN, while *Hhip*/*Galnt18* cells were preferentially localized in the dorsoanterior VCN (Extended Data Fig. 9g). *Hhip/Calb1* cells exhibited distinct firing properties from *Hhip*/*Galnt18* (Extended Data Fig. 9h, Extended Data Table 3). Additionally, the *Hhip/Calb1* cells generally had shorter dendritic projections, but more complex dendritic tufts compared to *Hhip*/*Galnt18* cells (Extended Data Fig. 9i, Extended Data Table 3). These analyses support functionally and topographically distinct subpopulations within each bushy cell subtype.

### T-stellate cells consist of two subtypes

Patch-seq mapping indicated that the *Fam129a*^+^ cluster corresponds to T-stellate cells (Fig. 2f-g). This cluster comprised two visibly separate lobes on the UMAP, suggesting two sub-populations of T-stellate cells (Fig. 1c and 4a). With more than 1100 DEGs identified between two putative sub-populations, we focused on *Dchs2* and *Fn1*, two top DEGs expressed almost exclusively in one of the two sub-populations with high coverage (Fig. 4a-b, Extended Data Fig. 10a-b). To validate their expression and determine how well these two genes separate T-stellate cell subpopulations, we performed FISH co-labeling for *Fam129a* and *Dchs2*, and for *Fam129a* and *Fn1*. *Fam129a* signals were restricted to VCN (Fig. 4c-d) and almost exclusively expressed in excitatory neurons (data not shown), thus validating *Fam129a* as a general marker for T-stellate cells (Fig. 2f, Extended Data Figs. 5a-b, 5d and 8d). Only a subset of *Fam129a^+^* neurons co-expressed *Fn1* (*Fam129a^+^*/*Fn1^+^*), and these double-labeled neurons were exclusively localized in the AVCN (Fig. 4c). Similarly, only a subset of *Fam129a ^+^* neurons co-expressed *Dchs2*, and these *Fam129a^+^*/*Dchs2^+^* neurons were restricted to the VCN nerve root area and to PVCN (Fig. 4d). These data suggest two spatially separated subpopulations of T-stellate cells. To further substantiate this conclusion, we examined the location of Patch-seq cells assigned as T-stellate cells. These cells were mapped to either the *Dchs2^+^* subcluster or the *Fn1^+^* subcluster almost equally (Fig. 4a), allowing for an unbiased comparison of their spatial locations. *Fn1^+^* T-stellate cells were restricted to the AVCN, while *Dchs2^+^* T-stellate cells were predominantly localized in the PVCN and the nerve root area of the VCN. Importantly, these two subpopulations were spatially non-overlapping in the VCN (Fig. 4e), indicating that T-stellate cells could be classified into two subtypes by their anatomical locations. The existence of two T-stellate cell subtypes was further supported by their distinct morphological and electrophysiological properties. While both *Fn1^+^* or *Dchs2^+^* T-stellate cells exhibited typical morphology of T-stellate cells (Fig. 4f)^43^, there were subtle differences in their dendritic arborization (Extended Data Table 4). The dendrites of *Fn1*^+^ cells in general branched earlier and more profusely around the soma compared to that of *Dchs2^+^*cells, while the dendrites of *Dchs2*^+^ cells arborized more extensively away from the soma with more branched, complex endings (Fig. 4f-g, Extended Data Table 4). In addition, the major dendrites and the terminal arbors of *Fn1*^+^ and *Dchs2*^+^ cells had distinct preferential angles with respect to their soma (Fig. 4h). Given that the major dendrites and the terminal arbors of T-stellate cells in general run parallel to tonotopically arranged ANFs (Fig. 4f) whose direction in each subregion is well-established^44,45^, such differences between two subtypes reflect their distinct anatomical locations in the VCN, further substantiating the conclusion that the subtypes of T-stellate cells are defined by their anatomical locations.

**Fig. 4:**
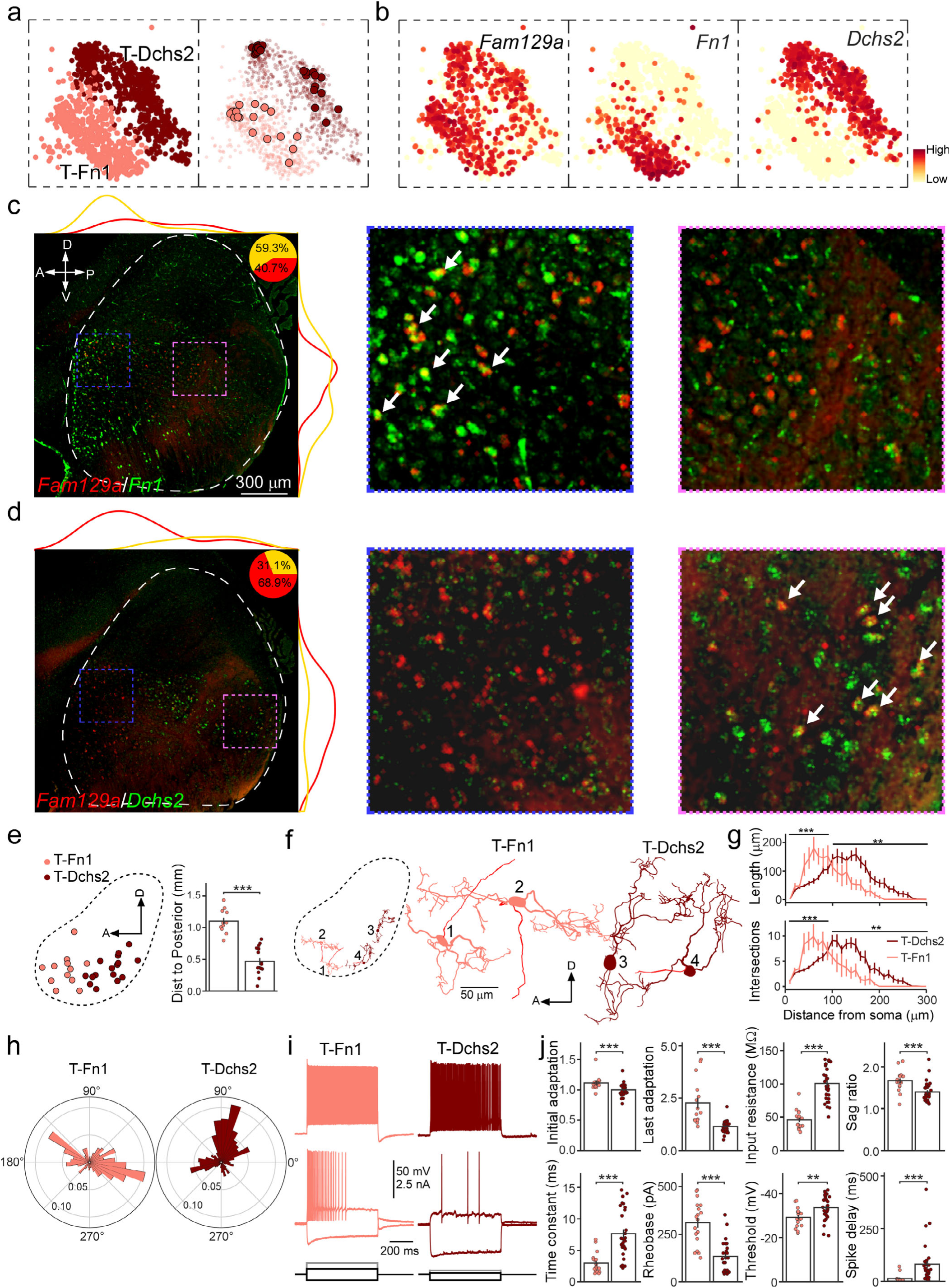
Two subtypes of T-stellate cells. **a**, Left: UMAP visualization of two subclusters of *Fam129a^+^* neurons (marked by high expression of *Dchs2* or *Fn1*:T-Fn1 and T-Dchs2), colored by subcluster. Right: All Patch-seq cells mapped onto *Fam129a^+^*cluster on the left. Individual dots represent each patch-seq cell. **b**, UMAP visualization of normalized expression of discriminatory markers (*Fam129a*, *Fn1*, *Dchs2*) for all T-stellate cells and their two subclusters. **c**, FISH co-staining for *Fam129a* and *Fn1* in a CN sagittal section. Inset pie chart: proportion of double-labeled cells among single-labeled *Fam129a*^+^ cells. Lines along images depict density of single-labeled or double-labeled neurons along the two axes of CN. Two images on the right are zoomed-in views of boxed regions on the left. Arrows point to double-labeled cells. **d**, FISH co-staining for *Fam129a* and *Dchs2* in a CN sagittal section. Two images on the right are zoomed-in views. **e**, Top: 2D spatial projection of Patch-seq T-Fn1 cells and T-Dchs2 cells onto a sagittal view of CN, colored by the subtypes. Bottom: Comparison of the distance to CN posterior edge between T-Fn1 and T-Dchs2. n=12 for T-Fn1, and n=16 for T-Dchs2; \*\*\**p* < 0.001, t-test. **f**, Left: Representative morphology of two *Fn1^+^* cells and two *Dchs2*^+^ cells in a CN sagittal view (soma locations). Axons in red. Right: zoomed-in of four T-stellate cells shown on the left. **g**, Sholl analysis of T-Fn1 and T-Dchs2 dendrites. n=6 for T-Fn1, and n=20 for T-Dchs2, \*\**p* < 0.01, \*\*\**p* < 0.001, two-way mixed model ANOVA. **h**, Polar histograms showing the distribution of dendritic branch termination points with respect to soma of T-Fn1 and T-Dchs2. The terminal distribution of T-Fn1 has two peaks at 145° and 345°, while the terminal distribution of T-Dchs2 has two peaks at 75° and 195°. *p* < 0.001 for difference between the distributions using the Kolmogorov-Smirnov statistic. **i**, Example responses of two subtypes (T-Fn1, T-Dchs2) to current steps. **j**, Comparison of electrophysiological features between two T-stellate subtypes. n=16 for T-Fn1, and n=29 for T-Dchs2, \*\**p* < 0.01, \*\*\**p* < 0.001, t-test.

T-stellate cells fire continuously in response to suprathreshold inputs, a biophysical property required for encoding sound intensity^42,50^. Both *Fn1*^+^ and *Dchs2*^+^ T-stellate cells fired tonically, as reported previously, but were physiologically distinguishable showing differences in numerous features (Fig. 4i-j, Extended Data Fig. 10c and Extended Data Table 4). Among these, spike delay (latency to first spike at rheobase), time constant, and input resistance alone separated almost all *Dchs2*^+^ cells from *Fn1*^+^ cells (Extended Data Fig. 10d). Such striking physiological distinctions well aligned with their top DEGs being dominated by numerous genes encoding potassium channels (Extended Data Fig. 10e). Notably, *Fn1*^+^ cells were enriched for expression of multiple leak potassium channels including K_2P_9.1 (*Kcnk9*), K_2P_10.1 (*Kcnk10*), and K_2P_12.1 (*Kcnk12*) and subthreshold-operating Kv channels including Kv7.4 (*Kcnq4*), Kv10.2 (*Kcnh5*) and Kv12.1 (*Kcnh8*), matching their lower input resistance and smaller time constant compared to *Dchs2*^+^ cells^46,47^(Fig. 4g, Extended Data Fig. 10d). In contrast, *Dchs2*^+^ cells were enriched for expression of Kv2 and Kv3 (*Kcnb2*, *Kcnc2*), two potassium channels required for auditory neurons to maintain high-frequency repetitive firing^39,42,48^, matching their more ‘fast-spiking’ phenotype (e.g., no adaptation) compared to *Fn1*^+^ cells.

### Molecular profiles of inhibitory neurons in CN

We also sought to determine the correspondence of glycinergic clusters to established inhibitory cell types in the CN. Among the six glycinergic/inhibitory clusters (including Golgi cells), four expressed both *Slc6a5* and genes for GABA synthetic enzymes (*Gad1/2*, Fig. 1c), consistent with dual transmitter phenotypes described in CN^25,49,50^. The other two clusters, *Slc6a5*^+^/*Sst*^+^ and *Penk^+^* cluster, expressed only *Slc6a5* and no *Gad1/2*, indicative of being pure glycinergic populations (Fig. 1c). Notably, the *Slc6a5*^+^/*Sst*^+^ cluster likely corresponds to D-stellate cells in VCN, a known pure glycinergic population with specific *Sst* expression^10,50^. A definitive correspondence of glycinergic clusters to established inhibitory cell types (Fig. 1a and 5a) was then determined by targeting glycinergic cell types for Patch-seq (see Methods). These included vertical, cartwheel, and superficial stellate cells (SSC) in DCN, and L- and D-stellate cells in VCN (Fig. 1a and 5a), each of which resides in distinct anatomical locations (except for L- and D-stellate cells) and exhibits unique morpho-electrophysiological features (Fig. 5a-b). As with excitatory neurons, we validated the expert classification and cell-type assignment of each Patch-seq cell. Both the random forest classifier and clustering-based k-means classifier demonstrated high accuracy in distinguishing each cell type pair using morphological features (see Methods, Fig. 5c, and Extended Data Fig. 4c-d). The performance was also robust using electrophysiological features, especially when anatomical location information was incorporated (see Methods, Fig. 5d, and Extended Data Fig. 4e-h). This robust performance across multiple modalities and methods supports cell-type assignment for each Patch-seq cell. UMAP projection of these Patch-seq cells resulted in 5 clusters which were then mapped onto the snRNA-seq dataset, successfully identifying transcriptomic clusters for established inhibitory cell types (Fig. 5e-g). As with excitatory neurons, we validated the molecular correspondences to inhibitory neurons using FISH. We validated the *Stac^+^* cluster as cartwheel cells by FISH staining for *Stac* or *Kctd12,* two major putative marker genes for cartwheel cells (Fig. 5h-i, Extended Data Fig. 5c,5e, Extended Data Table 1, and Extended Data Fig. 8e). FISH staining for *Penk* in wild-type mice and Cre-positive cells in an existing Penk-Cre mouse line validated the *Penk*^+^ cluster as vertical cells (Fig. 5i-j, Extended Data Fig. 11a), consistent with these inhibitory neurons being pure glycinergic^51,52^. FISH staining also validated Penk-Cre mice as a specific Cre driver for vertical cells, and thus electrophysiology and morphology of Cre-positive cells in DCN all matched with vertical cells (Methods; Fig. 5k-l). Interestingly, in addition to labeling vertical cells, *Penk* expression was also enriched in glycinergic neurons in the ‘small-cell cap’ (Fig. 5i-k, Extended Data Fig. 8f), a narrow CN region between the granule-cell and magnocellular domains of the VCN^53^. This topographically unique inhibitory subgroup was devoid of *Gad1* (i.e., purely glycinergic, Extended Data Fig. 11b), and thus genetically more similar to vertical cells than to other small glycinergic neurons in VCN (i.e., *Gad1*^+^ L-stellate cells, Fig. 1c, Extended Data Fig. 11c, also see below). The *Penk*^+^ cluster thus includes both vertical cells and glycinergic cells in the small-cell cap.

**Fig. 5:**
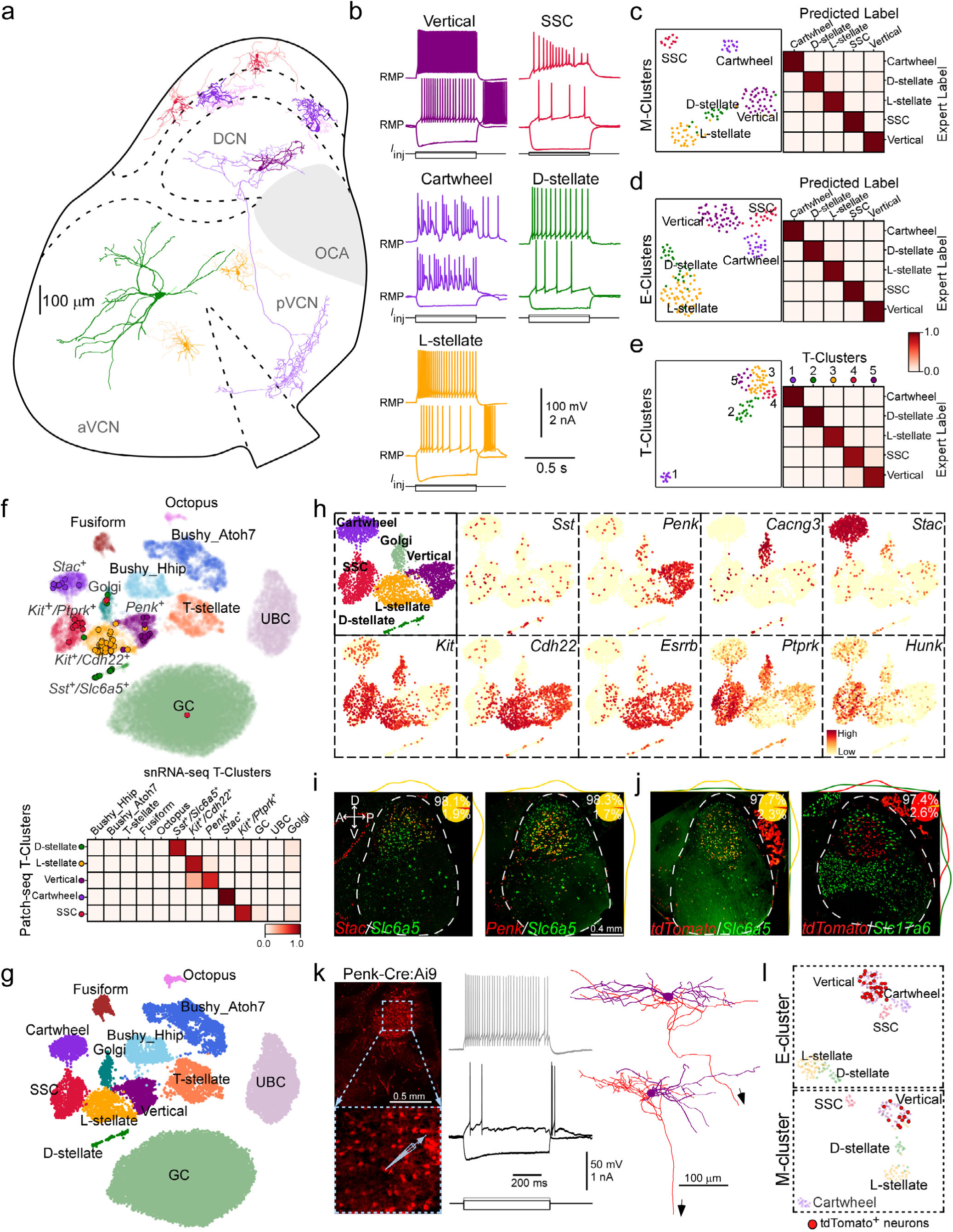
Annotation of molecularly distinct glycinergic cell types in CN. **a**, Examples of reconstructed inhibitory neurons positioned at their site of origin in a representative mouse CN section (a sagittal view), colored by cell type identity. **b**, Example responses of CN inhibitory cells to current steps. **c**, Left: UMAP visualization of 116 CN inhibitory neurons clustered by select morphological features (M-cluster), colored by expert cell type. n=38 for Vertical, n=13 for SSC, n=15 for Cartwheel, n=17 for D-stellate, n=33 for L-stellate. Right: Confusion matrix shows performance (average of 96.9% accuracy) of the classifier trained on morphological features. **d**, Left: UMAP visualization of 172 inhibitory neurons clustered by electrophysiological features and anatomic locations, colored by expert cell type. n=50 for Vertical, n=21 for SSC, n=22 for cartwheel, n=26 for D-stellate, n=53 for L-stellate. For anatomic locations, 0 represents VCN and 1 represents DCN in the cluster analysis. Right: Confusion matrix shows performance (an average of 97.9% accuracy) of the classifier trained with electrophysiological features and anatomic locations. Colored by the expert cell type. See Extended Data Fig. 4e-f for the analysis with electrophysiological features only. **e**, Left: UMAP projection of 94 Patch-seq inhibitory neurons clustered by over-dispersed genes, colored by transcriptomic clusters (T-clusters). Right: The matrix shows proportion of Patch-seq cells in each T-cluster assigned to a specific cell type. All cells in T-cluster 1 are cartwheel cells (n=12/12); almost all cells in T-cluster 2 are D-stellate cells (n=20/21); all cells in T-cluster 3 are L-stellate cells (n=35/35); almost all cells in T-cluster 4 are SSCs (n=10/12), and almost all cells in T-cluster 5 are vertical cells (n=12/14). **f**, Top: Patch-seq inhibitory neurons (n=94) mapped onto snRNA-seq UMAP space as shown in Fig. 1b. Individual dot represent a neuron, colored by T-cluster. Bottom: Matrix shows proportion of Patch-seq cells assigned to each T-cluster in (**e**) mapped onto one of 13 distinct clusters in UMAP space. **g**, UMAP visualization and annotation of the neuronal clusters with established morpho-electrophysiological types (including both excitatory and inhibitory) in CN. **h**, UMAP visualization of all glycinergic neurons and normalized expression of marker genes for each inhibitory cell type in CN. Each cluster is annotated by morpho-electrophysiological cell types. **i**, FISH co-staining for *Slc6a5* and *Stac* (left), or for *Slc6a5* and *Penk* (right) in sagittal sections. Inset pie charts show proportion of double-labeled cells among single-labeled cells. Yellow lines along the images indicate density of double-labeled neurons along the two axes of CN. **j**, FISH co-staining for *Slc6a5* and *tdTomato* (left), or for *Slc17a6* and *tdTomato* (right) in CN sagittal sections in Penk-Cre:Ai9 mice. Inset pie charts show proportion of double-labeled cells among all *tdTomato*^+^ cells. **k**, Left: micrographs show labeled cells targeted for recording in Penk-Cre:Ai9 mice. Middle: example responses of a labeled neuron to current steps. Right: two morphologies of labeled neurons. Axons (truncated) project towards the VCN (arrows). **l**, Top: Distribution of labeled cells (red circle) in electrophysiological UMAP space of all inhibitory neurons. Almost all labeled cells (30/31) clustered with vertical cells. Bottom: Distribution of labeled cells in morphological UMAP space. All labeled cells clustered with vertical cells.

FISH staining further validated the *Sst^+^*/*Slc6a5^+^*cluster as D-stellate cells that could be discriminated with a combination of two genes or even single genes^10^ (Extended Data Figs. 5e, 11d, and Extended Data Table 1). To validate *Kit^+^/Cdh22^+^*cluster as L-stellate cells, we performed co-labeling for *Kit* and *Cdh22*, and for *Kit and Esrrb*, two combinations predicted to distinguish L-stellate cells (Fig. 5f-h, Extended Data Fig. 5e). In line with snRNA-seq data (Fig. 1c, 5h, and Extended Data Fig. 5e), *Kit^+^* cells were almost exclusively glycinergic with a smaller soma size compared to *Kit^-^* glycinergic neurons (Extended Data Fig. 11e), and *Kit^+^* neurons labeled by *Cdh22* or *Essrb* transcripts (*Kit^+^/Cdh22^+^* or *Kit^+^/Essrb^+^*) were mainly detected in the VCN (Extended Data Fig. 11f), consistent with the location of L-stellate cells. Interestingly, double-labeled glycinergic neurons were sparsely detected in DCN as well (Extended Data Fig. 11f), suggesting that the *Kit^+^*/*Cdh22^+^*cluster may also include glycinergic neurons in the main body of DCN that are neither cartwheel cells nor vertical cells (Extended Data Fig. 11g-h), a hitherto undescribed cell type. Finally, to validate *Kit^+^*/*Ptprk^+^* clusters as SSCs, we performed co-staining for *Kit* and *Ptprk*, and for *Kit* and *Hunk,* two potential combinations to discriminate SSCs (Fig. 1c, 5f-h and Extended Data Fig. 5e). In line with our analysis, double-labeled glycinergic neurons (*Kit^+^/Ptprk^+^* or *Kit^+^/Hunk^+^*) were mostly detected in the molecular layer of the DCN (Extended Data Fig. 11i). Double-labeled cells were also sparsely detected at the VCN, likely reflecting the expression of *Hunk* or *Ptprk* in a small subset of L-stellate cells (Fig. 5h and Extended Data Fig. 11i).

### Molecular basis of CN cell type identity and wiring specificity

Our analysis indicates that all cell types in CN with anatomical and physiological identity (each with unique morphophysiological features, connectivity, projection patterns, and function)^4–6^ can be defined by their transcription profiles, suggesting that cellular specializations are rooted in differential gene expressions (Fig. 5g). Gene profiles conferring neuronal phenotypes are often dictated by differential expression of a small set of gene families^54–56^, and determined by the concerted action of transcription factors (TFs)^57–60^. We thus performed supervised computational screen using MetaNeighbor algorithm^54^ to identify gene families whose differential expression could reliably predict CN neuron types (Methods). We found that the top-performing gene families were dominated by three functional categories including transcription factors (TF) (Methods, Fig.6a-b, Extended Data Table 5), in line with the notion that TFs are key regulators of cell type identity^57–60^. We therefore set out to characterize cell types by their TF activity and ask if CN cell types are defined by specific yet overlapping TF network features. With a more comprehensive TF database with precisely annotated TF families^61^, we found differentially expressed TFs among CN cell types were predominantly from the C2H2 zinc finger family (Methods, Extended Data Table 5). Approximately one third of the most differentially expressed TFs belong to C2H2-Zinc finger TFs, a pattern of TF expression that appears unique to this brain region^54,62,63^. We did not find TFs whose expression correlates strictly with each cell type, indicating the importance of combinatorial TF codes in specifying cell type identity (Fig.6c-d; Extended Data Table 6). We identified a set of TFs that are selectively co-expressed in specific cell types and hence may define identities of cell types (Fig.6c-d, Extended Data Table 6). Many of these TFs are also expressed during early CN development, including in progenitor cells (Fig. 6c-d, highlighted in red, Extended Data Table 7)^64–68^, suggesting developmental continuity of TF expression from embryonic precursors to mature neurons. For example, while TFs like *Atoh7, Pax6, Lhx9, Sox11,* and *Klf3* are each enriched in a specific mature projection neuron type (Fig. 6c), they exhibit early and broad expression within the developing Atoh1 lineage that gives rise to these projection neurons^64,66,67^. These results suggest that developmental transcription programs, initiated early in post-mitotic differentiation, become progressively restricted to and maintained within specific mature cell types of the same lineage^54^. Similarly, many TFs enriched in each CN inhibitory cell type have been shown to be expressed during early neural development and play a role in the differentiation and specification of GABAergic/glycinergic neurons (Fig. 6d, highlighted in red, Extended Data Table 7). Our results suggest that most of the developmental programs for cellular differentiation and specification persist in mature neurons.

**Fig. 6:**
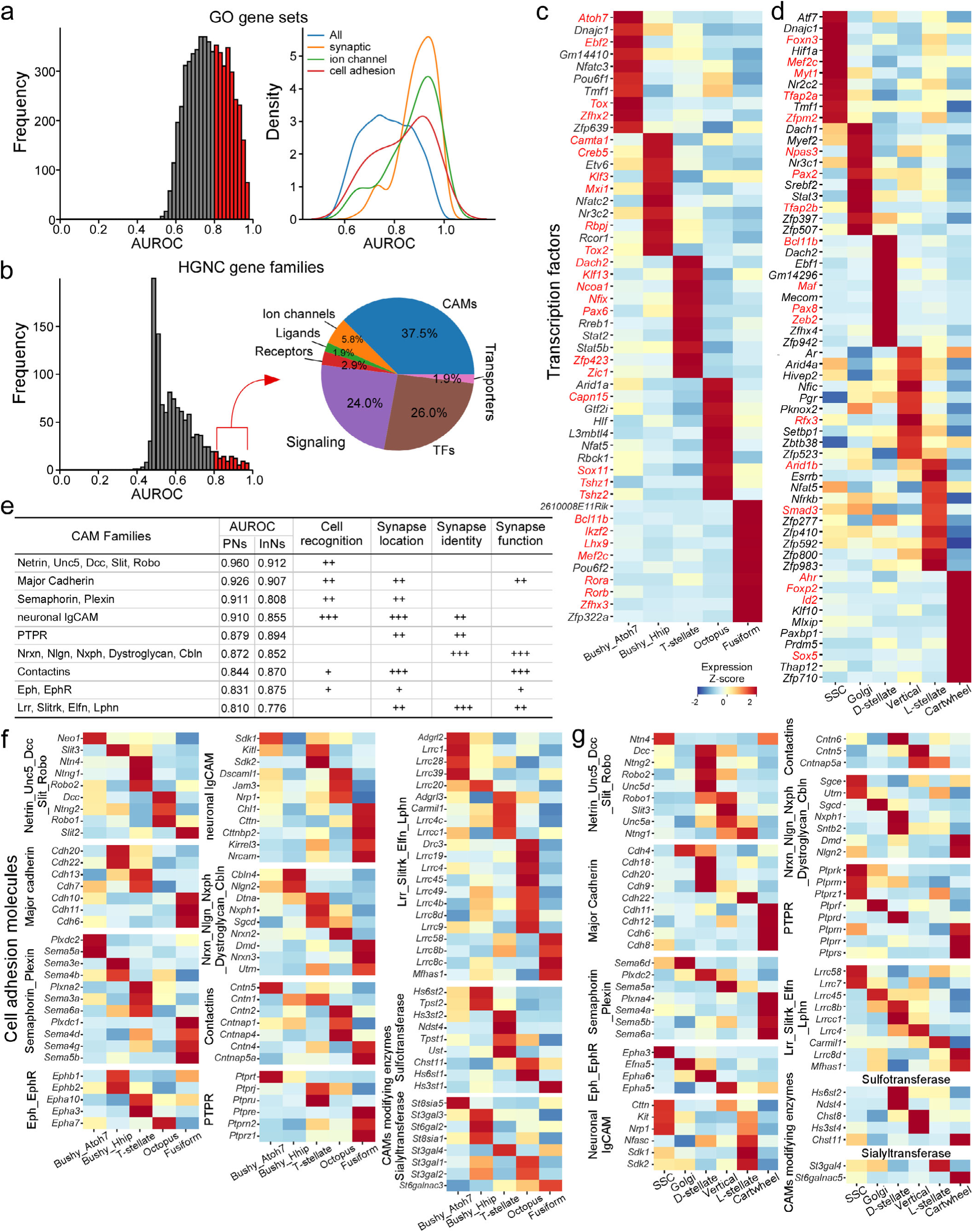
Molecular determinants of CN cell type identity. **a**, Distribution of AUROC values of 5,735 GO terms across all 13 cell types in the snRNA-seq dataset. Red, AUROC ≥ 0.8. Right: GO term probability density by keywords: “synaptic”, “cell adhesion” and “ion channel” are skewed with AUROC ≥ 0.8. **b**, AUROC value distribution of 1,424 HGNC gene families across all 13 cell types. 104 families with AUROC≥ 0.8 (red bars) are classified into 7 categories (pie chart). See Extended Data Table 5 for more details. **c**, Heatmap displays the z-scored expression levels of the top 10 specific TFs for each excitatory projection neuron type. TFs highlighted in red show expression in the developing cochlear nucleus, including progenitors and distinct developmental stages of projection neurons (see Extended Data Table 7 for reference support). **d**, Heatmap showing the z-scored expression levels of the top 10 specific TFs for each inhibitory cell type. TFs in red have been shown to involve in the differentiation and specification of glycinergic/GABAergic interneurons in cochlear nucleus or other brain regions (see Extended Data Table 7 for reference support). **e**.The top-performing cell-adhesion molecule (CAM) families, their AUROC value, and their roles in synaptic connectivity. “+” denotes the degree of involvement in the listed function. **f**, Heatmap showing the differential expression of 9 CAM and 2 carbohydrate-modifying enzyme families across excitatory projection neuron types. **g**, Heatmap showing the differential expression of 9 CAM and 2 carbohydrate-modifying enzyme families across inhibitory cell types

Cell type identity is often determined by a small set of core TFs that exhibit synergistic transcriptional regulation^58,69–76^. We thus leveraged a new computational method to predict the core identity TFs for each CN cell type (synergistic identity TF cores), a method capitalizing on key principles of core TF expression: relatively high and cell-type-specific expression and transcriptional synergy^69,77^. This analysis predicted a distinct combination of core TFs for each cell type (Extended Data Table 8). Most of these predicted TFs are recognized as essential for neuronal differentiation and specification^78–88^. Many (*Pax6*, *Ebf2*, *Rorb*, *Tshz3*, *Bnc2, Tead1, Meis1, Bcl11a, Prox1, Zfp536, Zfhx3, Ikzf2, Zfp322a*) are known for maintaining cell type identity^60,81,89–101^, and deletion of these TFs in postnatal age or adult result in the loss of markers and physiological properties characteristic to their identity^81,89,91,93–95,97,98^. Many TFs predicted in inhibitory neurons (e.g.,*Tfap2b*, *Myt1l*, *Npas3, Pax2, Sox5, Etv6, Id2, Nfia, Nfix, Nfib, Maf, Pparg, Rora, Maf, Mef2c*) are known to be essential for the specification and maintenance of inhibitory interneurons within both central auditory regions and other areas of the brain^54,68,96,102,103^. We thus identified a combination of potential identity TFs, many of which have not been described before in CN, that may might act in concert to specify each CN cell type (Extended Data Table 8).

CN neurons exhibit remarkable wiring specificity, each receiving inputs from and extending output to specific pre- and post-synaptic neurons, respectively^4–6^, while cell-adhesion molecules (CAMs) are key determinants of neuronal identity and wiring specificity^54,104^. In line with these, GO terms containing the keyword ‘‘synaptic” and CAM gene families gave the highest score in distinguishing CN cell types (Fig.6a-b, Extended Data Table 5). We selected 342 genes encoding all major neuronal CAMs implicated in neural development and organized them into 10 CAM groups according to sequence homology and receptor-ligand relationships^54,105,106^, and nearly all groups are among the most distinguishing families (Fig.6d, Extended Data Table 5). Among these CAM genes, 135 show highly distinct cell type profiles (Extended Data Table 6). Strikingly, multiple CAM families each manifest differential expression among CN cell types (Fig.6f-g). For example, *Slit2/3* and their receptors (*Robo1*/*2*); netrin family (*Ntn4, Ntng1/2*) and their receptors (*Dcc*, *Neo1*); contactins and their binding partners; semaphorins and their receptors (plexins), exhibited highly differential expression, often with binary on/off patterns among five major projection neuron types (Fig.6f). These CAM families also exhibited highly differential expression among inhibitory neuronal cell types, often with distinct family members being employed from those in projection neurons (Fig.6g). These receptor-ligand pairs mediate axonal guidance and cell-cell recognition (either attractive or competitive/repulsive) during the neuronal developments, and thus might be key molecules for the establishment and maintenance of specific projection pattern of each CN cell type^54,107^. Each of the major synaptic CAM families including neurexin, neuroligin, protein tyrosine phosphatases (PTPR), leucine-rich repeat proteins (LRR), cadherin, neural IgCAM was also differentially expressed among CN cell types (Fig.6f-g). Cell-specific expression of these families, particularly LRRs, might contribute to post- and trans-synaptic specializations that customize the property of synapse types defined by each neuron and their specific target neuron^54,107–109^. In addition, two families of carbohydrate modifying enzymes (sulfotransferases and sialyltransferases) also exhibited highly distinct cell type profiles (Fig.6f-g, Extended Data Table 5), which might increase the molecular diversity of glycosylated CAMs and proteoglycans on the cell membrane and in extracellular matrix^110–112^. Together, cell- and synaptic CAM families likely constitute a multi-faceted cell-surface molecule code throughout the neuronal membrane of each CN neuron to determine their specific wiring.

### Molecular underpinnings of functional and biophysical specializations of CN neurons

We observed that the families of genes encoding ion channels are also highly predictive of CN cell types (Figs.6a-b, 7a, Extended Data Fig.13a, Extended Data Table 5). Two top families in this category encode potassium channels, while the gene families encoding sodium channels were not predictive of cell type identity (Figs.6a-b, Fig.7a, Extended Data Table 5). This suggests that potassium channels, rather than sodium channels, are the major determinants for the remarkable biophysical features and firing properties of CN cell types (Figs.2 and 5). To further explore this contention, and identify potential molecular determinants, we took advantage of the bimodal gene expression and physiological data in the Patch-seq dataset and performed sparse reduced-rank regression to identify gene transcripts that were predictive of firing properties of CN neurons^113,114^. To enhance interpretability, our analysis was restricted to voltage-sensitive ion channels within excitatory projection neurons (Methods, interneurons were excluded from the analysis due to insufficient data). These neurons exhibit either phasic firing (single-spike: bushy cells and octuple cells) or tonic firing (T-stellate and fusiform cells), which are differentiated by a set of highly correlated physiological features (Figs. 2b, 7b, Extended Data Fig. 12d-f and Table 2). Our sparse reduced-rank regression model selected 25 ion channel transcripts that achieved near-maximal predictive accuracy (Figs. 7b, Extended Data Fig. 12b-c). With additional statistical evaluations and the literature review, we identified two sets of transcripts among these 25 that are strongly predictive of the physiological hallmarks of projection neurons (Method, Fig. 7b and Extended Data Fig. 12c-f). One set of transcripts, including *Kcna1* (Kv1.1), *Hcn1* (HCN1) and *Kcnq4* (Kv7.4), were predictive of single-spike firing and associated features including low input resistance, short time constant, a high rheobase and threshold for spike generation, large voltage sag, and small AP amplitude (Fig. 7b and Extended Data Fig. 12c-d). This analysis suggested Kv1.1 and HCN1 as candidate molecular determinants of single-spike firing property, in line with prior evidence^35–37,115–117^, but also suggested another subthreshold-operating K^+^ conductance conferring single-spike firing property of CN neurons, Kv7.4-mediated conductance^118–121^. In contrast, a distinct set of transcripts including *Kcnb2* (Kv2.2) and *Kcnc2* (Kv3.2) were predictive of high-frequency tonic firing and associated features including large AHP, large AP amplitude, quick AP repolarization and narrow APs (Fig. 7b and Extended Data Fig. 12c, 12e), matching the role of these channels enabling repetitive high-frequency firing (by expediting AP repolarization and speeding the recovery of Na^+^ channels from inactivation)^39,42,48,122–124^. Among other relevant transcripts selected, two encoding potassium channels for transient A-type current, *Kcna4* (Kv1.4) and *Kcnd3* (Kv4.3), correlated with AP delay (Fig. 7b and Extended Data Fig. 12c, 12f), matching the transient nature of A-type current that mediates the delay to AP onset^125–127^. Thus, Kv1.4 could be a potential major channel mediating prominent A-currents in fusiform cells that determine their three distinct *in vivo* response patterns (“pauser”, “buildup”, and “chopper”)^126,128^.

**Fig. 7:**
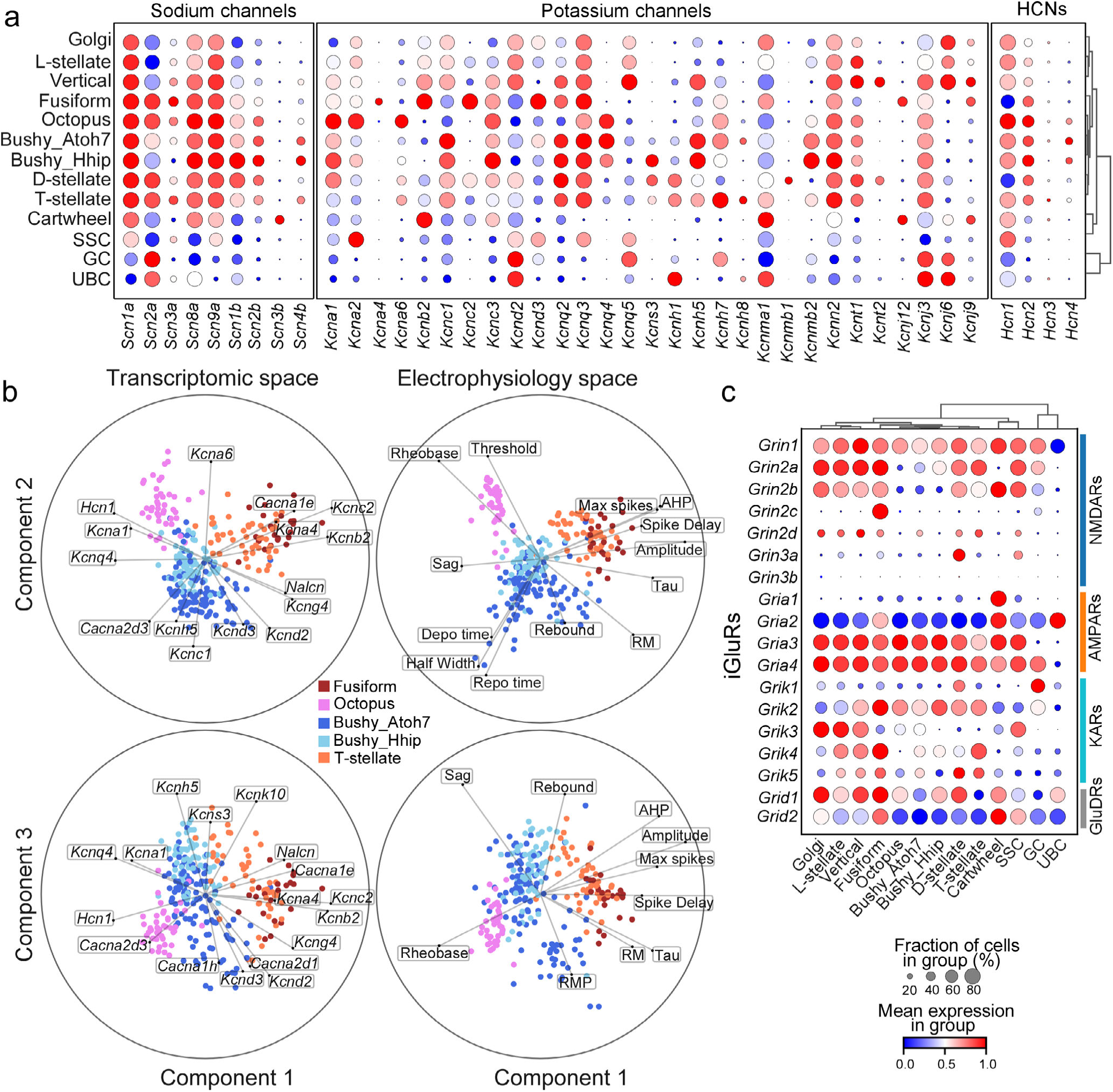
Single-cell transcriptomes provide insights into functional specializations of CN cell types. **a**, Dot plots showing scaled expression of the genes encoding ion channels across CN cell type (from snRNA-seq dataset). Sodium and potassium channel subtypes or subunits with low expression levels (< 20% fraction of the cell in any cell type) were excluded in the figure. Circle size depicts the percentage of cells in the cluster in which the marker was detected (≥ 1 UMI), and color depicts the average transcript count in expressing cells. **b**, Sparse reduced-rank regression (RRR) model to predict electrophysiological features by the expression patterns of 119 ion channel genes (middle, cross-validated *R*^2^ = 0.35). The models selected 25 genes. Cross-validated correlations between the first three pairs of projections were 0.83, 0.65, and 0.61. Both transcriptome (left) and electrophysiology (Right) were embedded in the latent space (bibiplots). In each biplot, lines represent correlations between a feature (gene expression or electrophysiology) and two latent components; the circle corresponds to the maximum attainable correlation (r =1). Only features with a correlation above 0.4 are shown. **c**, Dot plots showing scaled expression of genes encoding ionotropic glutamate receptors in each CN cell type. AMPARs: AMPA receptors, NMDARs: NMDA receptors, KARs: Kainate receptors. GluDRs: delta glutamate receptors. See Extended Data Fig. 12 for additional information.

In addition to biophysical specializations, CN neurons rely on specialized structural and synaptic features to fulfill their tasks^129,130^. Bushy cells, for example, require large presynaptic terminals (endbulbs of Held) with postsynaptic fast-gating AMPA receptors to enable precise spike timing with AN inputs^130^. These AMPA receptors are formed by GluR3 (*Gria3*) and GluR4 (*Gria4*), and are devoid of GluR2 (*Gria2*), thus conferring calcium permeability, large conductance and exceptionally rapid gating kinetics^131–133^. We therefore examined expression patterns of ionotropic receptors across cell types (Fig. 7c), and found that such exceptional AMPARs were also expressed in other CN cell types targeted by ANFs (i.e., T-stellate cells, octopus cells, vertical cells), but not in cell types targeted by non-ANF fibers, matching the observation that the very fastest synaptic currents are found only in neurons of the auditory system^129^. Thus, cartwheel cells, targeted by parallel fibers only, mainly expressed GluR1 (*Gria1*) and GluR2 (*Gria2*)^134^, an AMPAR composition with slow gating kinetics and calcium impermeability, while AMPARs in UBC, targeted by the mossy fibers, formed almost exclusively by GluR2 (*Gria2*, Fig. 7c), an uncommon form of native AMPAR^135^. Furthermore, fusiform cells, targeted by both ANF and parallel fibers, have mixed slow (GluR2) and fast AMPAR (GluR3/4) composition (Fig. 7c)^136,137^. This observation thus supports that postsynaptic AMPAR is specialized according to the source of synaptic input, likely illustrating a general rule governing synaptic specialization across the brain^134,136,137^. As for the other two types of ionotropic glutamate receptors (iGluRs; kainate and NMDA receptors), the expression of these relatively slow kinetic receptors was in general low in time-coding neurons (Fig. 7c), a necessary feature for fast high-fidelity transmission^138,139^.

CN neurons are subject to cell type-specific synaptic inhibition and neuromodulation to achieve or refine their functional specialization^140–142^. Our analysis indicates a wide variety of receptors for glycine, GABA, and neuromodulators in CN are cell-type specific (Extended Data Fig. 13b-c, Extended Data Table 5), in line with prior cellular and systems-level physiological observations. Particularly, a set of protein families in second messenger pathways (downstream of cell-surface receptors) including protein kinases, regulators of G-protein signaling, small guanosine triphosphatases (GTPase), and GTPase regulatory proteins (e.g., Rho-GEFs) are among the most differentially expressed gene families (Fig.6b, Extended Data Table 5). Differential expression of signaling proteins likely shapes the specificity and spatiotemporal dynamics of broadly acting second messengers, which might provide the mechanism and capacity to maintain the diversity of CN neuron morphology, synaptic connection, and neurite motility^143^. Finally, approximately half of the known deafness genes showed expression in single types or small sets of cell types, suggestive of their essential roles for central auditory processing (Extended Data Fig. 13d). Our dataset thus could provide a starting point to illustrate central components of hereditary deafness, and to differentiate poorly recognized central causes of deafness from peripheral causes (Extended Data Fig. 13d)^144,145^.

## Discussion

One of the biggest challenges in defining a cell-type taxonomy of the brain is to meaningfully reconcile definitions of cell types across the many modalities of measurement typically used to characterize brain cells^146^. Here in the CN, the first relay station in the auditory system where all high-order parallel processing pathways are initiated, we succeeded by first defining neuronal populations using unbiased single-nucleus RNA sequencing, and then relating these populations to anatomical, physiological, and morphological cell identities using Patch-seq combined with histology and FISH. We reveal a clear correspondence between molecularly defined cell populations and all previously defined cell types with known behavioral significance, as well as discover multiple new subtypes of the major projection neurons in the CN. We thus built a comprehensive and thoroughly validated atlas of cell types in the CN that harmonizes the cell type definitions across all modalities.

Neurons in the CN are highly specialized for processing different aspects of acoustic information and initiate distinct parallel pathways that encode sound location, intensity, frequency content, and spectrotemporal modulations^2,3^. Our analysis indicates that cellular and functional specializations for parallel auditory processing are bestowed by differential transcriptional programs dominated by a small set of gene families, and that these programs can be leveraged to define, discriminate, target, and manipulate the distinct processing pathways. Given the organizational similarities observed across different sensory systems^147^, we anticipate that this molecular design extends beyond auditory modality and applies to downstream auditory stations. Molecular definitions of cell types that initiate parallel pathways in sensory systems in general, and the auditory system in particular, open up new avenues for functional dissections of parallel processing. To date, no molecular tools are available to manipulate cell types in the CN to determine the impact on downstream sound processing and behavior. The novel molecular entry points and molecular basis of each cell type provided here ensure the rational design of a molecular toolbox (including the novel Cre lines and in vivo cell type-specific phototagging^148^) to link the activity of brainstem microcircuits to cortex population response patterns, and to perceptual learning or performance. We have validated an existing Penk-Cre line as a specific driver line for vertical cells^149^. Given their high cell type-specificity, development of Cre lines based on *Atoh7*, *Hhip*, *Necab2*, *Stac*, and *Phgdh* holds great potential for specifically targeting and manipulating SBC, GBC, fusiform cells, cartwheel cells, and octopus cells, respectively. Leveraging the extensive list of marker genes, transcription factors, CAMs, and ion channels with relatively specific expression patterns across cell types (Extended Data Table 1, Figs.6-7), intersectional approaches utilizing two or more of these genes could offer an alternative strategy for targeting and manipulating specific cell types. In addition, the cellular and transcriptomic atlas of the cochlear nucleus (CN) we generated offers an extensive resource and reference to investigate and identify hitherto unknown central components of hearing disorders^145^ and to illustrate central consequences of hearing loss and auditory trauma at a molecular level, potentially revealing the molecular underpinnings of neurological conditions such as tinnitus^149,150^. The atlas may also facilitate the rational design of auditory brainstem implants^150^.

### Identification of novel cell types in mouse CN

Our combined snRNA-seq/Patch-seq approach molecularly defined all previously described excitatory cell types, except for the sparse giant cells, which likely share the same molecular identity as fusiform cells^151,152^. This approach also revealed molecular subtypes within the classically defined cell types. By using transcriptomic clustering to parse electrophysiological and morphological datasets into different groups, we were able to identify subtypes of bushy cells and T-stellate cells with distinct properties. Two molecular subtypes of bushy cells have distinct anatomical preferences and response kinetics, reminiscent of GBC and SBC, a conventional category of bushy cells with distinct projection targets and functions^153–155^. We also found that within the groupings of *Atoh^+^* and *Hhip^+^* bushy cells, distinct variation exists in transcriptomic, topographic, and electrophysiological features, supporting the concept that SBCs and GBCs are broad cell types that each may contain distinct subtypes. More generally, using transcriptomic clusters to parse cells of similar electrophysiological features proved to be a powerful approach for extracting important biological meaning from the variance in a set of physiological measurements.

T-stellate cells play remarkably diverse roles in auditory function, having axonal projections within the CN and but also extending to multiple downstream nuclei as far as the auditory midbrain^6^. Such broad projections suggest diverse physiological functions, yet there has been little prior information to support subtypes of the T-stellate cell class. Our finding that T-stellate cells comprise two groups with strikingly distinct gene expression, intrinsic properties, and topography within the CN provides the first, clear evidence for subtypes of T-stellate cells. In vivo recordings refer to T-stellate cells as “choppers”, based on a frequency-independent, regular pattern of spikes observed in spike histograms. Interestingly, several classes of choppers have been distinguished in a wide variety of species including mice, with sustained and transient response patterns being the most common^156^. It is possible that the two T-stellate subtypes in the present study correspond to these different classes of chopper. Beyond morphological and electrophysiological properties, studies of these novel subtypes of bushy and T-stellate cells in terms of their synaptic inputs and axonal projections may reveal new parallel auditory pathways and new principles of auditory coding.

Until recently, the greatest diversity of inhibitory interneurons of CN was thought to be in the DCN^157^. The recent discovery of L-stellate cells in VCN indicated that VCN may also harbor multiple interneuron subtypes besides D-stellate cells^10^. Here, we show that L-stellate cells, defined simply as small glycinergic neurons of the VCN, are molecularly diverse. Those glycinergic neurons in the small cell cap resemble vertical cells in DCN (*Penk^+^*, purely glycinergic), differ from other L-stellate cells (dual transmitters, *Penk^-^*), and thus may comprise a novel type of inhibitory neurons in the VCN. Within the DCN *Penk* and *Stac* were selective markers for vertical and cartwheel cells, respectively. As these two cell types play distinct roles in gating auditory and multisensory signals in DCN, the identification and development of specific genetic tools based on their novel markers should facilitate future studies of their role in auditory processing. Another important novel interneuron marker is *Kit*, a gene specifically labeling those small glycinergic neurons across CN (Fig.5 and Extended Data Fig. 11) (i.e., excluding vertical cells, cartwheel cells, and D-stellate cells), and its mutations cause deafness suggestive of an important functional role in CN interneurons (Extended Data Fig. 13d)^158,159^. Guided by *Kit* expression, we observed small glycinergic neurons in the DCN that are neither vertical cells nor cartwheel cells, suggesting a hitherto unrecognized inhibitory cell type in the DCN. These glycinergic neurons in the DCN appear to be genetically similar to L-stellate cells in the VCN with dual transmitter phenotypes (Fig. 1C, Fig. 5, and Extended Data Fig. 11).

### Molecular underpinnings of cellular and functional specializations for auditory processing

Cell types with unique phenotypes are deeply rooted in the differential expression of many proteins. What types of genes distinguish cell types and also readily inform their phenotypes? TFs are widely thought to establish and maintain cell-type identity, while CAMs determine wiring specificity. Consistent with this contention, we found that TF and CAM gene families are among the most differentially expressed across CN cell types. The maintenance of cell identity involves the coordinated action of many TFs (‘combinatorial codes’). In addition, the expression of only a handful of key TFs (core TFs) is often sufficient to maintain the identity of a cell subpopulation^160,69–76^; depletion of these regulators causes significant alteration of cell identity, while forced expression of these regulators can effectively reprogram cells to a different cell type^161^. These TFs have been called ‘terminal selectors’ which activate or repress the cohort of effector genes that determine morphological, physiological, and molecular features specific for each cell type^57,59^. In the CN, while developmental programs governing early CN cellular differentiation are well defined^162,163^, the key TFs governing post-mitotic specification and maintenance of CN cell types remain largely unknown. Our analysis with bioinformatic tools predicts the potential candidate TFs for each CN cell type, offering novel targets for experimental validation. Furthermore, our dataset could serve as a valuable source to employ various approaches for identifying and exploring additional candidate TFs essential for CN cell type identity. Notably, most of these TFs have been shown to be expressed in different developmental stages of CN neurons^64,68,85,86^, suggesting that neuronal identities in adulthood are maintained by sustained expression of the same set of TFs^54,60^. Further investigation across different developmental stages may help reveal definitive TFs and regulatory programs that define each cell type. Ultimately, identifying these TFs will be critical for designing cell engineering strategies targeting distinct CN cell populations.

Synapses and neural circuits form with remarkable specificity during brain development ensuring the accurate flow of neural information. CN represents a ‘prototype’ sample; each CN neuron is organized tonotopically and receives the specific inputs from spinal ganglion neurons and projects downstream targets with high anatomical and target specificity^6^. Although this property of synapses and neural circuits in CN has been extensively investigated, molecular mechanisms underlying this property are largely unknown. Many studies suggest that the construction of the circuits, including central auditory circuits is at least partly defined by a combinatorial code of levels of cell-cell adhesion and axon guidance molecules^54,107,164–166^. The different types of CN neurons might assemble various types of CAMs and their receptors in time and space to mediate the selective contributions of diverse attractants and repellents from cells of their surrounding environment^166–168^. Our genome-wide mRNA sequencing data allows a comprehensive depiction of the molecular CAM framework dictating the anatomical/cell/synaptic-specific organization of CN circuits. Our analysis suggests such a molecular framework is encoded by a small set of CAM gene families, and combinatorial and coordinated expression of select members across these families together with their binding specificities contributes to precise neuronal connections and projection patterns. Many of these CAMs have been previously shown to be essential for the precise circuit wiring in the auditory system^167–172^ and in various other brain regions. A few seem specific to the auditory system, including Cntn5^173,174^, *Epha10*^175^, and *Carmil1* ^176^. Importantly, these CAMs are associated with various neurodevelopmental disorders including hearing loss (e.g., *Kirrel3, Ptprd, Dcc, Slit2/3, Robo1, Ntng1/2, Cntn5, Cntn6, Cdh13, Dmd, Utrin, Epha10, Carmil1*)^175–185^, dyslexia (e.g., *Cntnap5a, Robo1*)^186,187^, and speech sound language disorders (*Robo1)*^181^, autism spectrum disorder (ASD) and intellectual disability (ID) with hearing defect as an essential component, constituting novel candidate molecules for experimental follow-up.

Our transcriptomic dataset offers a comprehensive resource to unravel molecular underpinnings of morphological, synaptic, and biophysical specializations required for CN neurons to encode auditory signals. Our analysis suggests that the remarkable biophysical specialization and diversity of CN neurons is bestowed by a small set of ion channel families, particularly potassium channels. Notably, we predicted three transcripts including *Kcna1*, *Hcn1,* and *Kcnq4* as potential molecular determinants for single-spike firing property, a property required for time-coding neurons to encode precise AP timing for binaural sound detection^188^, in line with the observations that mutations of these genes impair binaural hearing^189–192^ (Extended Data Fig. 13d). While Kv7.4 currents in time-coding neurons were previously undescribed despite strong *Kcnq4* expression in CN, their role in enabling single-spike firing in concert with Kv1 is well-established in many other cell types^118–121^, and *KCNQ4* mutations causes profound hearing loss in human DFNA2 that could not be solely explained by hair cell dysfunctions^193,194^. Another important ion channel for CN projection neurons is high-threshold Kv3, and these neurons appear to employ distinct subunits of this family to fulfill their unique biophysical needs. While Kv3.2 (*Kcnc2*) seems crucial for the tonic firing of T-stellate and fusiform cells (Fig.7b), Kv3.3 transcript (*Kcnc3*) is specifically enriched in time-coding neurons (Fig. 7a) to possibly mediate their fast spike repolarization, a necessary feature for the use of both temporal and rate coding in acoustic processing^39,40^. This observation may explain why SCA13 patients (with *KCNC3* mutations) exhibit impaired interaural timing and intensity discrimination^195^ (Extended Data Fig. 13d). Interestingly, our analysis did not support a significant contribution of sodium channel diversity to remarkable biophysical properties of CN projection neurons (Extended Data Table 5)^196^. These neurons exhibited similar expression patterns of sodium channel α subunits, primarily *Scn1a*, *Scn2a*, *Scn8a*, and unexpectedly, *Scn9a* (Fig. 7a), a subunit previously thought to be restricted to the peripheral nervous system^197^. However, these genes were more differentially expressed among CN inhibitory neuron cell types (Fig. 7a), suggesting a greater role for sodium channels in shaping the diverse firing properties of CN interneurons. Further investigation is warranted to fully elucidate the contribution of ion channel diversity, including both potassium and sodium channels, to the biophysical properties of CN interneurons.

## Methods

### Animals

All experiments were performed according to the guidelines of the Institutional Animal Care and Use Committee (IACUC) of Baylor College of Medicine and Oregon Health and Science University. Wild-type mice or transgenic mice of both sexes on the C57/BL6/J background at the age of postnatal (P) 22-28 were used for single-nucleus RNA sequencing and slice electrophysiology (including Patch-seq). To help distribute our sampling across cell types and to increase the chance of recording certain specific cell types, GlyT2-EGFP (RRID:MGI:3835459) mouse line was used to target glycinergic/inhibitory neuronal populations (labeled neurons), while unlabeled cells in this line were targeted for excitatory neuron recording^10^. SST-Cre (RRID:IMSR_JAX:013044) cross with Ai9 (RRID:IMSR_JAX:007909) mouse line was crossed with GlyT2-EGFP mice to identify D-stellate cells and L-stellate cells in the CN for patch clamp recordings^10^. Some recordings from excitatory neurons were performed using wild-type mice as well. In this study, we used 222 mice in total (121 males and 101 females), including 132 wild-type mice, 61 GlyT2-EGFP mice, 17 Penk-Cre:Ai9 mice, and 3 Gabra6-Cre (RRID:IMSR_JAX:007909):Ai9 mice. Penk-Cre (RRID:IMSR_JAX:025112) mice are a gift from Yong Xu Lab at Baylor College of Medicine^149^, while Gabra6-Cre mice are a gift from Susumu Tomita Lab at Yale University. All animals were maintained in the animal facility with a light cycle from 6 am to 6 pm daily, with temperatures ranging from 68 to 72 °F and humidity ranging from 30% to 70%.

### Single nucleus extraction and single-nucleus RNA sequencing (snRNA-seq)

Mice were deeply anesthetized with 3% isoflurane and decapitated immediately, and their brains were immediately removed from the skull and then transferred into an iced oxygenated NMDG solution (93 mM NMDG, 93 mM HCl, 2.5 mM KCl, 1.2 mM NaH_2_PO_4_, 30 mM NaHCO_3_, 20 mM HEPES, 25 mM Glucose, 5 mM Sodium Ascorbate, 2 mM Thiourea, 3 mM Sodium Pyruvate, 10mM MgSO_4_ and 0.5 mM CaCl_2_, pH 7.4) for further dissection. Cochlear nuclei (CNs) were dissected from each brain under a stereoscopic microscope and then transferred into a 1.5 mL Eppendorf tube, and tubes were immediately immersed in liquid nitrogen. Brain tissues were then transferred into a −80 °C freezer for long-term storage. Each tissue sample was pooled from 8 to 10 mice for single-nucleus extraction and single-nucleus RNA sequencing (snRNA-seq).

Single nucleus extraction from brain tissues followed a published protocol^198^. Briefly, frozen tissues were homogenized with 2 mL homogenization buffer (HB, 250 mM Sucrose, 25 mM KCl, 5 mM MgCl_2_, 10 mM Tris, 1mM DTT, 1x Protease Inhibitor, 0.4 units/µL Rnase Inhibitor, 0.1% Triton X-100) in Wheaton Dounce Tissue Grinder. Cell debris and large clumps were removed with a 40 µm cell strainer (Flowmi). The homogenate was centrifuged at 1,000 g for 5 min at 4 °C. The palette was resuspended with 25% iodixanol in the centrifugation buffer (CB: 250 mM sucrose, 25 mM KCl, 5 mM MgCl_2_, 10 mM Tris) and layered on top of 29% iodixanol. The mixture was centrifuged at 13,500 g for 15-30 min at 4 °C. The supernatant was carefully removed without disrupting the nuclei pellet. The nuclei pellet was resuspended with the resuspension buffer (RB: 1 % UltraPure BSA in Rnase-free PBS).

Nuclei were counted using a hemocytometer or CellCounter (Countess II, Invitrogen) and diluted to ∼ 1,000 nuclei/µL for single-nucleus capture on the 10x Genomics Chromium Next GEM Single Cell 3’v3 system following the standard user guide. Droplet-based snRNA-seq libraries were prepared using the 10x Genomics Chromium Single Cell kit according to the manufacturer’s protocol and were sequenced on the Illumina NovaSeq 6000 or Hiseq 4000 platform with a pair-end 150 bp strategy (average depth 40k–50k reads/nucleus). Cell Ranger (version 5.0.1) with the *Mus musculus* genome (GRCm38) and annotation GTF (version M23) was used to generate the output count matrix.

### Slice preparation and electrophysiology

Brain slices were prepared from the mouse CN as previously described with a slight modification^199–201^. Briefly, P22-28 mice were deeply anesthetized with 3% isoflurane and then immediately decapitated. The brain was removed and placed in the iced oxygenated NMDG solution (for the recipe, see above). The brain tissue containing CN was sliced into 200-300 µm thick sections parasagittally in the NMDG solution with a microtome (Leica, VT1200). The slices were transferred into the oxygenated NMDG solution (34 ± 0.5 °C) for 10 min and then incubated in the artificial cerebrospinal fluid (ACSF, 119 mM NaCl, 2.5 mM KCl, 1 mM NaH_2_PO_4_, 25 mM NaHCO_3_, 1 mM MgCl_2_, 25 mM glucose and 2 mM CaCl_2_, pH 7.4) at 34 ± 0.5 °C for 40-60 mins before recording.

Whole-cell recordings were obtained from CN neurons as previously described ^201^. Borosilicate glass pipettes (3–5 MΩ) were pulled with micropipette pullers (P-1000, Sutter) and filled with intracellular solution containing 120 mM potassium gluconate, 10 mM HEPES, 4 mM KCl, 4 mM MgATP, 0.3 mM Na_3_GTP, 10 mM sodium phosphocreatine, and 0.5% biocytin (pH 7.25). Whole-cell recordings were performed at 32 ± 0.5 °C with the EPC 10 amplifiers (HEKA Electronics, Lambrecht, Germany). PatchMaster (HEKA) was used to operate the recording system and record the electrophysiology data. The membrane potential response of each neuron to increasing current steps (600 ms) every 1s was digitized at 25 kHz. To compare the intrinsic properties and firing pattern across cell types, their membrane responses to a hyperpolarizing current step, to the current step at the rheobase, and to a current step at 2X rheobase were recorded and shown for each cell type (Fig. 2-5). Recorded neurons were then fixed for *post hoc* morphological recovery.

### Patch-seq

Patch-seq was performed on CN neurons following our published protocol^202^. Briefly, the RNA was extracted from each neuron upon completion of whole-cell recordings (including recordings of synaptic events) with a modified internal solution (111 mM potassium gluconate, 4 mM KCl, 10 mM HEPES, 0.2 mM EGTA, 4 MgATP, 0.3 Na_3_GTP, 5 mM sodium phosphocreatine, 0.5% biocytin and 0.48 units/µL RNase inhibitor, pH 7.25). Recorded neurons were then fixed for *post hoc* morphological recovery. The RNA was reverse transcribed with Superscript II Reverse Transcriptase (Invitrogen). The cDNA samples that passed quality control were used to generate sequencing libraries using the Nextera XT DNA Library Preparation Kit (Illumina, FC-131-1096), following the user guide. Each library was sequenced at a depth of 1∼2 million pair-end 150bp reads on the Illumina NovaSeq 6000 or Hiseq 4000 platform. Raw reads were aligned to the *Mus musculus* genome (GRCm38) with annotation GTF (version M23) using STAR 2.7.7a^203^, and an expression matrix was generated with FeatureCounts^204^.

### RNA fluorescence in situ hybridization

RNA fluorescence in situ hybridization (FISH) was performed on 25 µm thick sagittal sections containing the CN at the RNA In Situ Hybridization Core at Baylor College of Medicine, using a previously described in situ hybridization method with slight modifications^205^. Digoxigenin (DIG) and Fluorescein isothiocyanate (FITC) labeled mRNA antisense probes were generated from reverse-transcribed mouse cDNA. Primer sequences for the probes were derived from Allen Brain Atlas (http://www.brain-map.org). DIG-labeled probes were visualized with tyramide-Cy3 Plus and FITC-labeled probes were visualized with tyramide-FITC Plus. Images were captured at 20X magnification using LSM 710 or 880 confocal microscope (Zeiss).

### Immunofluorescence staining

GlyT2-EGFP mice at P22-28 were perfused transcardially with 20 mL PBS followed by 20 mL of 4% PFA in 0.1 M PB. The brain was removed, post-fixed in 4% PFA for 1 day, and then sagittally sectioned into 50 µm thick slices in PBS. The slices were washed 3 times in PBS before blocking with 10% goat serum and 1% Triton X in PBS for 1 hour. The slices were incubated first with mouse anti-PHGDH (Invitrogen, PA5-54360, 1:400) or rabbit anti-NECAB2 (Sigma-Aldrich, HPA013998, 1:500) as primary antibodies, and then incubated with Alexa-Fluor goat anti-mouse 633 (Invitrogen, A-21052) or goat anti-rabbit Alexa-Fluor 647 (Invitrogen, A-21244) as secondary antibodies for at least 1 hour.

Slices were then mounted with the mounting medium (Abcam, ab104139). To stain PHGDH or NECAB2 in recorded neurons, brain slices were immediately fixed after whole-cell recording with 4% PFA for 1 day. The slices were first incubated with primary antibodies (see above) for at least 1 hour, and then incubated with Alexa Fluor 568 conjugates of streptavidin (Invitrogen, S11226) to visualize biocytin together with those secondary antibodies (see above) to visualize putative signals of PHGDH or NECAB2 in recorded neurons. Images were captured at 10X or 20X magnification using LSM 710 confocal microscope (Zeiss).

### Morphological recovery

Morphological recovery of recorded neurons and light microscopic examination of their morphology were carried out following previously described methods^201^. In brief, upon the completion of whole-cell recordings, the slices were immediately fixed in 2.5% glutaraldehyde/4% paraformaldehyde in 0.1M PB at 4 °C for 10-14 days and then processed with an avidin-biotin-peroxidase method with an ABC kit (Vector Laboratories, PK-4000) to reveal cell morphology. Recovered neurons were examined and reconstructed using a 100X oil-immersion objective lens and camera lucida system (Neurolucida, MicroBrightField). For morphological feature extraction, only cells with intact dendritic structures were included. Axons were excluded from this assessment as they are typically severed during slice preparation. We took special care to exclude reconstructions of the neurons that showed any signs of dendritic damage (due to the slicing procedure), or poor overall staining due to the aspiration procedure involved in Patch-seq (invisible or unclear soma and dendrite, likely caused by the collapse of the soma during nucleus extraction and subsequent leakage of the biocytin).

### Computational analysis of snRNA-seq data

For snRNA-seq data, we used CellRanger (version 5.0.1) to generate a barcode table, gene table, and gene expression matrix. Downstream analysis and visualization were performed using Scvi-tools and SCANPY^206^. For preprocessing, we followed the standard protocols used in the field^207^. Specifically, nuclei with > 5% UMIs mapped to mitochondrial genes and those with less than 200 genes detected were excluded from the analysis. We removed doublets sample-by-sample using Scrublet^208^. Doublet score and doublet predictions histogram of each sample were manually inspected, adjusted, and then concatenated. A putative single nucleus with a doublet score exceeding the 95^th^ percentile was removed from the downstream analysis. Those nuclei co-expressing multiple canonical marker genes for different cell types as follows were also considered as doublets and were excluded for further analysis: *Mobp/Mog/Mag* (ODC), *Gabra6* (granule cell), *Ptprc/Csf1a/C1qa* (microglia), *Slc1a2/Aqp4/Gja1* (astrocyte), *Dcn* (fibroblast), *Gad1/Gad2/Slc6a5* (inhibitory neurons), *Flt1/Cldn5* (endothelial). Counts of all the nuclei were normalized by the library size multiplied by 10,000 and then logarithmized. The top 6,000 highly variable genes, which were identified by dispersion-based methods, were used for principal component analysis. Leiden graph-clustering method and UMAP were used for clustering and visualization.

Due to the proximity of the CN to the cerebellum (CB), paramedian lobule (dorsal to the CN), to the principal sensory nucleus of the trigeminal nerve (PSN; ventral to CN), and to the dentate nucleus, our microdissection inevitably includes more or less of these adjoining tissues. To remove potential CB-derived nuclei from our dataset (Extended Data Fig. 2), we compared our data to a recently published CB snRNA-seq dataset^21^. By examining the expression of marker genes for each CB cell type in our snRNA-seq data, we identified several putative neuronal and non-neuronal CB clusters in our dataset. Firstly, two clusters show clear molecular signatures of Bergmann cells and Purkinje cells respectively (Extended Data Fig. 2a-b), indicating they are contaminated nuclei from these two CB-specific cell types. Secondly, two large clusters assigned to granule cells (i.e., enriched with the canonical marker gene *Gabra6*) could be discriminated by the expression levels of *Grm4* and *Col27a1* (Extended Data Fig. 2c). Based on the molecular signatures of CB granule cells^21^ and Allen Brain Atlas ISH data (Extended Data Fig. 2c), we concluded that *Grm4^-^*/*Col27a1^+^* cluster corresponds to granule cells from CN, while the *Grm4^+^*/*Col27a1^-^*cluster corresponds to granule cells from CB (CB_Granule). After removing these CB clusters from our dataset, we computationally isolated the neuronal population (except for granule cells and UBC) and then performed subclustering analysis to reveal distinct neuronal clusters (Extended Data Fig. 2d). One neuronal cluster enriched with *Htr2c* and two clusters enriched with *Necab2* were first identified (Extended Data Fig. 2e-f). As shown on Allen Brain Atlas (Extended Data Fig. 2e), *Htr2c* expression is restricted to the dentate nucleus, and thus this *Htr2c-*enriched cluster in our dataset is from the dentate nucleus. Two *Necab2^+^* clusters are either *Tnnt1*^+^ or *Tnnt1*^-^, and only the *Tnnt1*^+^ nuclei are from the CN while *Tnnt1*^-^ nuclei are from the PSN, based on Allen ISH data (Extended Data Fig. 2e-f) and our FISH data (Extended Data Fig. 7b5).

We then identified two pure GABAergic clusters (*Gad1*^+^/*Slc6a5^-^)* enriched with *Tfap2b* (Extended Data Fig. 2g), reminiscent of MLI1 and MLI2 from the CB^21^. Since pure GABAergic interneurons in the CN are absent or extremely rare (Extended Data Fig. 2h), we concluded these two clusters are indeed MLI1 and MLI2 from the CB. We also identified two *Tfap2b*-enriched glycine/GABA combinatorial clusters reminiscent of Golgi cells^21^, which were either *Scrg1^+^* or *Scrg1^-^*. We concluded that the *Scrg1^+^* cluster results from contaminated nuclei from Golgi cells in the CB (CB_Golgi) based on the CB snRNA-seq dataset^21^ and our FISH data (Extended Data Fig. 2i); only the *Scrg1^-^* cluster corresponds to CN Golgi cells (CN_Golgi, Extended Data Figs. 2i and 5). We also identified a glycine/GABA combinatorial cluster enriched with both *Klhl1 and Slc36a2* (Extended Data Fig. 2g), reminiscent of Purkinje layer interneurons (PLIs) from the CB^21^. We conclude that this indeed corresponds to PLIs contaminated from the CB based on the expression patterns of *Klhl1 and Slc36a2* across CN and CB (Extended Data Fig. 2j).

The potential inclusion of non-neuronal nuclei from adjacent regions (with the exception of Bergmann glia) in our dataset was not addressed, since non-neuronal populations are not the focus of our analysis. Thus, the non-neuronal nuclei as shown in Extended Data Fig. 1b may include those from adjoining regions. The UBC population in our dataset may include CB-derived UBCs, but we believed that this contamination, if any, was minimal. Firstly, using the ratio of the UBC population versus GC population in the CB (∼1: 300)^21^, we estimated ∼80 UBCs from CB might be included in our dataset (based on 24,118 CB-derived GCs in our dataset). In addition, the region of the CB close to the CN (i.e., CB paramedian lobule) has a smaller population of UBCs compared to other CB regions. We thus concluded that out of 3,653 UBCs in our dataset, only a very small number may be UBCs from the CB, which would not significantly affect our analysis.

After the removal of the nuclei from adjoining tissues, the remaining 61,211 nuclei were used for further clustering analysis, and each cluster and cell type were annotated with the commonly used marker genes as listed in Extended Data Table 1 (including the marker genes for CB cell types). Calculation of the local inverse Simpson’s index^209^ demonstrated that these populations were well mixed and contained cells from each sample, indicating successful integration (Extended Data Fig. 3). Our initial clustering combined the single-nucleus expression data from male (n = 31,115 nuclei) and female samples (n = 13,140 nuclei); each cluster comprised cells from both sexes, indicating that there were few sex-dependent differences (Extended Data Fig. 3). The data from both sexes were thus pooled for subsequent analyses.

### Patch-seq data Analysis

In total, we performed Patch-seq from 438 cells (323 excitatory neurons, 115 inhibitory neurons, from 69 mice) that passed initial quality control. Among these sequenced 438 cells, 387 neurons (293 excitatory neurons, 94 inhibitory neurons) passed further quality control, yielding 2.40 ± 0.08 million (mean ± s.e.) reads and 8,025 ± 70 (mean ± s.e.) detected genes per cell (Extended Data Fig. 3). Patch-seq data were clustered and visualized following a similar method with SCANPY. Cells with a mapping rate of < 60% and gene features of < 4000 were not included in the analysis. We used the batch-balanced KNN method for batch correction^210^. Patch-seq data were mapped to the snRNA-seq data following a similar protocol as used previously^28,114^. Briefly, using the count matrix of 10x Genomics snRNA-seq data, we selected 3000 “most variable” genes, as described previously^211^. We then log-transformed all counts with log2(x+1) transformation and averaged the log-transformed counts across all cells in each of the identified clusters in the snRNA-seq dataset to obtain reference transcriptomic profiles of each cluster (N×3000 matrix). Out of these 3,000 genes, 2,575 are present in the mm10 reference genome that we used to align reads in our Patch-seq data. We applied the same log2(x+1) transformation to the read counts of Patch-seq cells, and for each cell, computed Pearson correlation across the 2,575 genes with all clusters. Each Patch-seq cell was assigned to a cluster to which it had the highest correlation. To illustrate the mapping results, each Patch-seq cell was positioned at the median UMAP coordinates of its ten nearest neighbors in snRNA-seq UMAP embeddings, as determined by Pearson correlation^28,114^.

### DEG analysis and marker gene identification

To identify the DEGs for clusters and subclusters, we used SCANPY to rank genes using the T-test_overestim_var test with Benjamini–Hochberg correction. SCANPY generated logFC (with the base as 2) and FDR (False discovery rate) corrected P value and DEGs were identified based on p-value < 0.01 and |log2FC| > 1. This method could detect more DEGs in the clusters that have large numbers of cells, while in relatively small cell populations only those with much higher fold changes could be identified as DEGs.

### Electrophysiological feature extraction and analysis

All cells that passed the electrophysiology data quality were included in the analysis. Electrophysiological data were analyzed with a python-based GUI named PatchView that we have developed^212^ or a custom-made IDL code. To access the IDL code, the raw electrophysiological data was exported to ASCII format using Nest-o-Patch. To highlight the differences in the intrinsic properties and firing patterns, the membrane responses to a hyperpolarizing current step, to the current step at the rheobase, and to a current step at 2X rheobase were shown for each cell type (Fig. 2-5). 20 electrophysiological features were extracted from the electrophysiological traces following a similar method as previously described (Extended Data Fig. 14)^114^.

To calculate the time constant (Tau), all the traces in response to negative current steps (from the onset of the current step to the minimal potential during the stimulation period) were fitted with an exponential function as follows:

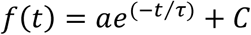

Here, *f(t)* is the voltage at the time *t*, τ is the membrane time constant, *a* and *C* are two constants. The 𝜏 was calculated for each trace. Boxplot was used to remove the outliers and the mean averaged across all traces after outlier removal was used as the time constant.

We calculated the sag ratio and rebound voltage using the hyperpolarizing trace elicited by the maximal negative current injection. The sag ratio was defined as the difference between the sag trough of the trace and the resting membrane potential (RMP) (ΔV1), divided by the difference between steady-state membrane voltage and the RMP (ΔV2). The trace from the stimulation offset to the peak of the depolarizing trace following the offset was fitted with an exponential function, and the potential difference between the maximal value of the fitting curve and the RMP was defined as the rebound value.

To calculate the input resistance and RMP, we measured the steady-state membrane potentials (determined by the average values of the last 100 ms of the hyperpolarization) in the traces elicited by the first five negative-going currents in the step protocol, and then plotted these values (y-axis) with each of the corresponding currents (x-axis). We fit the plot with a linear function shown as follows:

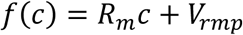

Here, *f(c)* steady-state voltages, c is the injected currents, *Rm* and *Vrmp* are constants that represent the slope of the linear function and the potential at the y-axis intersection, corresponding to the input resistance and the RMP respectively.

Rheobase is defined as the minimum current that could elicit any action potentials (APs) during the step-current injection. To extract the properties of action potential (AP), we used the first AP elicited by our step stimulation protocol. To calculate the threshold of the AP, the trace within 20 ms preceding the AP peak was used to calculate the third derivative and the data point with the maximal third derivative value was defined as the AP threshold^213^. The AP amplitude is the difference between the threshold potential and the peak potential of the AP. The AP half-width was defined as the time difference between the voltage at AP half-height on the rising and falling phase of the AP. The afterhyperpolarization (AHP) was defined as the potential difference between the AP threshold and the minimal value of the trough after the AP. The time difference from the stimulation onset to the first AP’s threshold at the rheobase sweep was defined as the AP delay time. The depolarization time was the time difference from the AP threshold to the overshoot. The time from the overshoot to the repolarizing voltage that equaled the threshold was the AP’s repolarization time. Spike-frequency adaptation was quantified with the membrane potential trace in response to the injection of the step current at 2X Rheobase. The interspike interval (ISI) between the first two APs divided by the ISI between the second and the third APs was the initial adaptation index. The ISI between the first two APs divided by the ISI between the last two APs was the last adaptation index.

The features used for E-type clustering analysis and UMAP embedding calculation were shown in Extended Data Fig. 4. The features were standardized using StandardScaler from scikit-learn Python library (version 1.3.2). UMAP embeddings were generated from the standardized electrophysiological features using the umap-learn Python library (version 0.5.3) (Fig. 2c, Fig. 5c, Extended Data Fig. 4b, f, h). K-means clustering analysis was performed on standardized electrophysiological features (using KMeans from scikit-learn library) to identify inherent groupings within the dataset. The results of this analysis were visualized on UMAP embeddings (Extended Data Fig. 4b, h). A confusion matrix was used to show the performance of K-means clustering in differentiating cell types as labeled manually (Extended Data Fig. 4b and h). To include the recording location (either VCN or DCN) as an additional feature in the inhibitory neuron clustering analysis, inhibitory neurons recorded in the VCN were assigned a value of 0, while those recorded in the DCN were assigned a value of 1. These values were then incorporated alongside the electrophysiological values for K-mean clustering analysis and to generate UMAP embedding (Extended Data Fig. 4h and Fig.5d). Different value assignments were tested and did not impact clustering analysis.

To determine whether tdTomato^+^ cells in Penk-Cre mice aligned with the electrophysiological profile of vertical cells (Fig. 5l), we extracted the electrophysiological features of those tdTomato^+^ cells recorded from Penk-Cre mice, and these features were then used to cluster tdTomato^+^ cells together with all other recorded inhibitory neurons (Fig. 5l). Visualizing tdTomato^+^ cells within the resulting UMAP embeddings revealed their relationship (co-clustering) with the population of recorded vertical neurons (Fig. 5l).

### Morphological reconstruction and analysis

All cells that passed the morphological criteria were reconstructed and included in the analysis. Neuronal morphologies were reconstructed with Neurolucida (MicroBrightField) and analyzed with Neurolucida Explorer (including Sholl analysis of the dendritic arbors and the soma shape analysis). For a more comprehensive morphological feature analysis, each morphology file was converted into the SWC format with NLMorphologyConverter 0.9.0 (http://neuronland.org) for feature extraction using MorphoPy (https://github.com/berenslab/MorphoPy). See https://github.com/bcmjianglab/ cn_project/tree/main/patchseq/M-clusters for a detailed description of the morphological features.

The features used for M-type clustering analysis and UMAP embedding calculation are shown in Extended Data Fig. 4. UMAP embeddings were generated from standardized morphological features using the umap-learn Python library (version 0.5.3) (Fig. 2d, Fig. 5c, Extended Data Fig. 4b, d). K-means clustering analysis was performed on these features as did on electrophysiological features and the results were visualized on UMAP embeddings (Extended Data Fig. 4b, d). The confusion matrix was used to show the performance of K-means clustering (Extended Data Fig. 4b, d).

To determine whether tdTomato^+^ cells in Penk-Cre mice aligned with the morphological profile of vertical cells (Fig. 5l), we extracted the morphological features of those tdTomato^+^ cells recorded from Penk-Cre mice, and these features were then used to cluster tdTomato^+^ cells together with all other recorded inhibitory neurons (Fig. 5l). Visualizing tdTomato^+^ cells within the resulting UMAP embeddings revealed their relationship (co-clustering) with the population of recorded vertical neurons (Fig. 5l).

### Sparse reduced-rank regression, and simple linear correlation and regression

Sparse reduced-rank regression (RRR) was performed as previously reported^114^. In brief, we used 14 electrophysiological features extracted from 293 Patch-seq excitatory neurons (Fig. 2e) to predict the electrophysiological features from gene expression. Interneurons were excluded from the analysis due to data insufficiency. To enhance interpretability, we only focused on voltage-sensitive ion channel genes and 119 genes out of 330 genes in this category (https://www.genenames.org/) were chosen for the analysis (those expressed in less than 10 of the Patch-seq neurons were excluded). We optimized the hyperparameter combinations based on the model’s predictive performance (measured by cross-validated R-squared), and chose a model with rank = 5, zero ridge penalty (α = 1), and regularization strength (λ = 0.28) tuned to yield a selection of 25 genes, which achieved near-maximal predictive accuracy (Extended Data Fig.12b). Two biplots (bi-biplots) were used to show the RRR analysis results^114^, where the directions and magnitudes of lines representing individual genes and electrophysiological features reflect their correlations to two underlying components. Only features with a correlation above 0.4 are shown in the bibiplots. By comparing these directions and magnitudes between the two biplots, potential associations between specific genes and electrophysiological traits can be inferred.

To provide an additional, more direct assessment of the relationship between each ion channel gene and the electrophysiological features, the simple linear correlation and regression (including effect size and significance) between each of ion channel genes (n=119, including 25 gene selected by the sparse RRR model) and the electrophysiological features were independently calculated (Extended Data Fig. 12c). The correlation coefficients, significance and linear regression lines are visualized in dot plots and scaled plots (Extended Data Fig. 12c-f). For all ion channel genes, only the top 25 with the highest correlation coefficients are displayed, and most of these overlap with the genes selected by the sparse RRR model. This substantial overlap serves as compelling validation for the accuracy and effectiveness of the sparse RRR model.

### Registering the anatomic location of cells

To register each Patch-seq cell to the CN 3-dimensional (3D) location, we first generated a 3D CN reference space based on the Allen Mouse Brain Atlas and then aligned each brain slice used for our Patch-seq recordings to this reference space. We reconstructed a contour on each slice that delineates the entire CN area in that slice, and then iteratively searched for the contour within the 3D CN reference space that best matched the reconstructed contour (i.e., estimating the 3D location and orientation of that slice). Since our slices were cut in the para-sagittal plane, we limited our grid search parameters range: pitch angle (0, −25 radians) and roll angle (0, −20 radians), median to lateral shift (0, 850 µm), in-plane orientation (−90, 90). For each parameter set, we calculated the overlapping area of the reconstructed contour with the corresponding contour in the 3D CN reference space. The parameters that result in a maximal overlapping area were used as the final parameter for estimation.

We visually inspected the mapping results to rule out erroneous registration. For some ambiguous slices (i.e., having very small CN regions), we further limited the search parameter range to improve the accuracy. The location of neurons in that slice was then projected into the 3D CN reference space.

### Machine learning algorithms to validate expert classification

Expert classification was performed by four individuals (TN, LT, JJ, MM), all with extensive expertise in studying CN cell types. Each individual independently assessed each cell based on its location, morphology, input resistance, and responses to a family of depolarizing and hyperpolarizing current steps. The results of these independent evaluations were then compared to established features in the CN literature^4–6^. Any disagreements were resolved through discussions in group meetings until a consensus identification was reached for each recorded cell. A random forest classifier based on electrophysiological or morphological features was then used to validate expert classification. Briefly, the electrophysiological or morphological features of each neuron were extracted, normalized, and scaled. To include the recording location (either VCN or DCN) within electrophysiological features during the classifier training process (for inhibitory neuron classification), inhibitory neurons recorded in the VCN were assigned a value of 0, while those recorded in the DCN were assigned a value of 1 (Extended Data Fig. 4g). Different value assignments were tested and did not impact classifier performance. The data were split into training and testing datasets randomly, and the classifier was trained with the training data. The accuracy in distinguishing the cell types as labeled manually by the classifier was evaluated with the testing dataset. The accuracy varied with different training datasets used, so we trained the classifier 10,000 times, each with a different training dataset. The classifier settings with the top 100 performance outcomes were used to weigh the electrophysiological or morphological features for cell type discrimination. The overall accuracy in discriminating cell types by the classifier was reported as the mean accuracy within the chosen settings. The top 15 most important features contributing to the performance were used for UMAP visualizations (Extended Data Fig. 4a, 4c, 4e, 4g). A confusion matrix was used to show the overall accuracy in distinguishing the cell types as labeled manually by the classifier (Fig. 2c-d; Fig.5c-d, Extended Data Fig. 4f).

### MetaNeighbor analysis and gene expression specificity analysis

To identify the gene families or sets whose differential expression could robustly predict cell types, MetaNeighbor analysis was conducted on our snRNA-seq dataset (either the whole dataset, projection neurons or inhibitory neurons only) utilizing pyMN tool, following established user tutorials^214^. In brief, we split our dataset into two halves and aimed to measure the replicability of cell type identity between them when considering only a given gene set or family. Cell type identities were held back from one set at a time and predicted from the other by neighbor voting based on a Spearman correlation subnetwork built from the expression of the gene set (or family) being tested. The accuracy of recovering cells of the same cell types was evaluated through cross-validation and reported as AUROC (Area Under Receiver-Operator Characteristics Curve) values. We then ranked the gene sets or families by the mean AUROC value. We first performed this analysis on gene sets annotated with mouse Gene Ontology (GO) terms (obtained via pyMN). To identify more specific gene categories, we also applied the test to all 620 gene families annotated in the HUGO Gene Nomenclature Committee (HGNC) database (https://www.genenames.org/cgi-bin/genegroup/download-all). Furthermore, we curated 342 cell adhesion molecule (CAM) genes or CAM-modifying enzyme genes into 12 families (Extended Data Table 5), following the strategy by Paul A. et al., 2017^54^. We conducted additional tests on these curated families. Supplementary tests and AUROC scoring were also performed on 70 transcriptional factor families (1611 mouse transcription factors) sourced from AnimalTFDB4 (https://guolab.wchscu.cn/AnimalTFDB4/), the most comprehensive animal transcription factor database with precise identification and thorough annotation^61^. AUROC values for each family within these gene sets was documented in Extended Data Table 5. The gene sets or families with AUROC scores >0.75 (regarded as a stringent threshold) are considered as being discriminating^54^.

To identify cell type-specific gene expression patterns, we applied the tspex Python package (version 0.6.2) to our snRNA-seq dataset to calculate gene expression specificity (z-score) for each cell type^215^. We focused on cell adhesion molecules (CAMs) and transcription factors (TFs), identifying 135 CAMs and 299 TFs with a specificity z-score > 0.75 (Extended Data Table 6). Since CN neurons are comprised of three major classes arising from distinct lineages — projection neurons, inhibitory neurons, and granule cells/unipolar brush cells (UBCs)^62,151,205^— we also calculated gene expression specificity within each of these major neuronal classes for TF and CAMs (Fig. 6). ^64,163,216^

### Synergistic core identification

Based on the notion that cell type identity can be determined by a combination of a few key TFs that exhibit synergistic transcriptional regulation, we used a novel computational platform that performs a dynamic search for an identity core composed of TFs that possess an optimal synergistic coexpression pattern using an information theoretic measure of synergy computed from the snRNA-seq dataset^77^. For each target subpopulation, a set of TFs that are preferentially expressed in the target cell type (“specific TFs”) is identified. Then most synergistic subset of specific TFs is sought by computing multivariate mutual information (MMI). Once a most synergistic core is found, it is extended by computing MMI with TFs expressed in both target and background cell types (non-specific TFs). Each TF core thus form by a set of specific TFs and a few non-specific TFs given that non-specific TFs also contribute to the synergistic expression specifying cell type identity^77^. We performed the synergistic core analysis on principal projection neurons or inhibitory neurons using default parameters^77^. The synergistic TF cores for each cell type were reported in Extended Data Table 8.

### FISH image analysis and quantification

Confocal images were captured from FISH-stained brain slices using LSM 710 or 880 confocal microscopes (Zeiss), and the image files (.lsm) containing image stacks were loaded into IDL for analysis of the distribution and co-localization of various mRNA transcripts. From each slice, the images were generated to capture either green fluorescence signals (FITC) or red signals (DIG). The maximum projected image was filtered with a Gaussian filter to remove background signals. To detect a cell with expression signals, cell regions were segmented using a customized IDL code (Extended Data Fig. 15). A cell region with an area smaller than 80 µm^2^ or larger than 800 µm^2^ was not counted as a cell. A cell region with a circularity of less than 0.4 was not counted as a cell either. The circularity of a cell was calculated using the function: 𝐶 = 4𝜋𝑎^2^⁄𝑝^2^. Here, *C* is the circularity, *a* is the area of a cell region and *p* is the perimeter of the cell region. Cell regions outside the CN area were not included for further analysis. The spatial position of a cell was set as the median of the x or y positions of the pixels within a detected cell region. The fluorescence intensity (F) of a detected cell (either green or red) was set as the mean intensity of all pixels within a detected cell region. After removing fluorescence intensity values within a cell region, the images of each channel were then smoothed by a 60 µm length smoothing window. The smooth images were counted as the background signals of each channel. The background signals (F_0_) of a detected cell were set as the mean values of the background intensities within a cell region. The intensity differences (ΔF) between F and F_0_ divided by F_0_ of a detected cell from both channels (green and red) were used to generate a 2-D scatter plot in Python, with each axis representing either channel. From this scatter plot, we used a method similar to the Flow Cytometry gated analysis to determine whether a cell was single-labeled or double-labeled (Extended Data Fig. 15). The number of single-labeled and double-labeled cells was counted across 5-8 consecutive sagittal sections of CN. The distribution of labeled cells (single or double) across CN were quantified through the A-to-P and D-to-V axis with Jointplot from Seaborn python package. See https://github.com/bcmjianglab/cn_project/tree/main/FISH_analysis for more details about FISH image analysis.

ISH images in Extended Data Fig. 8 were acquired from the Allen Mouse Brain Atlas (www.mouse.brain-map.org/)^217^. Single discriminatory markers were shown for each cell type, with several ISH images being repeated. ISH images were acquired with minor contrast adjustments as needed to maintain image consistency.

## Data Availability

Original electrophysiological, morphological, and RNA-sequencing data supporting the findings of this study have been deposited to the Mendeley Data (doi:10.17632/2n8m8k6b2g.1). The raw snRNA-seq and Patch-seq datasets have been deposited in the NCBI database with GEO accession number: GSE273145.

## Code Availability

The original code has been deposited at GitHub (https://github.com/bcmjianglab/cn_project).

## Acknowledgments

We thank the members of Jiang and Trussell labs for technical assistance, discussions, and comments on the manuscript. Research reported in this publication is supported by research grants R01 MH122169, R01 MH120404, R01 NS110767 (to X.J.), R01 DC004450, R35NS116798 (to L.O.T.), and R01 DC017797 (to M.J.M) from the National Institutes of Health. Research reported in this publication was also supported by the Main Street America Fund, a Shared Instrumentation grant from the NIH (1S10OD016167), by the Eunice Kennedy Shriver National Institute of Child Health & Human Development of the National Institutes of Health under Award Number P50HD103555 for use of the Microscopy Core facilities and the RNA In Situ Hybridization Core facility at Baylor College of Medicine, and by the NEI Core Grant for Vision Research (EY-002520-37) for use of the Bioengineering Core. We thank Drs. Lisa Goodrich and Arpiar Saunders for the comments on the manuscript and Drs. Dmitry Kobak, Philipp Berens, Yves Bernaerts, and Satoshi Okawa for helping with data analysis. The content is solely the responsibility of the authors and does not necessarily represent the official views of the National Institutes of Health.

## Author Contributions

J.J., T.N., X.J., Q.M., S.L., C.L. performed the experiments, J.J., M.H., X.J., T.N. analyzed the data. X.J., L.T. M.M. supervised all experiments and data analyses. X.J., L.T. prepared the manuscript with input from all co-authors.

## Extended Data Figures and Legends

**Extended Data Fig. 1:**
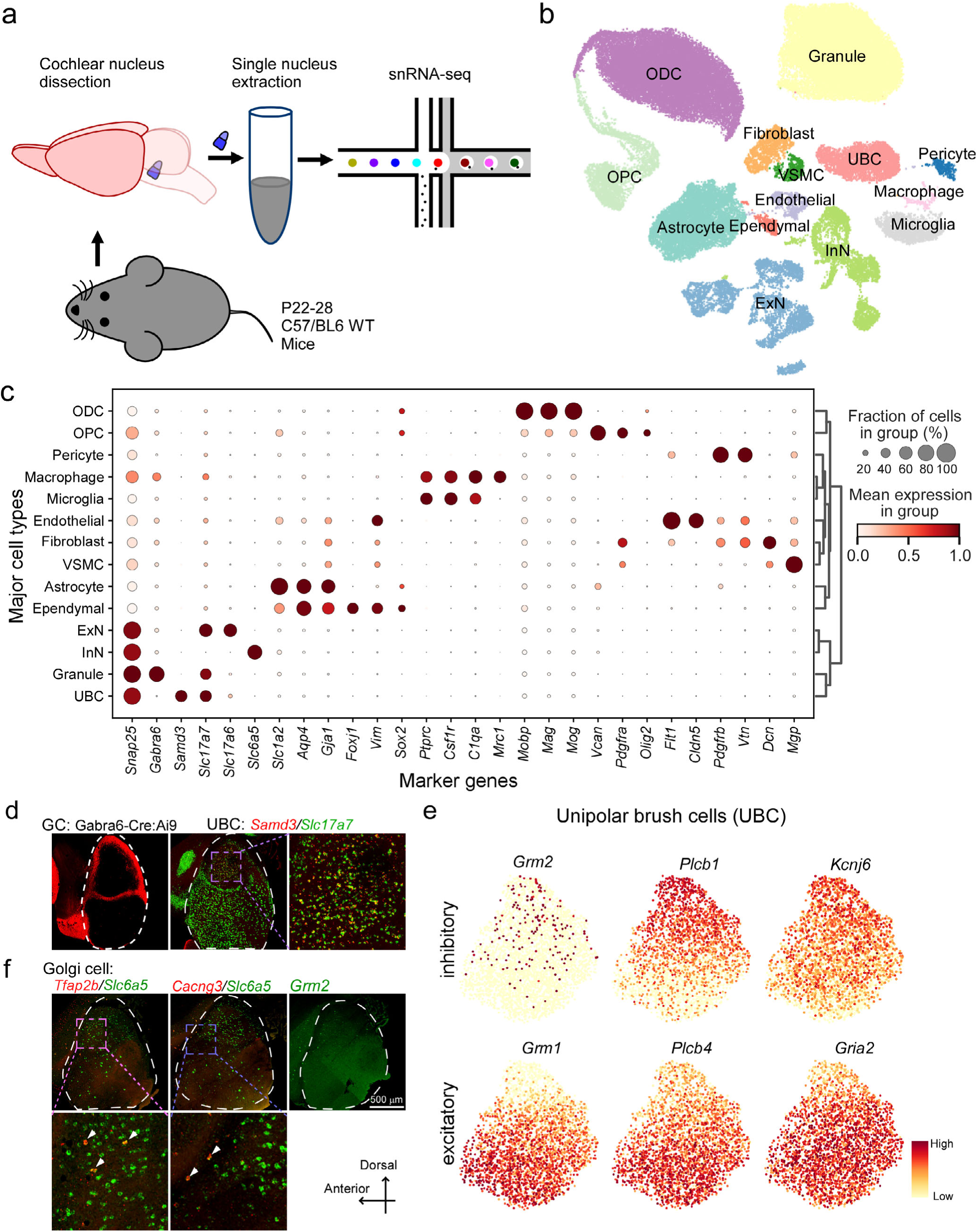
Single-nucleus transcriptomic profiling of mouse CN. Related to Fig. 1. **a**, Method flowchart. **b**, UMAP visualization of 61,211 nuclei from the CN (after profile quality control, removing nuclei from adjoining brain tissues, and cell type annotation; Methods), colored by cell type identity. ODC, oligodendrocyte; OPC, oligodendrocyte precursor cell; ExN: excitatory neurons; InN: inhibitory neurons; VSMC: vascular smooth muscle cells; Endothelial: endothelial cells; Fibroblast: fibroblast-like cells; Ependymal: ependymal cells; Granule: granule cells; UBC: unipolar brush cells. **c**, Dendrogram indicating hierarchical relationships between cell types (right) with a dot plot (left) of scaled expression of marker genes for major cell types as shown in (**b**). Cell type assignments, based on the expression of known marker genes, are indicated on the right (also see Extended Data Table 1). Circle size depicts the percentage of cells in the cluster in which the marker was detected (≥ 1 UMI), and the color depicts the average transcript count in expressing cells. **d**, Left: A representative confocal micrograph showing labeled cells in the CN of Gabra6-Cre:Ai9 mice. Note: labeled cells are restricted to granule cell layer. Middle: FISH co-staining for *Slc17a7* and *Samd3* in a representative CN sagittal section. Note: *Samd3*^+^ excitatory neurons (*Samd3*^+^/*Slc17a7*^+^, yellow signals) were restricted to the deep layer of DCN. Right: zoom-in of boxed region of the image in the middle. **e**, UMAP visualization of normalized expression of glutamate inhibitory signaling pathways (*Grm2, Plcb1,*and *Kcnj6*, genes enriched in OFF UBCs) and glutamate excitatory signaling pathways (*Grm1, Plcb4,* and *Gria2,* genes enriched in ON UBC). **f**, Top: FISH co-staining for *Tfap2b* and *Slc6a5* (left), or for *Cacng3* and *Slc6a5* (middle), and FISH single-staining for *Grm2* (right) in CN sagittal sections. Bottom: zoom-in of boxed region of the images on the top. Arrows indicate double-labeled cells.

**Extended Data Fig. 2:**
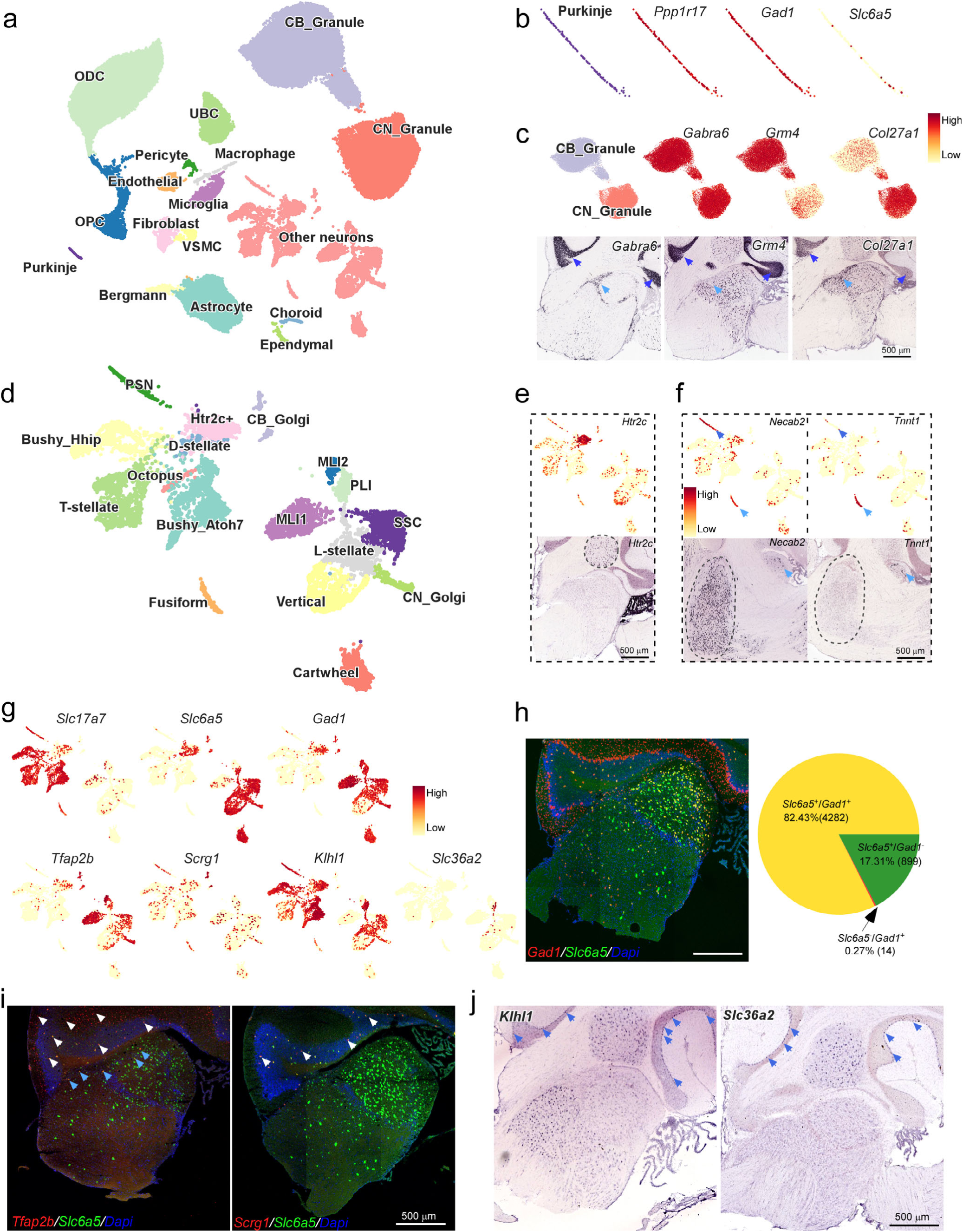
Identification of cell populations from adjoining brain tissues. Related to Fig. 1, and Extended Data Fig. 1. **a**, UMAP visualization of 93,644 nuclei (after quality control) reveals major cell types and clusters from the cochlear nuclei (CN) and neighboring cerebellum (CB). Cell type and cluster identity is color coded. Identification of cell types and clusters from CB, including cerebellar granule cells (CB_Granule), Purkinje cells, and Bergmann cells was based on their previously-identified molecular signatures^14^ and ISH data from Allen Brain Atlas. **b**, UMAP visualization of the expression of *Ppp1r17*, *Gad1*, and *Slc6a5* in the cluster as labeled as Purkinje in (**a**). **c**,Top: UMAP visualization of the expression of *Gabra6*, *Grm4*, and *Col27a1* in two clusters assigned to granule cells in (**a**) reveals a granule cell population from CB^14^ (CB_Granule) (i.e., *Grm4*^+^/*Col27a1*^-^). Bottom: Representative Allen Brain Atlas micrographs of ISH expression of Gabra6, Grm4, and Col27a1 in representative sagittal sections containing the CN and CB indicate that granule cells in CN (CN_Granule) could be discriminated from CB granule cells (CB_Granule) by their lack of *Grm4* expression. Note: Dark blue arrows point to granule cell layers in CB and light blue arrows point to granule cell layers in CN. Image credit for all ISH images: Allen Institute. **d**, Subclustering of the neuronal population (except for granule cells and UBC) reveals distinct neuronal clusters from the CN and adjoining regions including CB, the principal sensory nucleus of the trigeminal nerve (PSN), and the dentate nucleus. Cell type and cluster identity is color coded. Identification of neuronal clusters from CB, PSN and the dentate nucleus is based on identified molecular signatures of CB cell types^14^ and ISH data from Allen Brain Atlas. **e**, Top: UMAP visualization of the expression of *Htr2c* reveals a *Htr2c-*enriched cluster corresponding to cells from the dentate nucleus. Bottom: A representative micrograph (from Allen Brain Atlas) of ISH expression of *Htr2c* in a sagittal section containing CN and CB shows that *Htr2c* expression is restricted to the dentate nucleus (dash circle). **f**, Top: UMAP visualization of the expression of *Necab2* and *Tnnt1* reveals two *Necab2*-enriched clusters, one of which corresponds to cells from the PSN (*Necab2*^+^/*Tnnt1*^-^ cluster). Bottom: Micrograghs of ISH expression of *Necab2* (right) or *Tnnt1* (left) in a sagittal section containing a small region of DCN and PSN show that *Necab2*^+^/*Tnnt1*^-^ cells are restricted to the PSN (dash circles), while *Necab2^+^/Tnnt1^+^*cells (light blue arrows) are restricted to the DCN (i.e., fusiform cells; see Fig. 2). **g**, UMAP visualization of the expression of gene markers for excitatory neurons (*Slc17a7*), for inhibitory neurons (*Gad1* and *Slc6a5*), for CB interneuron cell types (*Tfap2b*, *Scrg1*, *Klhl1*, and *Slc36a2*) reveals four *Tfap2b*-enriched inhibitory clusters, and one inhibitory cluster co-enriched with *Klhl1* and *Slc36a2*. Two of *Tfap2b*-enriched clusters are pure GABAergic (*Gad1*^+^/*Slc6a5*^-^), which correspond to MLI1 and MLI2 from the CB^14^ (also see below). The other two *Tfap2b*-enriched are glycine/GABA combinatorial clusters (*Scrg1*^+^or *Scrg1*^-^), which correspond to Golgi cells in the CB (CB_Golgi^14^, also see below) and CN (CN_Golgi, also see below). An inhibitory cluster is co-enriched with *Klhl1* and *Slc36a2*, reminiscent of PLI in the CB^14^. **h**, Left: Double FISH staining of Gad1 and Slc6a5 in a sagittal section containing CN and CB. Right: Pie chart indicates the proportion of pure GABAergic neurons (*Gad1*^+^/*Slc6a5*^-^), pure glycinergic neurons (*Gad1*^-^/*Slc6a5*^+^), and glycine/GABA neurons (*Gad1*^+^/*Slc6a5*^+^). Note: *Gad1*^+^/*Slc6a5*^-^ neurons in the CN are absent or very sparse. **i**, Double FISH staining of Golgi cell marker gene (*Tfap2b* or *Scrg1*) and *Slc6a5* in representative sagittal sections containing CN and CB. Note: *Scrg1* expression is restricted to the CB. White arrows indicate the gene expression in CB and light blue arrows indicate the expression in CN. **j**, Representative micrographs (from Allen Brain Atlas) of ISH expression of *Klhl1* and *Slc36a2* (CB PLI marker gene) in the sagittal section containing the CN and CB. Note: *Slc36a2* is not expressed in CN. Arrows indicate the gene expression in CB Purkinje cell layer.

**Extended Data Fig. 3:**
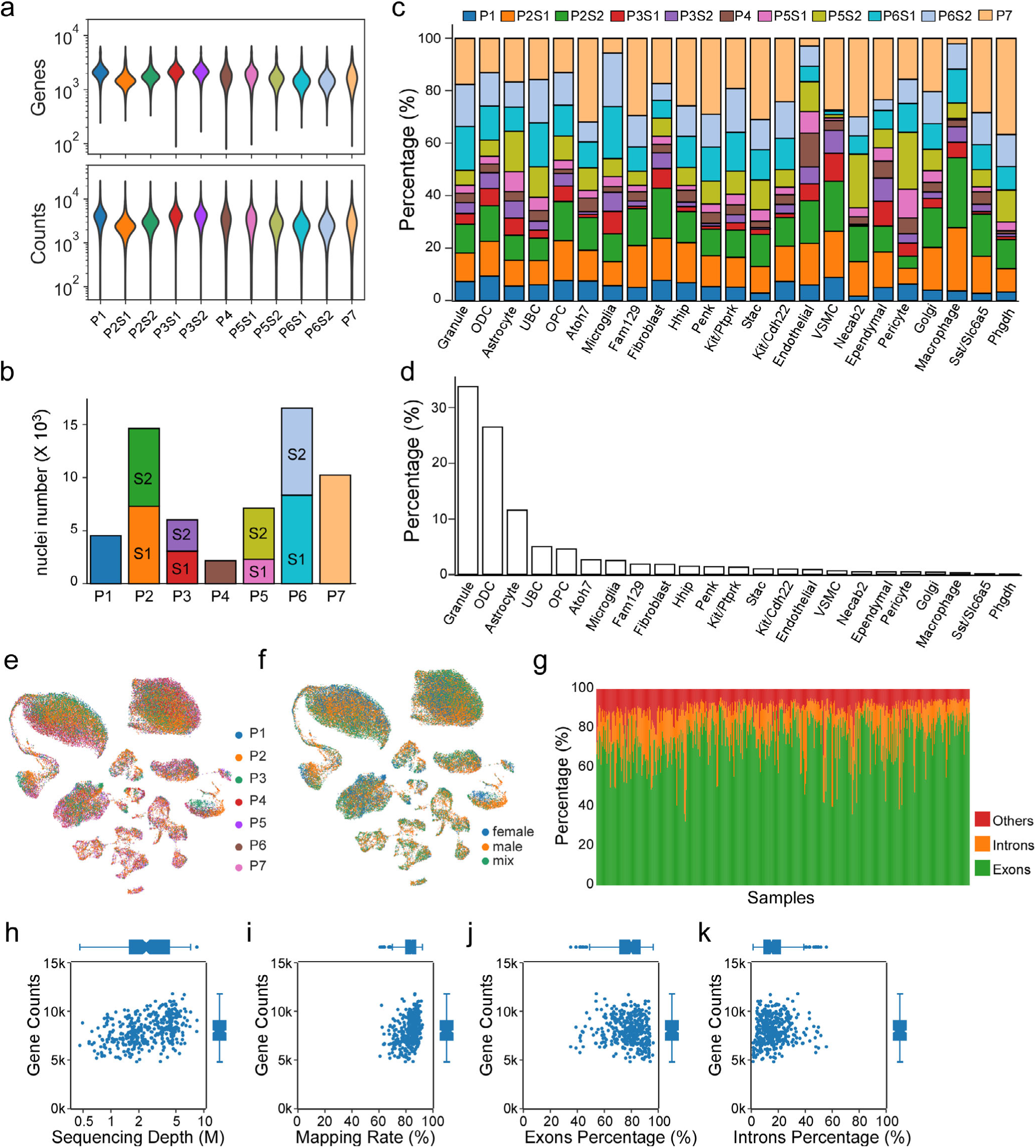
Quality control analysis for snRNA-seq and Patch-seq data. Related to Fig. 1-2, 5. **a**,Violin plots of the gene counts (top) and UMI counts (bottom) of each sample sequenced with Drop-seq. P1, P4, P7: the sample from Preparation 1, 44, and 7 respectively. P2S1 and P2S2: Sample 1 and 2 from Preparation 2; P3S1 and P3S2: Sample 1 and 2 from Preparation 3; P5S1 and P5S2: Sample 1 and 2 from Preparation 5; P6S1 and P6S2: Sample 1 and 2 from Preparation 6. **b**, Sample size (the number of nuclei) of each preparation and sample in snRNA-seq. Preparation 2, 3, 5, and 6 has two samples (S1, S2). **c**, The contribution of each sample (represented as a percentage) to the total population of each cell type (or cluster). See Fig. 1 and Extended Data Fig. 1 for annotation of cell types and clusters. **d**, The proportion of each cell type (or cluster) in the total sequenced population (i.e., combining all samples). **e**, UMAP visualization of all nuclei from CN, colored by preparations (P1-P7). **f**, UMAP visualization of all nuclei from CN, colored by gender. **g**, The proportion of sequencing reads mapped to exons (CDS, 5’UTR, 3’UTR), introns, and other intergenic regions in each Patch-seq cell. As expected, most reads map to exons regions rather than introns and intergenic regions. **h-k**, Correlations of gene counts with sequencing depth, STAR mapping rate, the percentage of reads mapped to exons and introns in Patch-seq cells.

**Extended Data Fig. 4:**
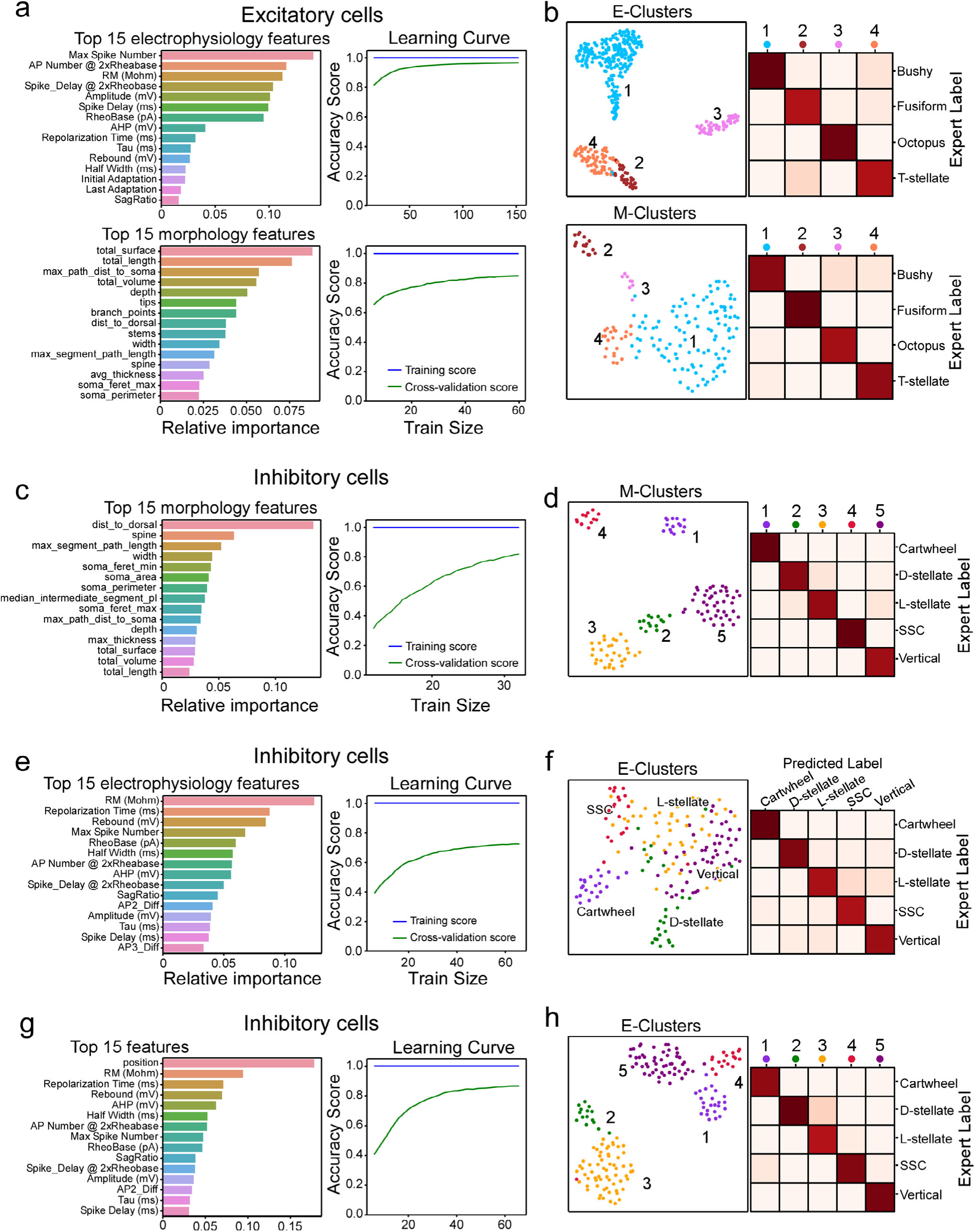
Distinguish cell types by electrophysiological or morphological features. Related to Fig. 2, 5. **a**, Left: The top 15 electrophysiological (top) or morphological (bottom) features contributing to the overall performance of the random forest classifier (for excitatory cell types) are shown, with weights indicateing their relative importance. Right: The performance of the classifier trained with electrophysiological (top) or morphological (bottom) features. Note: its performance improves as the training dataset size increases, indicating a good cell type classifier. **b**, Top left: UMAP visualization of 404 CN excitatory neurons clustered by select electrophysiological features, colored by k-means clusters. Top right: Confusion matrix shows the performance of k-means clustering (with electrophysiological features) in distinguishing CN excitatory cell types as labeled manually (an average of 93.8% accuracy). Bottom left: UMAP visualization of 157 CN excitatory neurons clustered by select morphological features, colored by k-means clusters. Bottom right: Confusion matrix shows the performance of k-means clustering (with morphological features) in distinguishing CN excitatory cell types (an average of 93.0% accuracy). **c**, Left: The top 15 morphological features contributing to the classifier’s performance (in identifying inhibitory cell types) are shown, with weights indicating their relative importance. Right: The performance of the classifier trained with morphological features. **d**, Left: UMAP visualization of 116 CN inhibitory neurons clustered by select morphological features, colored by k-means clusters. Right: Confusion matrix shows the performance of k-means clustering (with morphological features) in distinguishing CN inhibitory cell types (an average of 92.2% accuracy). **e**, Left: The top 15 electrophysiological features contributing to the classifier’s performance (in identifying inhibitory cell types) are shown, with weights indicating their relative importance. Right: The performance of the classifier trained with electrophysiological features only. **f**, Left: UMAP visualization of CN inhibitory neurons (n=172) clustered by select electrophysiological features, colored by expert cell type as in Fig. 5c. Right: Confusion matrix shows the performance (an average of 86.8% accuracy) of the cell-type classifier trained with electrophysiological features only. **g**, Left: The top 15 features contributing to the classifier’s performance (in identifying inhibitory cell types) are shown, with weights indicating their relative importance. These features include both electrophysiological features and anatomical location (VCN=0, DCN=1). Right: The performance of the classifier trained with electrophysiological features and anatomic location. **h**, Left: UMAP visualization of 172 CN inhibitory neurons clustered by electrophysiological features and anatomic location, colored by k-means clusters. Top right: Confusion matrix shows the performance of k-means clustering (with electrophysiological features and anatomic location) in distinguishing CN inhibitory cell types as labeled manually (an average of 90.7% accuracy).

**Extended Data Fig. 5:**
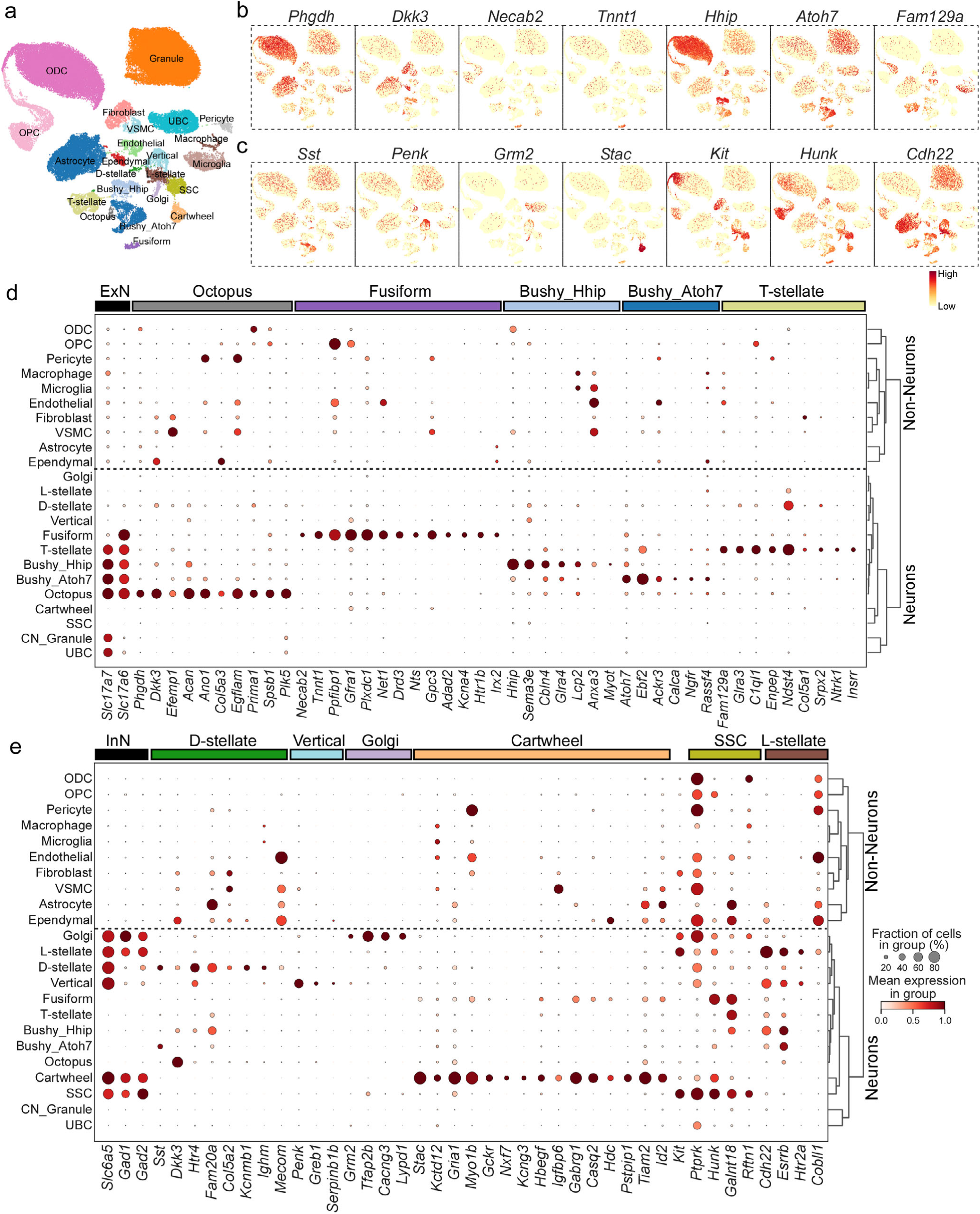
The expression of marker genes for major CN neuronal types. Related to Fig. 3-5. **a**, UMAP visualization of all CN nuclei (after profile quality control and annotation; Methods), colored by cell type and cluster identity. **b**, UMAP visualization of the normalized expression of the key discriminatory genes for each CN excitatory cell type across all cell types. **c**, UMAP visualization of the expression of the key discriminatory genes for each CN inhibitory cell type across all cell types. **d**, Dendrogram indicating hierarchical relationships between cell types (or clusters, right) with a dot plot of scaled expression of selected marker genes for major excitatory cell types (left). **e**, Dendrogram indicating hierarchical relationships between cell types (or clusters) with a dot plot of scaled expression of selected marker genes for inhibitory cell types.

**Extended Data Fig. 6:**
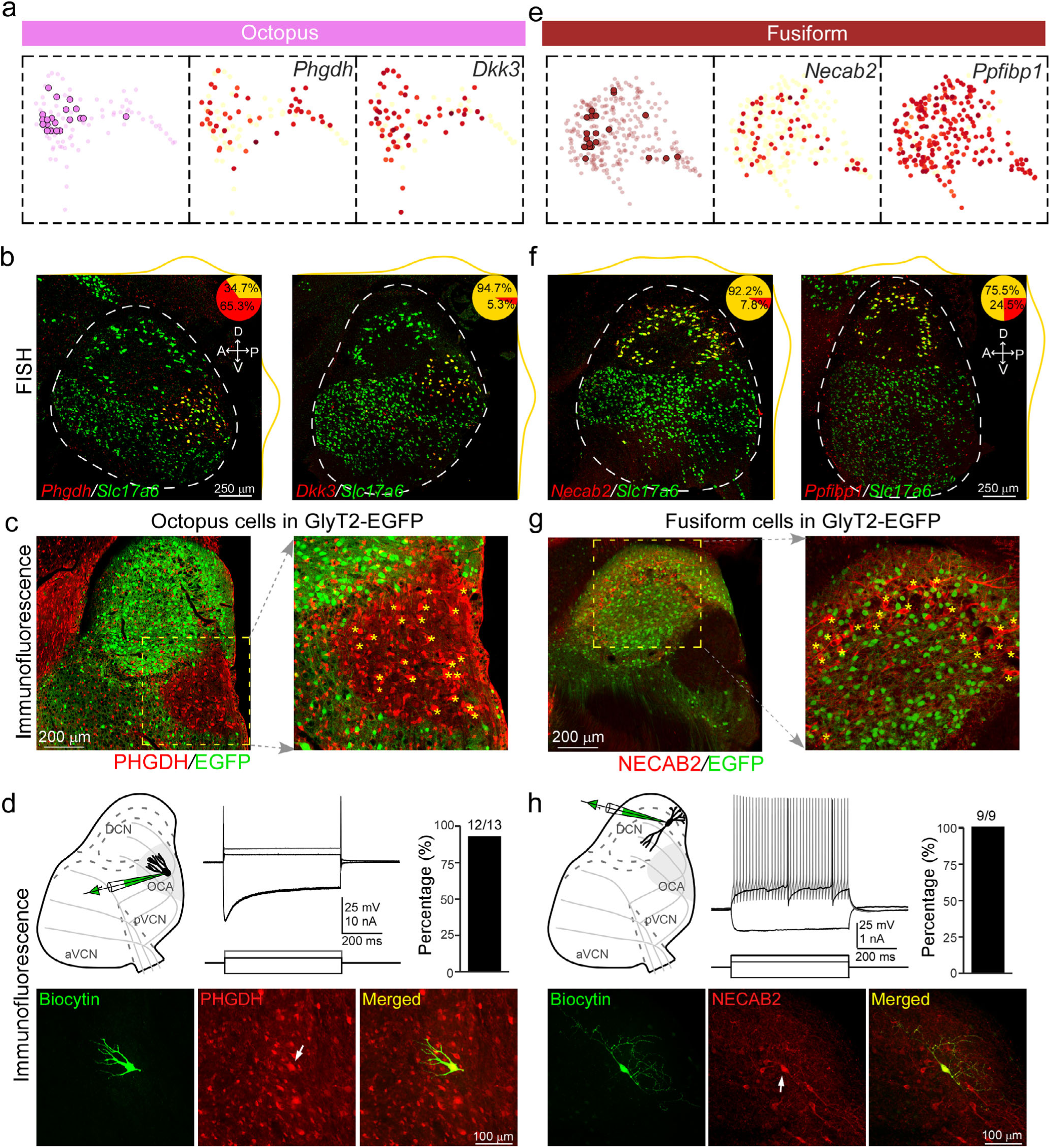
Novel marker genes for CN excitatory neurons. **a**, UMAP visualization of the *Phgdh*^+^ cluster (n=88) alongside all Patch-seq-derived octopus cells (highlighted as dark pink circle, left). Middle and Right: UMAP visualization of normalized expression of two discriminatory markers for octopus cell cluster. **b**, FISH co-staining for *Slc17a6* and *Phgdh* (left), or *Slc17a6* and *Dkk3* (right) in sagittal CN sections. D, dorsal; V, ventral; A, anterior; P, posterior. Inset pie charts depict proportion of double-labeled cells in single-labeled cells (*Phgdh*^+^ or *Dkk3*^+^). Yellow lines along images show density of double-labeled neurons along two axes of CN. **c**, Immunostaining for PHGDH (red) in a sagittal section from GlyT2-EGFP mice (green EGFP indicates glycinergic neurons). Boxed region is magnified as on the right. Star signs (*) point to each soma with PHGDH signals. Labeled neurons are restricted to the OCA, which exhibits as a dark area under the fluorescent microscope due to a lack of glycinergic cells^10^. **d**, Top-left: diagram showing patch recording and labeling with biocytin for octopus cells. Top-middle: example responses of an octopus cell to current steps. Top-right: proportion of octopus cells immunopositive for PHGDH. Bottom: example octopus cell filled with biocytin (green) positive for PHGDH (red). **e**, UMAP visualization of *Necab2*^+^ cluster (n=274) and all Patch-seq fusiform cells positioned on UMAP space (left). Middle and Right: UMAP visualization of normalized expression of two discriminatory markers for fusiform cells. **f**, FISH co-staining for *Slc17a6* and *Necab2* (left), or *Slc17a6* and *Ppfibp1* (right) in CN sagittal sections. Inset pie charts depict proportion of double-labeled cells in single-labeled cells (*Necab2*^+^ or *Ppfibp1*^+^). **g**, Immunostaining for NECAB2 (red) in a sagittal section from GlyT2-EGFP mice. Boxed region is magnified as on the right. Star signs (*) point to each soma with NECAB2 signals. **h**, Top-left: diagram showing patch recording and labeling for fusiform cells. Top-middle: example responses of a fusiform cell to current steps. Top-right: proportion of fusiform cells immunopositive for NECAB2. Bottom: example fusiform neuron filled with biocytin (green) shows positive for NECAB2 (red).

**Extended Data Fig. 7:**
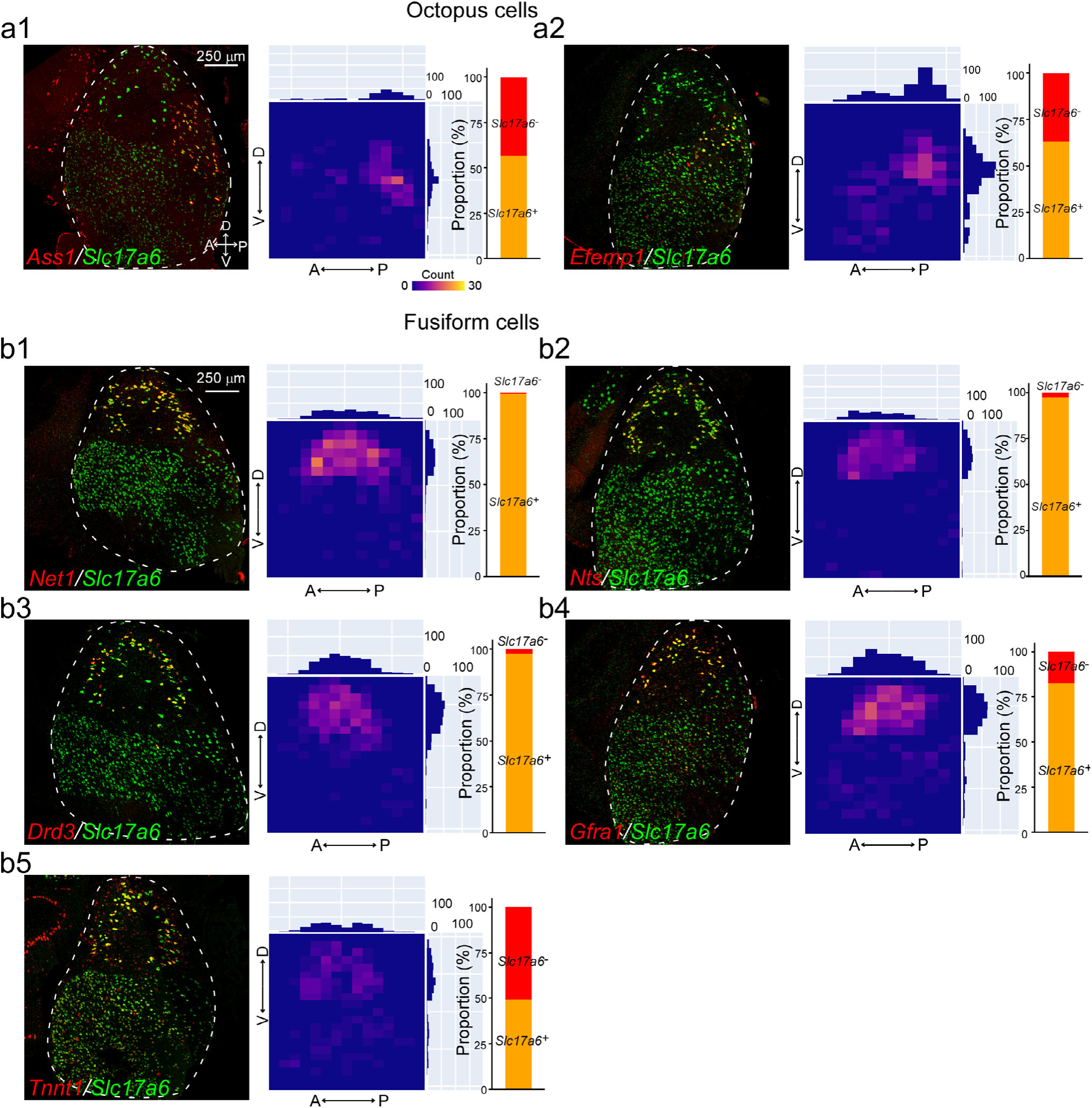
FISH staining and immunostaining of gene markers for fusiform cells and octopus cells. **a1**, FISH co-staining for *Slc17a6* and *Ass1* in a representative CN sagittal section (left). Heatmap depicts density distribution of double-labeled cells across the CN, and bar graph depicts proportion of double-labeled cells within the total *Ass1*^+^ cells. **a2**, FISH co-staining for *Slc17a6* and *Efemp1* in a sagittal section (left). Heatmap depicts density distribution of double-labeled cells and bar graph depicts proportion of double-labeled cells within the total *Efemp1*^+^ cells. **b1**, FISH co-staining for *Slc17a6* and *Net1* in a sagittal section (left). Heatmap depicts density distribution of double-labeled cells and bar graph shows proportion of double-labeled cells within the total *Net1*^+^ cells. **b2**, FISH co-staining for *Slc17a6* and *Nts* in a sagittal section (left). Heatmap depicts density distribution of double-labeled cells and bar graph depicts proportion of double-labeled cells within the total *Nts*^+^ cells. **b3**, FISH co-staining for *Slc17a6* and *Drd3* in a sagittal section (left). Heatmap depicts density distribution of double-labeled cells and bar graph depicts proportion of double-labeled cells within the total *Drd3*^+^ cells. **b4**, FISH co-staining for *Slc17a6* and *Gfra1* in a sagittal section (left). Heatmap depicts density distribution of double-labeled cells and bar graph depict proportion of double-labeled cells within the total *Gfra1*^+^ cells. **b5**, FISH co-staining for *Slc17a6* and *Tnnt1* in a sagittal section (left). Heatmap depicts density distribution of double-labeled cells and bar graph depicts proportion of double-labeled cells within the total *Tnnt1*^+^ cells.

**Extended Data Fig. 8:**
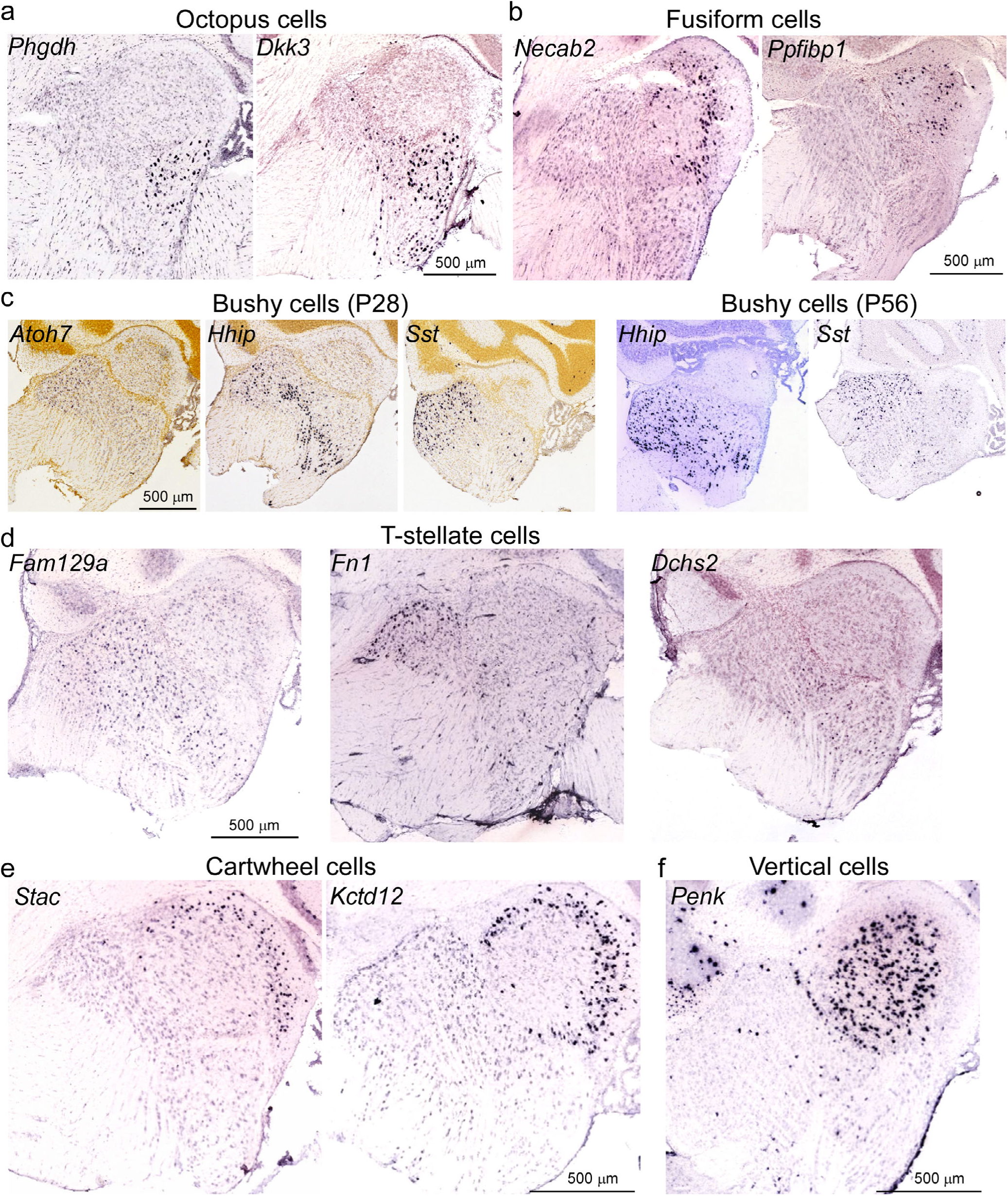
Allen Brain Atlas ISH expression of single discriminatory markers for major CN cell types. Related to Fig. 3-5. **a-b**, Representative micrographs of ISH expression of single discriminatory markers (*Dkk3* and *Phgdh*) for octopus cells(**a**) or single discriminatory markers (*Necab2* and *Ppfibp1*) for fusiform cells(**b**) in the CN sagittal sections. **c**, Representative micrographs of ISH expression of single discriminatory markers (*Atoh7*, *Hhip,* and *Sst*) for bushy cell subtypes in sagittal sections containing CN (corresponding to middle CN) at P28 (left) and P56 (right). The expression data of *Atoh7* at P56 are not available. **d**, Representative micrographs of ISH expression of *Fam129a*, a general discriminatory marker for T-stellate cells (right), and of the subtype-specific markers (*Fn1* and *Dchs2*) in the CN sagittal sections. **e-f**, Representative micrographs of ISH expression of single discriminatory marker for cartwheel cells (*Stac* and *Kctd12*, **e**) or for vertical cells (*Penk*, **f**) in CN sagittal sections. Image credit for all ISH images: Allen Institute.

**Extended Data Fig. 9:**
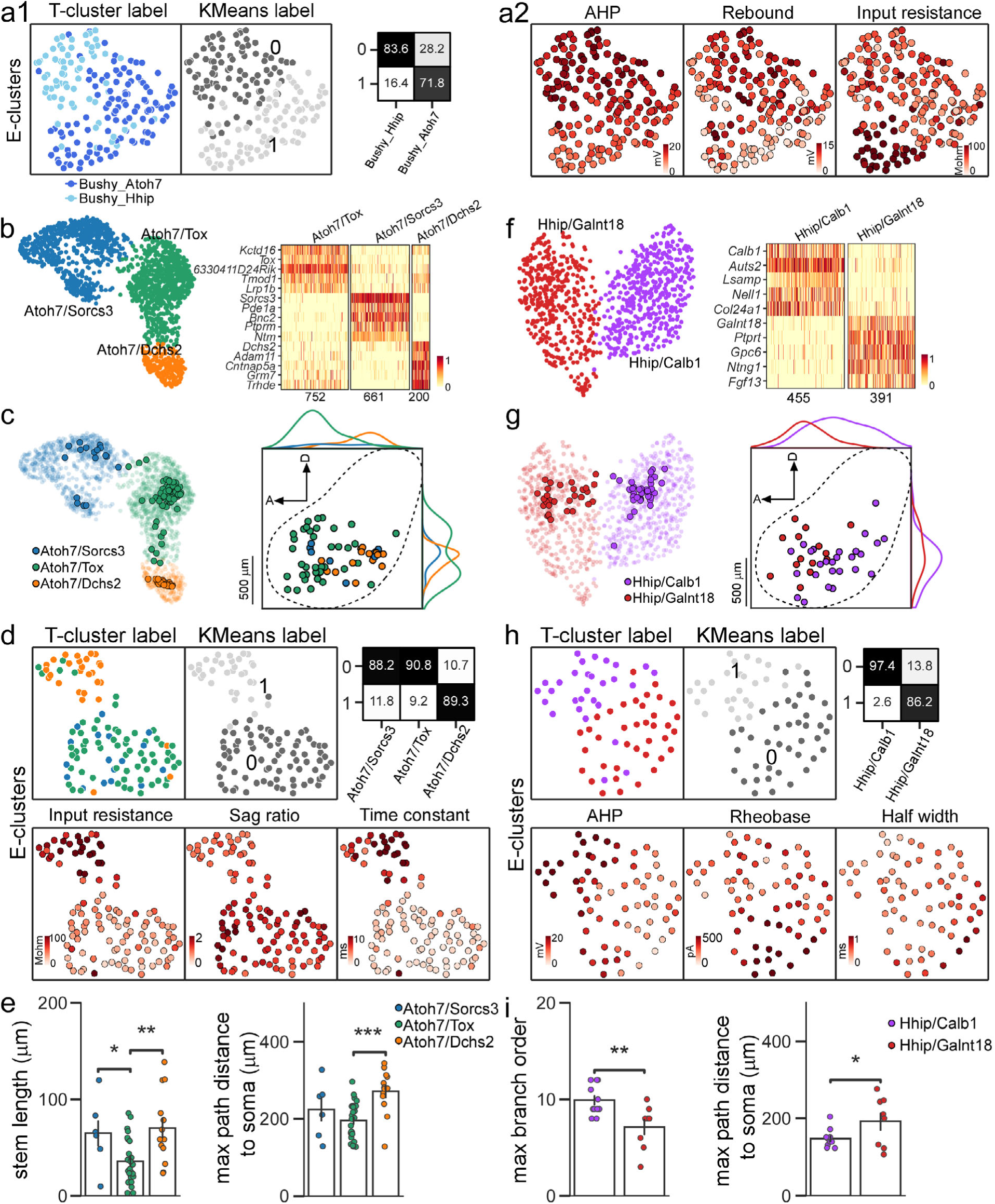
Molecular and phenotypic variability of bushy cells. Related to Fig. 3. **a1**, Left: UMAP visualization of bushy cells clustered by over-dispersed electrophysiological features, colored by identified subtypes (Bushy-Atoh7 and Bushy-Hhip). Middle: UMAP visualization of bushy cells clustered by electrophysiological features, colored by K-means clusters (Cluster 0 and 1). Right: Confusion matrix depicts the performance of k-means clustering in distinguishing two molecular subtypes, with an average accuracy of ∼80%. **a2,** UMAP visualization of busy cells by over-dispersed physiological features, colored by three key electrophysiological features: AHP, rebound, and input resistance. **b**, Left: UMAP visualization of *Atoh7^+^* bushy cells, colored by subcluster (Atoh7/Sorcs3, Atoh7/Dchs2 and Atoh7/Tox). Right: Heat map displaying normalized expression of the top 15 DEGs for three subclusters. Scale bar: Log2 (Expression level). The cell number per subcluster is indicated below. **c**, Left: All Patch-seq *Atoh7^+^* bushy cells mapped to each subcluster. Right: 2D spatial projection of each Patch-seq *Atoh7^+^* bushy cell onto a sagittal view of CN, colored by the subcluster. **d**, Top Left: UMAP visualization of *Atoh7^+^* bushy cells by over-dispersed electrophysiological features, colored by identified subclusters. Top Middle: UMAP visualization of *Atoh7^+^*bushy cells clustered by electrophysiological features, colored by K-means clusters (Cluster 0 and 1). Top Right: Confusion matrix depicts the performance of K-means clustering in distinguishing three subclusters. Bottom: UMAP visualization of *Atoh7^+^* bushy cells clustered by over-dispersed physiological features, colored by three electrophysiological features: input resistance, sag ratio, and time constant. **e**, Two morphological features in three *Atoh7^+^* subclusters. For the other features see Extended Data Table 3. **f**, Left: UMAP visualization of *Hhip^+^* bushy cells, colored by subcluster (Hhip/Galnt18; Hhip/Calb1). Right: Heat map displaying normalized expression of the top 10 DEGs for two subclusters. **g**, Left: All Patch-seq *Hhip^+^* bushy cells mapped to each subcluster. Right: 2D spatial projection of each Patch-seq *Hhip^+^* bushy cell onto a sagittal view of CN, colored by the subcluster. **h**, Top Left: UMAP visualization of *Hhip^+^* bushy cells clustered by over-dispersed electrophysiological features, colored by identified subclusters. Top Middle: UMAP visualization of *Hhip^+^* bushy cells by electrophysiological features, colored by K-means clusters (Cluster 0 and 1). Top Right: Confusion matrix depicts the performance of k-means clustering in distinguishing two subclusters. Bottom: UMAP visualization of *Hhip^+^* bushy cells, colored by three electrophysiological features: AHP, rheobase, and half-width. **i**, Two morphological features in two *Hhip^+^* subclusters. For the other features see Extended Data Table 3.

**Extended Data Fig. 10:**
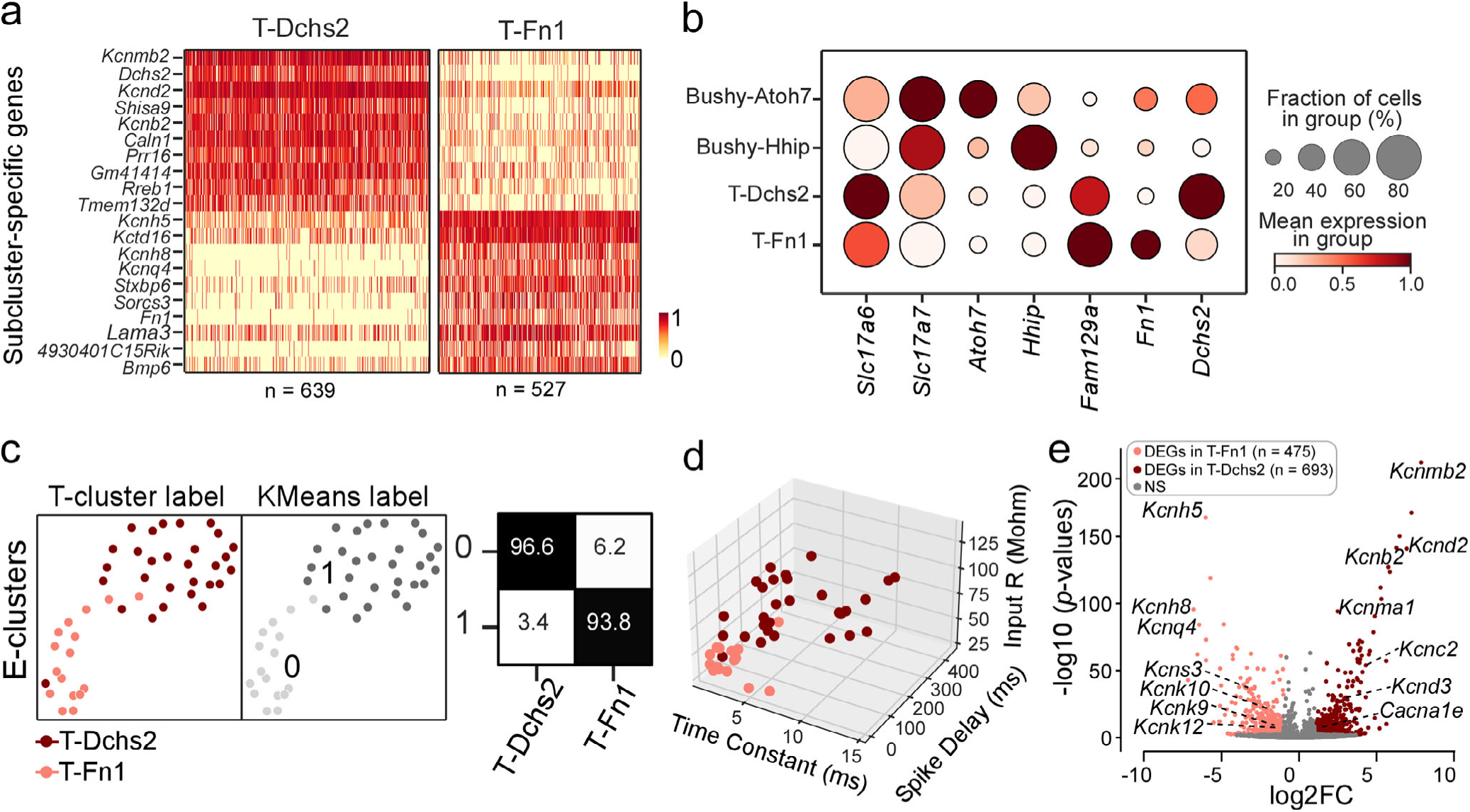
Molecular and phenotypic variation of T-stellate cells. Related to Fig. 4. **a**, Heat map displays normalized expression of the top 20 DEGs for two T-stellate cell subclusters. **b**, Dot plot illustrates the expression patterns of each marker gene for bushy cell and T-stellate cell subtype based on Patch-seq. **c**, Left: UMAP visualization of T-stellate cells clustered by over-dispersed electrophysiological features, colored by subtypes. Middle: UMAP visualization of T-stellate cells by electrophysiological features, colored by K-means clusters (Cluster 0 and 1). Right: Confusion matrix shows the performance of K-means clustering in distinguishing two subtypes. Volcano plot displays the log2 fold change (log2FC) and −log10(*p*-value) of detected genes comparing T-Fn1 to T-Dchs2. **d**, 3D scatter plot of three electrophysiological features of T-stellate cell. **e**, Volcano plot highlights 13 genes encoding potassium channel subunits and one gene encoding calcium channel subunit among DEGs. NS: not significant

**Extended Data Fig. 11:**
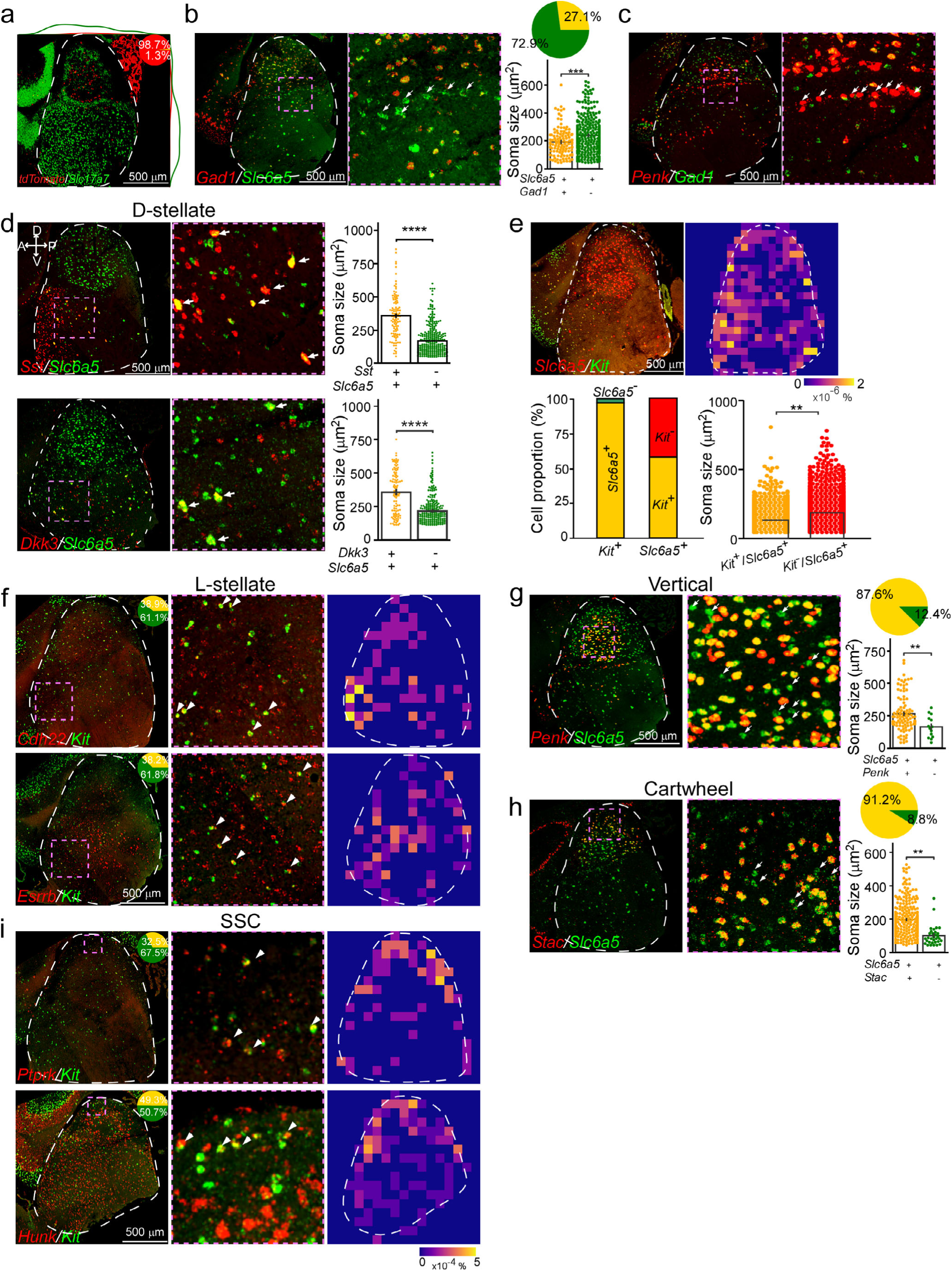
FISH staining for marker genes of inhibitory cell types. Related to Fig. 5. **a**, FISH co-staining for *Slc17a7* and *tdTomato* in Penk-Cre:Ai9 mice. Inset pie chart depicts the proportion of double-labeled cells (expressing both markers) within the total *tdTomato*^+^ cells. Colored lines illustrate density distribution of double-labeled neurons or single-labeled cells. **b**, Left: FISH co-staining for *Slc6a5* and *Gad1*. Middle: zoom-in of boxed area shown on the left. Arrows highlighting those *Gad1*^-^/*Slc6a5*^+^ cells in small-cell cap. Right: soma size of *Gad1*^-^/*Slc6a5*^+^ and *Gad1*^+^/*Slc6a5*^+^ cells. Pie chart depicts proportion of *Gad1*^-^ and *Gad1*^+^ glycinergic neurons in CN. **c**, Left: FISH co-staining for *Penk* and *Gad1*. Right: zoom-in of boxed area shown on the left. Arrows highlighting those cells in small-cell cap that are *Penk*^+^ but *Gad1*^-^. **d**, Left: FISH co-staining for *Sst* (top) or *Dkk3* (bottom) with *Slc6a5* in a representative CN sagittal section. Middle: Zoom-in of boxed region shown on the left. Right: soma size comparisons between *Sst*^+^ and *Sst*^-^ glycinergic neurons (top) or *Dkk*^+^ and *Dkk*^-^ glycinergic neurons (bottom). \*\*\*\**p* < 0.0001 with t-test. **e**, Top left: FISH co-staining for *Slc6a5* and *Kit*. Top right: Distribution of double-labeled cells. Bottom left: Bar graphs depict proportion of double-labeled cells (*Slc6a5*^+^/*Kit*^+^) and single-labeled cells (*Slc6a5*^-^/*Kit*^+^) in all *Kit*^+^ cells and proportion of *Kit*^+^ glycinergic neurons (*Kit^+^/Slc6a5^+^*) and *Kit*^-^ glycinergic neurons (*Kit^-^/Slc6a5^+^*). Note: only 2.8% of *Kit*^+^ cells are *Slc6a5*^-^. Bottom right: soma size of *Kit*^+^ and *Kit*^-^ glycinergic neurons. \*\**p* < 0.01 with t-test. **f**, Left: FISH co-staining for *Cdh22* or *Esrrb* with *Kit*. Inset pie chart depicts proportion of double-labeled cells in all *Kit*^+^ cells. Middle: zoom-in of boxed region shown on the left. Right: distribution of *Cdh22*^+^/*Kit*^+^ (top) or *Esrrb*^+^/*Kit*^+^ (bottom) cells. **g**, Left: FISH co-staining for *Slc6a5* and *Penk*. Middle: zoom-in of boxed area shown on the left. Right: soma size of *Penk*^-^/*Slc6a5*^+^ cells and *Penk*^+^/*Slc6a5*^+^ cells. Pie chart depicts proportion of *Penk*^-^ and *Penk*^+^ glycinergic neurons in the deep layer of DCN. **h**, Left: FISH co-staining for *Slc6a5* and *Stac*. Middle: zoom-in of boxed area shown on the left. Arrows point to those *Stac*^-^/*Slc6a5*^+^ cells. Right: soma size of *Stac*^-^/*Slc6a5*^+^ cells and *Stac*^+^/*Slc6a5*^+^ cells. Pie chart depicts proportion of *Stac*^-^ and *Stac*^+^ glycinergic neurons in DCN. **i**, Left: FISH co-staining for *Ptprk* or *Hunk* with *Kit*. Inset pie chart shows proportion of double-labeled cells in all *Kit*^+^ cells. Middle: zoom-in of boxed region shown on the left. Right: distribution of *Ptprk*^+^/*Kit*^+^ (top) or *Hunk*^+^/*Kit*^+^ (bottom) cells.

**Extended Data Fig. 12:**
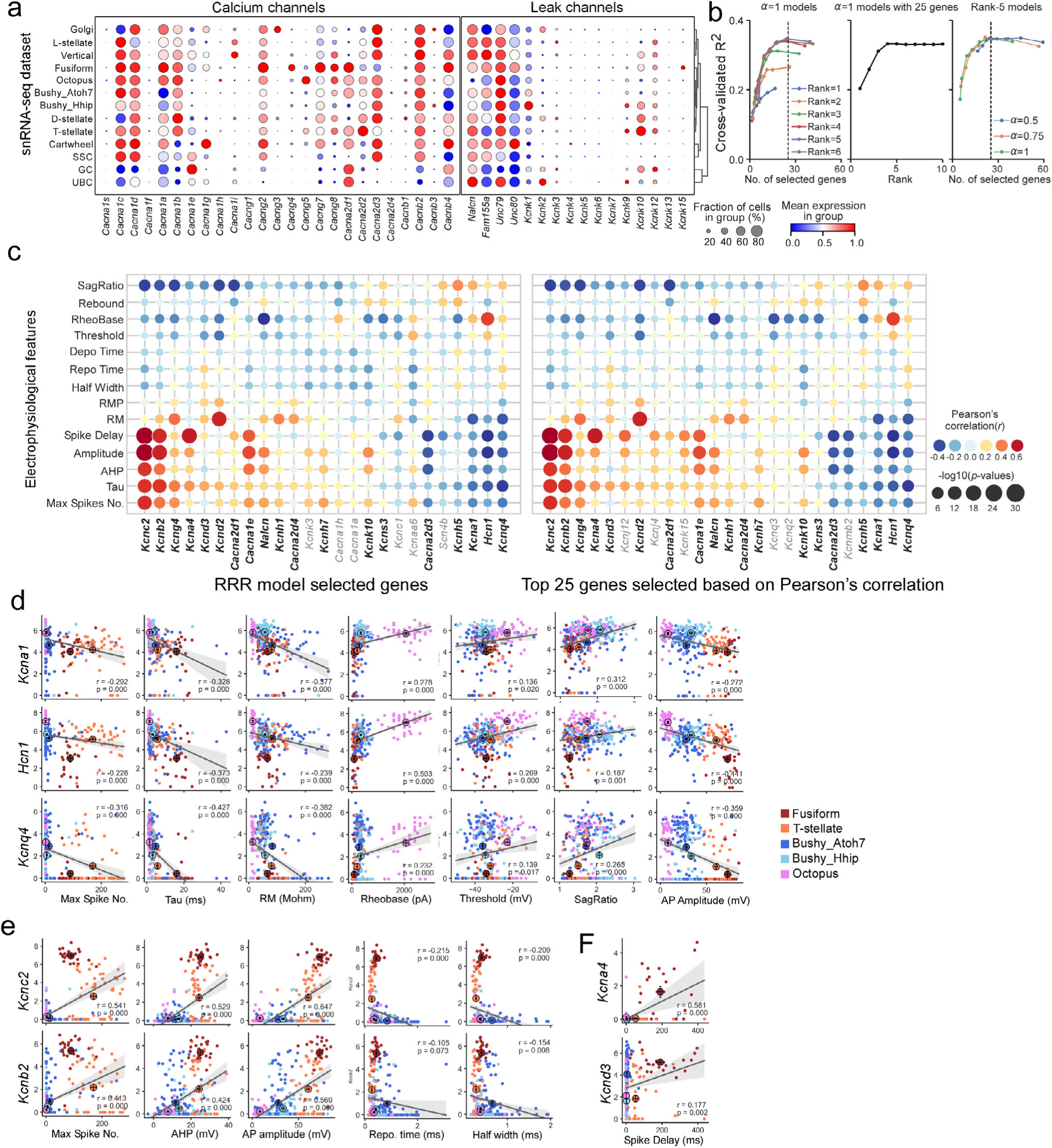
Single-cell transcriptomes provide insights into functional specializations of CN cell types. Related to Fig. 7. **a**, Dot plot displaying scaled expression of calcium channel-encoded genes and the genes encoding leak channels and associated proteins (from snRNA-seq dataset). **b**, Cross-validation performance (cross-validated R-squared) of the sparse RRR model across varying hyperparameters (alpha, rank, gene number). Vertical lines indicate performance at 25 selected genes. Near-optimal predictive accuracy is achieved with Rank = 5, alpha = 1, and 25 genes. **c**, Left: Dot plot depicting correlation coefficients between the expression of 25 ion channel genes selected by sparse RRR model and electrophysiological features (Pearson correlation analysis). Dot color indicates correlation coefficient (r), and dot size indicates −log10(p-values). Right: Dot plot depicting correlation coefficients between the expression of 119 ion channel genes and electrophysiological features. Only the top 25 genes with the highest correlation coefficients are shown, with 19 genes overlapping with sparse RRR model-selected genes (highlighted in black bold font). **d**, Scatter plots and linear correlations between gene expression and electrophysiological features across CN cell types. Significant negative correlations are observed between *Hcn1*, *Kcna1*, or *Kcnq4* and spike number (AP), Tau, input resistance (Rm), spike amplitude, respectively, and significant positive correlations are observed between these genes with rheobase, threshold, or sag. Each small circle represents a neuron (colored by cell type), and each large circle represents a cell type (mean ± SEM). Solid lines represent linear regression lines with 95% confidence intervals (gray lines). **e**, Scatter plots and linear correlations reveal significant positive correlations between *Kcnb2* and *Kcnc2* with spike number, AHP, or spike amplitude, and negative correlations with repolarization time and half-width. Each small circle represents a neuron (colored by cell type). Solid lines represent linear regression lines with 95% confidence intervals (gray lines). **f**. Scatter plots and linear correlations show significant positive correlations between *Kcna4* or *Kcnd3* and spike delay. Solid lines represent linear regression lines.

**Extended Data Fig. 13:**
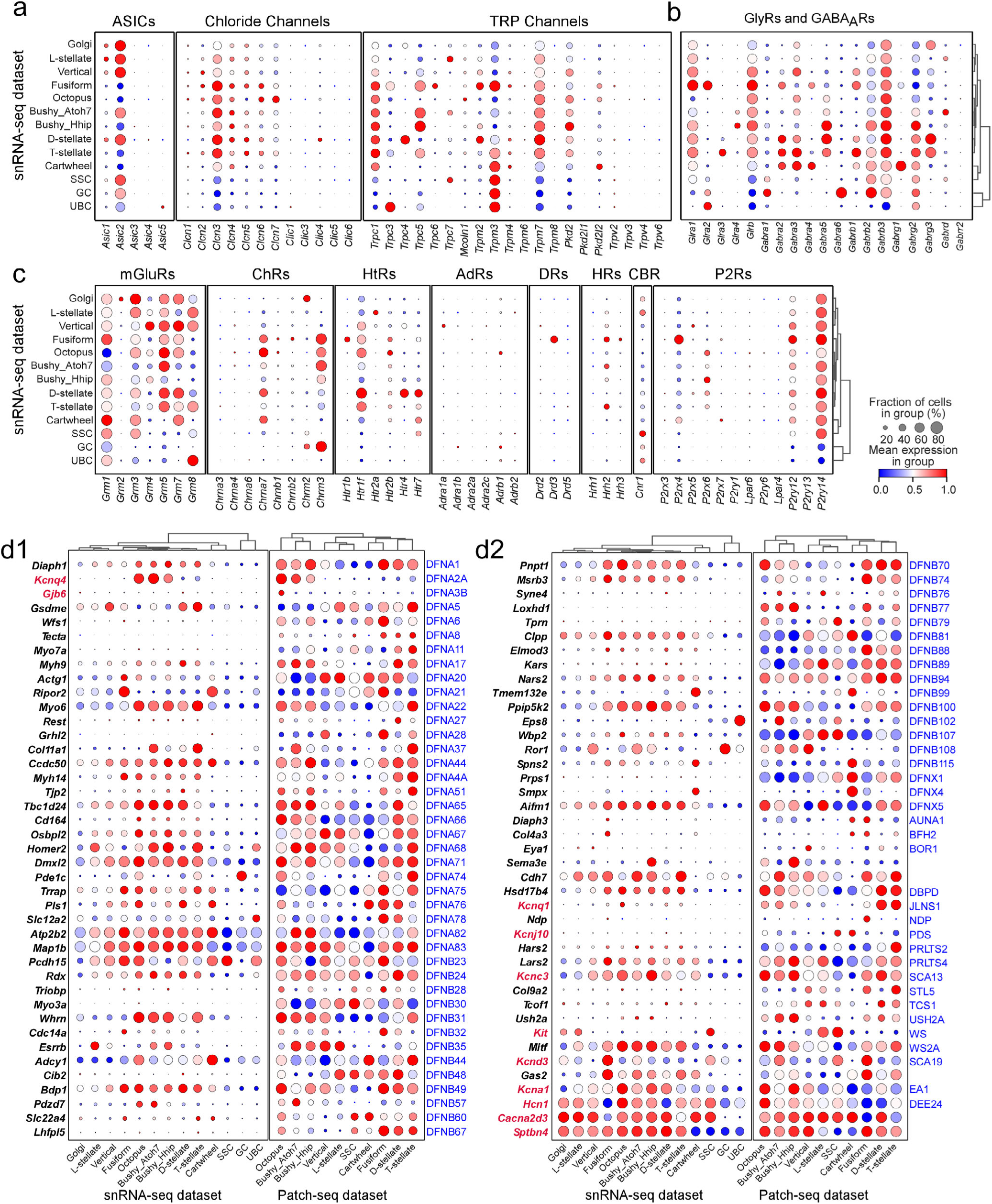
The expression of ion channels, GABA_A_ receptor, glycine receptors, metabotropic receptors, and deafness-related genes in CN. Related to Fig. 7. **a**, Dot plot displaying scaled expression of the genes encoding acid-sensing ion channel (ASICs), chloride channels, and transient receptor potential (TRP) channels across CN cell types (from snRNA-seq dataset). **b**, Dot plot displaying scaled expression of the genes encoding glycine and GABA_A_ receptors (from snRNA-seq dataset). **c**, Dot plot displaying scaled expression of the genes encoding metabotropic glutamate receptors (mGluR), cholinergic receptors (ChRs), serotonergic receptors (HtRs), adrenergic receptors (AdRs), dopaminergic receptors (DRs), histaminergic receptors (HRs), cannabinoid receptors (CBR), and purinergic receptors (P2Rs) across CN cell types (from snRNA-seq dataset). The Patch-seq data (not shown) generally agree with snRNA-seq data but detected a higher expression of several receptors in additional cell types, consistent with the more sequencing depth in Patch-seq compared to the 10X drop-seq protocol. **d1-2**, Dot plot displaying scaled expression of 82 deafness genes (out of 167 known genes) and the genes whose mutations cause hearing loss in human syndromes and mice (Left: snRNA-seq dataset; Right: Patch-seq dataset). Those deafness genes with consistently low expression (<20% cell fraction in any cell type across both datasets) were excluded (n=85). AUNA1: auditory neuropathy, autosomal dominant 1; BFH2: benign familial hematuria; BOR1: Branchiooterenal syndrome 1; DBPD: D-bifunctional protein deficiency; JLNS1: Jervell and Lange-Nielsen syndrome; NDP: Norrie disease; PDS: Pendred’s syndrome; PRLTS2: Perrault syndrome, type 2; PRLTS4: Perrault syndrome, type 4; SCA13: Spinocerebellar ataxia type 13; STL5: Stickler syndrome, type 5; TCS1: Treacher Collins syndrome-1; USH2A: Usher syndrome type II; WS: Williams syndrome; WS2A: Waardenburg syndrome type 2; SCA19: Spinocerebellar Ataxia Type 19. EA1: Episodic ataxia type 1, DEE24: Developmental and epileptic encephalopathy-24.

**Extended Data Fig. 14:**
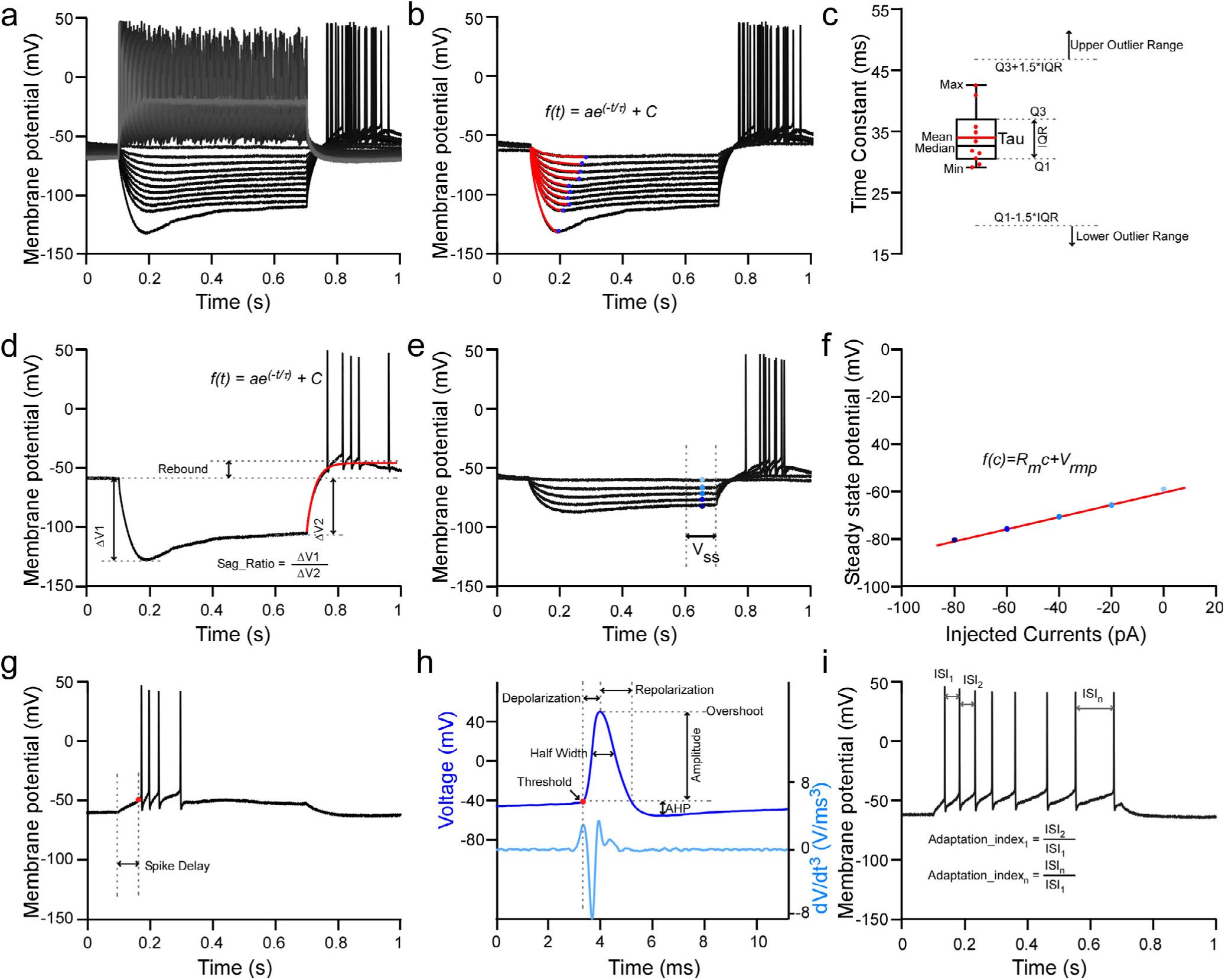
Extraction of the electrophysiological features. Related to Methods. **a**, Membrane potential responses of an example neuron to consecutive step current injections. **b**, A series of membrane potential traces in response to hyperpolarizing current steps, each fitted with an exponential function. The membrane time constant (τ) is calculated for each trace. **c**, Box plot summarizing τ values from one sample neuron, including maximum, minimum, mean, median, first quartile (Q1), third quartile (Q3), and interquartile range (IQR). Outliers (values > Q3 + 1.5*IQR or < Q1 - 1.5*IQR) are removed before calculating the mean τ. **d**, Calculation of the sag ratio and rebound voltage using the first hyperpolarizing trace in the step protocol (inset equations). **e**, Membrane potential responses to the last five negative current injections in the step protocol. Steady-state potentials (V_ss_) are indicated by dashed lines. **f**, Steady-state potentials (V_ss_) plotted against the injected currents and fitted with a linear function. The resting membrane potential (V_rmp_) and input resistance (R_m_) are calculated using the inset equation. **g**, Representative trace showing action potentials (APs) evoked during consecutive step current injections. Spike delay is defined as the time duration between the dashed lines. **h**, (Top) Representative AP trace. (Bottom) Its third derivative (dV/dt^3^). The AP threshold is the membrane potential where the third derivative is maximal (red dot). Definitions of AP depolarization time, AP amplitude, AP repolarization time, AP half-width, and afterhyperpolarization (AHP) amplitude as shown with double-arrow lines and dash lines. **i**, Membrane potential trace in response to 2x rheobase current injection. Initial and last adaptation indices (adaptation_index1 and adaptation_index2) are calculated using the inset equations after determining the interspike interval (ISI) between each AP.

**Extended Data Fig. 15:**
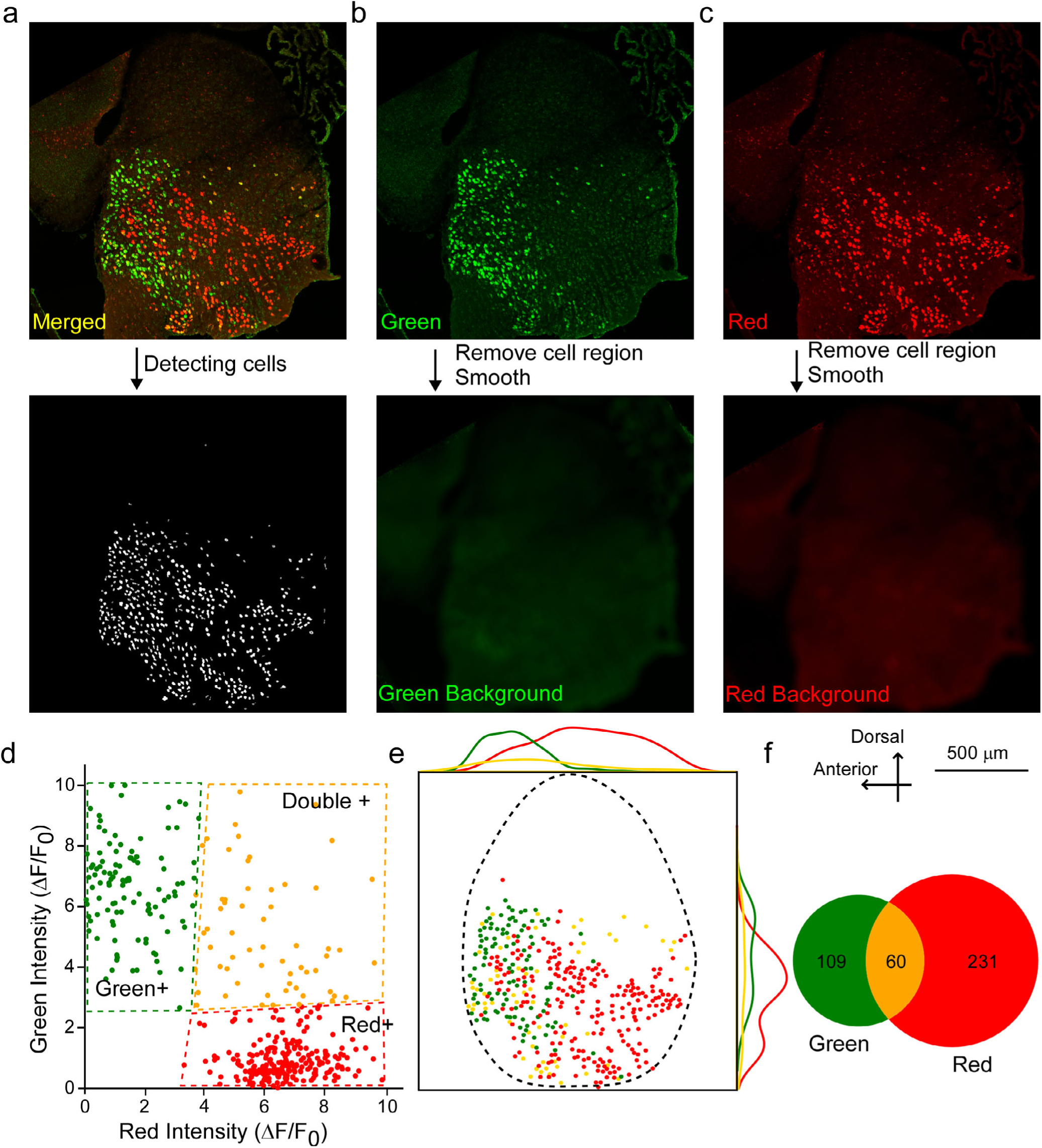
Analysis of dual-color fluorescence in situ hybridization (FISH). Related to Methods. **a**, Top: A representative dual-color FISH staining image (merged). Green channel represents the FITC-labeled RNA probe and red channel represents the DIG-labeled RNA probe. Bottom: each cell with green and/or red signals (F) is detected with a custom-made IDL code, as indicated with a white dot. **b**, Top: A green signal-only image. Bottom: the image indicates the background green signals (Green F_0_), derived by subtracting the signals of detected cells from the images on the top, smoothed by a 60 µm length window. **c**, Top: A red signal-only image. Bottom: the image indicates the background red signals (Red F_0_), derived by subtracting the signals of detected cells from the images on the top, smoothed by a 60 µm length window. **d**, A 2-D scatter plot: the intensity differences (ΔF) between F and F_0_ divided by F_0_ of a detected cell from one channel (ΔF/F_0_) are plotted against ΔF/F_0_ from another channel. Quadrilateral gates are drawn on the plot to determine whether a cell was single-labeled or double-labeled. **e**, Each identified cell (Red^+^, Green^+^, or Double^+^) in their real CN anatomic location, and 2-D density histograms are plotted through the anterior to posterior and dorsal to the ventral axis to show the distribution of Red^+^, Green^+^, and Double^+^ cells along the two axes. **f**, Venn diagrams show the number of Red^+^, Green^+^, and Double^+^ cells in the images shown in (a).

